# Myeloid PINK1 represses mtDNA release and immune signaling that impacts neuronal pathology in patient-derived idiopathic PD models

**DOI:** 10.64898/2026.01.07.694713

**Authors:** Sherilyn Junelle Recinto, Adam MacDonald, Shobina Premachandran, Lin Liu, Armin Bayati, Lilia Rodriguez, Mai Nguyen, Florence Petit, Sriparna Mukherjee, Julien Larmanjat, Alexis Allot, Moein Yaqubi, Peter McPherson, Thomas M. Durcan, Samantha Gruenheid, Louis-Eric Trudeau, Janelle Drouin-Ouellet, Heidi M McBride, Jo Anne Stratton

**Affiliations:** Dept. of Neurology and Neurosurgery, Montreal Neurological Institute-Hospital, McGill University, Montréal, QC, Canada; Aligning Science Across Parkinson’s (ASAP) Collaborative Research Network, Chevy Chase, MD 20815, USA; Dept. of Microbiology and Immunology, McGill University, Montréal, QC, Canada; Faculty of Pharmacy, Université de Montréal, Montréal, QC, Canada; Department of Neuroscience, Faculty of Medicine, Université de Montréal, Montréal, QC, Canada; Dept. of Pharmacology and Physiology, Faculty of Medicine, Université de Montréal, Montréal, QC, Canada; Neural Signaling and Circuitry research group (SNC) and the Center for Interdisciplinary Research on the Brain and Learning (CIRCA)

**Keywords:** Myeloid cells, monocytes, macrophages, PINK1, Parkinson’s Disease

## Abstract

Parkinson’s disease (PD) is a neurodegenerative disorder marked by the development of cardinal motor deficits preceded by a protracted prodromal period of non-motor symptoms often involving the gastrointestinal (GI) tract. There is an emerging consensus that both the peripheral immune system and local neuroinflammation play key roles in the etiology of PD. We previously demonstrated a critical function for the Parkinson’s related proteins PINK1 and Parkin as repressors of the innate to adaptive immune response in cultured cells and mouse models of infection. However, it remained unclear whether these processes were conserved in patient-derived models, and precisely how immune signaling may ultimately drive the death of dopaminergic neurons. Here we show that GI infection of PINK1 knockout (KO) mice triggered acute neurodegeneration which was evident early in the enteric nervous system. Treating wild type enteric or dopaminergic neurons with conditioned medium from immune-stimulated PINK1 KO macrophages was sufficient to promote neuronal disruption in both mouse and human neurons *in vitro*. Within immune-activated macrophages, we reveal that loss of PINK1 led to an enhanced release of mitochondrial DNA (mtDNA) within mitochondrial derived vesicles, leading to the activation of cGAS/STING pathways. These changes were seen in both mouse/human *in vitro* models and in PD patient-derived primary macrophages. Notably, pharmacological modulation using a PINK1 activator with high therapeutic potential attenuated pro-inflammatory profiles elicited by the mtDNA-dependent STING/NF-κB pathway in idiopathic patient-derived macrophages. Ultimately, our study lays the foundation for understanding PINK1-related peripheral macrophage mechanisms in idiopathic PD and provides a target for further development to treat the disease at early stages.

**Graphical abstract:** PINK1-related immune mechanisms of Parkinson’s disease and associations with early neurodegenerative events.

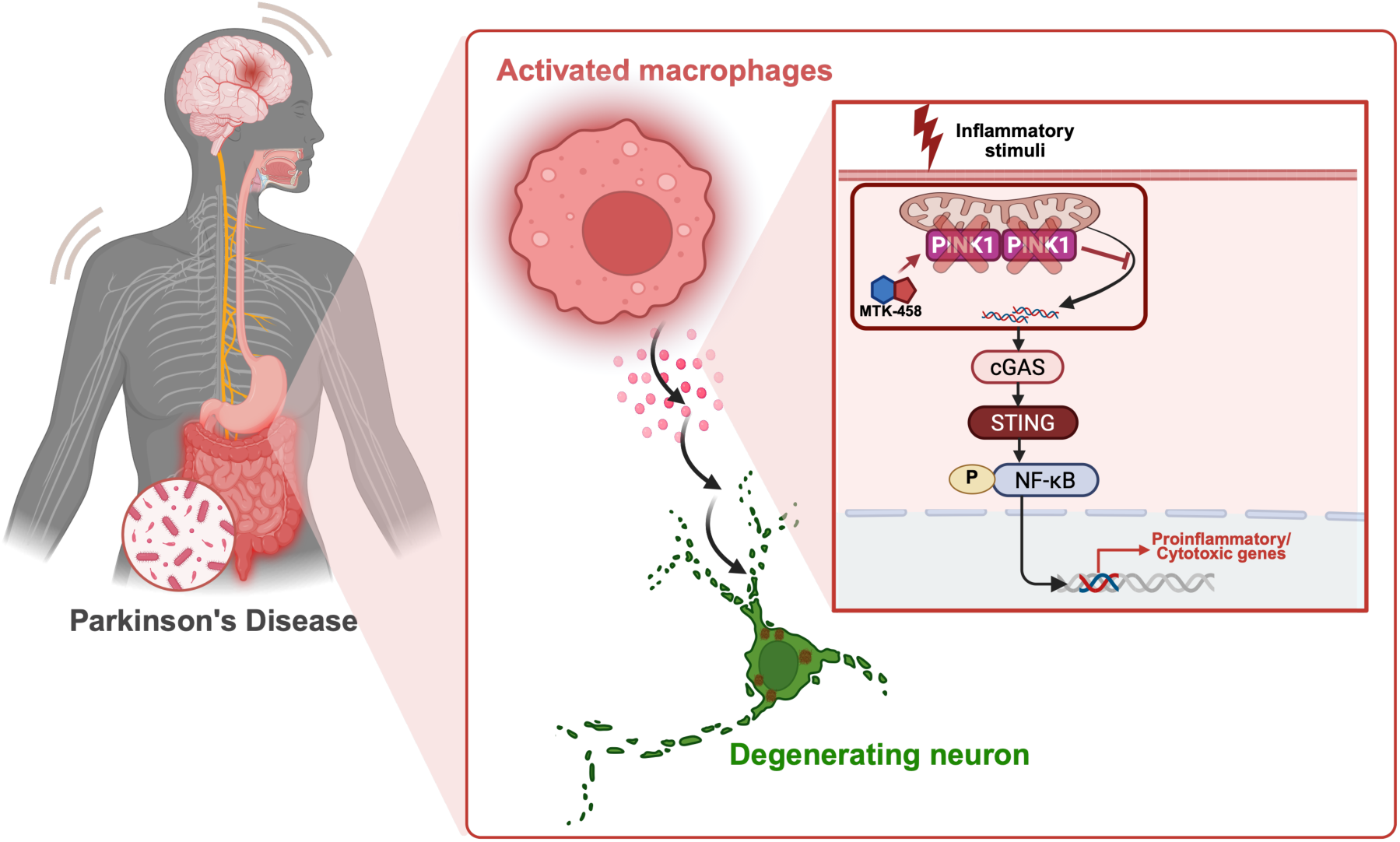

## Introduction

Parkinson’s disease (PD) is a progressive neurodegenerative disorder characterized by the loss of dopaminergic (DA) neurons in the substantia nigra, which drives the appearance of cardinal motor symptoms, such as bradykinesia, tremor, and rigidity. By the time motor symptoms emerge, an appreciable proportion of DA neurons have already been irrevocably lost, and alpha-synuclein (α-syn) pathology is widespread across the central nervous system (CNS), as well as in the periphery, such as in the skin and gastrointestinal (GI) tract. Non-motor symptoms, including constipation, cardiovascular abnormalities, and sleep disturbances, accompany motor deficits, but often precede them by several years or even decades [1–5], These patterns of symptom progression and pathology has culminated in a growing consensus that PD is not a brain-centered disorder albeit a multisystemic disease with significant peripheral involvement, even early in disease. Consistent with Braak’s hypothesis [6, 7], recent evidence supports a substantial subset of PD patients exhibit a “body-first” disease-type, where early autonomic, GI symptoms, as well as rapid eye movement (REM) sleep behavior disorder (RBD) are present years before cardinal motor symptoms [8, 9].

Alongside this evolving understanding of PD’s origin and progression, there is a mounting recognition that the immune system plays a key role in both the initiation and progression of disease. Peripheral immune changes are observed decades before diagnosis; for instance, elevated neutrophil-to-lymphocyte ratios correlate with worse motor outcomes in prodromal PD [10, 11], and immune profiling across the innate and adaptive compartments has revealed broad dysregulation in monocytes, natural killer cells, T cells, and B cells [12–35]. Among these immune cells, monocytes stand out as key players implicated across the entire disease course. Peripheral blood mononuclear cell (PBMC) analyses show that classical monocytes undergo dynamic, stage-specific changes, displaying abnormal metabolic and migratory profiles in early or prodromal PD, and enhanced antigen-presenting and phagocytic capacity in later stages (>5 years post-diagnosis) [15, 18, 23, 36].

Our previous work demonstrated that the PD-related proteins PINK1 and Parkin act to repress the formation of mitochondrial derived vesicles (MDVs) following immune stimulation in culture macrophage cell lines [37]. Upon abrogation of these proteins, mimicking the loss-of-function seen in rare, inherited forms of PD, the mitochondrial proteins within MDVs are processed and presented on major histocompatibility complex I (MHC-I) molecules for presentation to CD8+ T-cells. This suggested that autoimmune mechanisms may contribute to the progression of PD. Consistent with this hypothesis, GI infection of PINK1 knockout (KO) mice led to the development of L-DOPA reversible PD-like motor symptoms [38]. To understand the impact of PINK1 deficiency on early events after infection in the gut, we recently employed single-cell transcriptomics in our GI-infection PINK1 KO mouse model [39]. The data highlighted myeloid cells (particularly monocytes/macrophages) as the earliest and most prominently dysregulated immune population in the periphery, well before motor symptoms appear. These findings positioned peripheral myeloid cells as instrumental in the early progression of PD, emphasizing the need to define the mechanisms driving myeloid dysregulation and its role in instigating neuroinflammation/neurodegeneration from the periphery to the brain. Nevertheless, many questions remained to understand the role of PINK1 within innate immune signaling, and the impact of PINK1 loss on cell-cell interactions that promote neurodegeneration within human and mouse models of infection and inflammation.

Using our established GI-targeted, infection-induced mouse model, we now show that PINK1 KO mice develop early enteric neurodegeneration concurrent with heightened activation of peripheral myeloid cells in the colon. We therefore explored how activated macrophages lacking PINK1 may contribute to the degeneration of neurons early after infection using mouse and human cellular models. Guided by our previous work and computational analysis of the global changes identified in single-cell RNA sequencing (RNAseq), we provide evidence that PINK1 loss leads to the release of mtDNA into the cytosol and activation of cGAS/STING pathways. PINK1’s control of mtDNA release is a critical step to dampen innate immune signaling. Incubation of cultured enteric or dopaminergic neurons with conditioned media from activated PINK1 KO macrophages induced neuronal cell death. Importantly, pharmacological activation of PINK1 in idiopathic patient-derived macrophages repressed mtDNA-dependent STING/NF-κB activation and the pro-inflammatory responses after stimulation. Collectively, these findings provide novel insight into PINK1-dependent immune mechanisms in idiopathic PD and identify a putative targetable pathway in peripheral macrophages. Future work is warranted to dissect the mitochondrial signaling in immune cells and how other PD-associated genes may intersect with inflammatory pathways to promote disease onset and progression.

## Results

### Gastrointestinal infection induces enteric neurodegeneration and pro-inflammatory peripheral myeloid cells in PINK1 KO mice at the early stages of disease

GI dysfunction, including dysphagia, gastroparesis and constipation, are prevalent in ∼60% of PD patients and considered one of the earliest non-motor symptoms [40]. Although the etiology is still unclear, clinical data suggests that these symptoms arise from enteric nervous system (ENS) aberrations [41, 42]. Histological evaluations of the colon in idiopathic PD have reported neuronal loss and atrophy within enteric plexuses [43–45], and a marked reduction in myenteric DA neurons in patients with severe constipation [45]. Increased α-synuclein (α-syn) immunoreactivity has also been described in the gut [46–48], which inversely correlates with neuronal density [43].

Rodent models are thus increasingly being developed to interrogate the mechanisms of PD-related GI dysfunction, including how enteric neuron damage is instigated [49–55]. However, most rely on toxin-based or α-syn-overexpression approaches, which produce acute pathology that poorly reflects the progressive and multifactorial nature of human disease. To address this limitation, our group established a physiologically relevant model that integrates PD-linked genetic predisposition (PINK1 KO) with an environmental insult (GI infection). This paradigm generates a prolonged prodrome with evidenced of intestinal inflammation followed by L-DOPA-reversible motor symptoms and midbrain DA neurodegeneration [38, 39]. That said whether early GI immune perturbations in this context can drive enteric neurodegeneration has not been investigated.

As such, we evaluated whether PINK1 KO mice subjected to GI infection recapitulate early ENS neurodegenerative phenotypes manifesting PD patients (Fig. 1A). Immunohistochemical staining for a pan-neuronal marker β3-tubulin (TUBB3) of colonic tissues at 2-weeks post-infection revealed diminished proportions of TUBB3-expressing neurons in PINK1 KO infected mice compared to uninfected controls, indicating enteric neuron loss (Fig. 1B). To probe potential enteric neuron subpopulations that may be selectively vulnerable to damage in this context, we used dopamine transporter (DAT)-Cre tdTomato reporter mice, coupled with immunolabeling for DA neuron subtypes selectively affected in the midbrain. We focused on G protein-activated inward rectifier potassium channel 2 (GIRK2)-expressing DA neurons, previously identified in the gut [56] and implicated in selective vulnerability in PD [57–59]. In accordance with TUBB3, GIRK2-expressing tdTomato-immunoreactive DA neurons in the ENS were significantly reduced in infected PINK1 KO mice relative to wild type (Fig. 1C). Although overall enteric DA neuron density in PINK1 KO infected mice trended lower without reaching significance (p=0.075 *versus* uninfected controls, data not shown), these findings suggest functional compromise, at the least, within GIRK2-expressing subpopulation.

**Figure 1:**
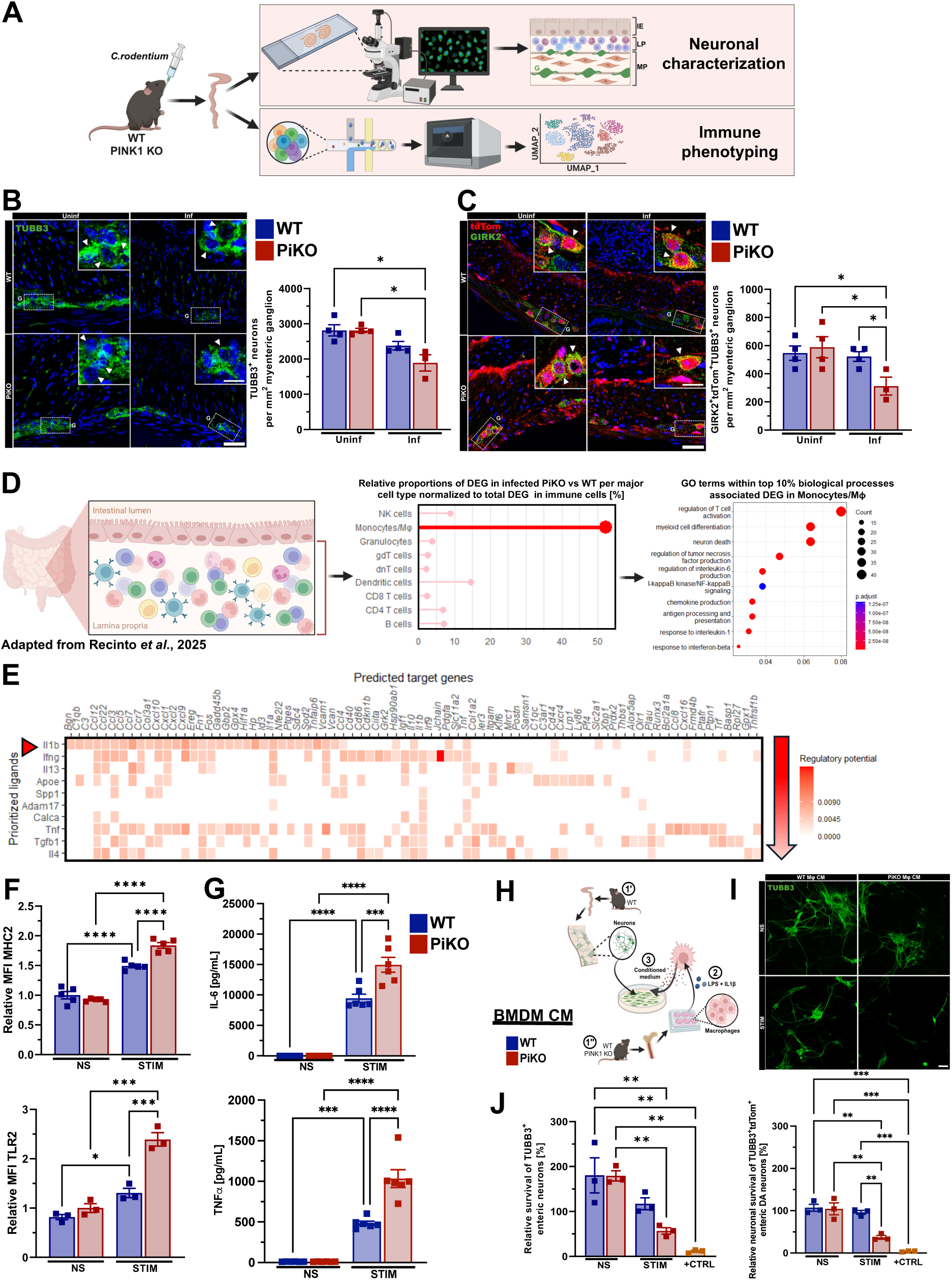
PINK1 KO infected mice show evidence of enteric neuronal pathology and cytotoxic peripheral myeloid cells. **A)** Schematic diagram illustrating gastrointestinal infection with *C. rodentium* via gavage of mice where tdTomato (Ai9) is controlled under the dopamine transporter promoter (referred to as DAT-Cre). Both wild type (WT) and PINK1 KO (PiKO) mice were assessed including uninfected controls. Immunofluorescence was performed for enteric neuronal characterization at 2-w.p.i.; single-cell RNAseq was conducted on lamina propria cells for immunophenotyping at 1-w.p.i. (previously published by Recinto *et al*. [39]) **B, C)** Representative immunofluorescence staining with cross-section view of colonic tissue from DAT-Cre mice. Anti-tubulin beta 3 class III (TUBB3) antibody was used to detect enteric neurons in green (B). Dopaminergic neurons immunoreactive for tdTomato are labelled in red and cells expressing G-protein-regulated inward-rectifier potassium channel 2 (GIRK2) are shown in green (C). Nuclei are stained with Hoechst (blue). Myenteric ganglia are outlined by white dotted box. The inset is a magnified view of enteric neurons and arrow heads indicate immunoreactive cells. Scale bars: 50 μm and 20 μm (for the inset). The density of neurons normalized to ganglionic area (in mm^2^) is quantified across all groups. Two-way ANOVA followed by Tukey’s multiple comparison test was applied. Mean ± SEM, n=3-4 mice per group with 5 ganglia per mouse evaluated. **D)** Re-analysis of single-cell RNAseq of mouse intestinal immune compartment following *C. rodentium* infection taken from Recinto *et al*. [39]. Lollipop plot displays proportions of significant differentially expressed genes (DEG) per major immune cell type previously annotated between infected WT and PiKO mice normalized to total DEGs detected in all immune cells. Wilcoxon Rank Sum test using Bonferroni correction was applied. Enriched Gene Ontological (GO) terms in the monocytes/macrophages (Mϕ) population were assessed. Amongst the top 10%, key GO terms were highlighted in the dot plot. The number of DEG underlying a specified GO term is denoted by the size of the dots while the color scale represents the adj. p-value. The False Discovery Rate (FDR) test was applied. **E)** Heatmap depicting the likelihood of a given ligand from neighboring immune cells could influence the expression of the DEG in infected PiKO monocyte/Mϕ population, termed as regulatory potential. Regulatory potential is determined computationally based on the pearson correlation coefficient between a ligand’s target predictions and the observed transcriptional response. Arrowhead indicates the highest ranked prioritized ligand, likely contributing to transcriptional alterations in infected PiKO monocyte/Mϕ population. **F, G)** Bone marrow-derived Mϕ (BMDM) were cultured from WT and PiKO mice and stimulated for 24 h with lipopolysaccharide (LPS) and mouse recombinant interleukin-1beta (IL-1β). Cell surface expression of major histocompatibility complex 2 (MHC2) and Toll-like receptor 2 (TLR2) were assessed by flow cytometry (F) as well as cell supernatant for secreted amounts of interleukin-6 (IL-6) and tumor necrosis factor alpha (TNFα) by ELISA (G). Shown are relative mean fluorescence intensities (MFI) of cell surface activation markers and relative concentrations of pro-inflammatory cytokines (in pg/mL) across all groups and normalized to the average level in non-stimulated (NS) WT cells. Two-way ANOVA followed by Tukey’s multiple comparison test was applied. Mean ± SEM, n=3-5 mice per group. **H)** Graphical representation of a functional *in vitro* assay to examine the impact of PiKO macrophages on enteric neuronal survival. Primary enteric neurons (1’) and BMDM (1’’) were cultured separately from WT only or both WT and PiKO mice, respectively. Following differentiation of BMDM, cells were stimulated with LPS and mouse recombinant IL-1β for 24h to induce activation (2). Seven days after plating, enteric neurons were treated with conditioned medium (CM) from BMDM for 48h (3). **I)** Representative immunofluorescence staining of mouse primary enteric neurons using a pan-neuronal marker TUBB3. Scale bars: 100 μm. **J)** The proportion of TUBB3^+^ and TUBB3^+^DAT-tdTOM^+^neurons across all groups normalized to the cells receiving BMDM basal medium in the presence of stimuli (to account for possible effects from base medium) was quantified using an automated software for high-throughput imaging (MetaXpress). Staurosporine was used as positive control. Two-way ANOVA followed by Tukey’s multiple comparison test was applied. Mean ± SEM, n=3 independent experiment with each experiment consisted of pooled CM from 3 mice and comprised of triplicates each with 45 sites analyzed.

To determine what may be driving this early neurodegenerative phenotype in the gut, we data mined our published single-cell RNAseq dataset from the colon of PINK1 KO infected mice [39] (Fig. 1A). As illustrated before, we documented a prominent dysregulation of intestinal innate immune populations, particularly monocytes/macrophages, in infected PINK1 KO mice compared to wild type (Fig. 1D). Gene Ontology (GO) analysis of this compartment unveiled enrichment in processes linked to T cell activation, neuron death, myeloid differentiation as well as several inflammatory processes, where we suspect this innate immune signature may mediate the death or damage of enteric neurons

### PINK1 KO in mouse bone marrow-derived macrophages drives a pro-inflammatory response and enteric neuron death in vitro

We next addressed whether PINK1 deficiency in macrophages directly leads to ENS degeneration using an *in vitro* assay, allowing controlled testing of isolated cell responses. To this end, we first interrogated our *in vivo* single-cell RNAseq dataset to identify transcriptional evidence of ligand/receptor pairing shedding light on the relevant cell-cell interactions occurring within the colon of infected PINK1 KO mice using *NicheNet* [60]. To determine ligand activity, *NicheNet* calculates the pearson correlation coefficient between a ligand’s target predictions and the observed transcriptional response (*i.e.,* differentially expressed genes considered significant at adjusted p-values < 0.05 using Wilcoxon rank sum test with Bonferroni correction). This analysis revealed interleukin-1 beta (IL-1β) as the top-ranked ligands, putatively accounting for 60% of PINK1 KO myeloid transcriptional responses, which led us to speculate IL-1β as a major inducer of macrophage activation (Fig 1E). Based on these insights, we selected IL-1β, myeloid differentiation cues essential for bacterial clearance, and lipopolysaccharide (LPS) – found in the outer membrane of Gram-negative bacteria (*e.g., C. rodentium*) – as *in vitro* stimuli to mimic infection-associated signals.

Primary bone marrow-derived macrophages (BMDM) from both PINK1 KO and wild type littermates were thus stimulated under these conditions, and we then used flow cytometry to quantify changes in cell surface marker expression. We indeed demonstrated that PINK1 KO activated BMDM displayed features consistent with intestinal myeloid activation *in vivo*, including elevated cell surface expression of major histocompatibility complex 2 (MHC2) and Toll-like receptor 2 (TLR2) (Fig. 1F). Additionally, supernatants from stimulated PINK1 KO BMDM contained significantly higher levels of pro-inflammatory cytokines interleukin-6 (IL-6) and tumor necrosis factor alpha (TNFα) compared to wild type controls and non-stimulated cells (Fig. 1G), corroborating a heightened pro-inflammatory state.

Next, we interrogated whether the secretome of PINK1 KO macrophages could impair enteric neurons *in vitro*. Longitudinal muscle and myenteric plexuses (LMMP) isolated from colons of adult DAT-tdTomato-expressing wild type mice were dissociated, and primary enteric neurons were matured *in vitro* for 7 days adapted from Smith *et al.* [61]. We subsequently performed exchange experiments using conditioned medium from wild type and PINK1 KO BMDM, following 24-hour stimulation with or without LPS/IL-1β. BMDM conditioned medium were applied to neuronal cultures for 48 hours prior to analysis using immunocytochemistry (Fig. 1H, I). While the medium from wild type activated macrophages did not damage primary enteric neurons, conditioned medium from PINK1-deficient macrophages prompted neurodegeneration, as demonstrated by a loss of TUBB3-expressing primary enteric neurons (in either DAT-tdTomato positive or negative neurons) (Fig. 1J). Treatment of neurons with staurosporine known to promote apoptosis was used as a positive control [62].

### PINK1 KO in human iPSC-derived macrophages drives a pro-inflammatory response and alpha-synuclein dopaminergic neuronal pathology

The most dominant models of PD are associated to common mutations in α-syn, or the expression of α-syn fibrils. We recently correlate the spread of α-syn fibrils to immune signaling, where the generation of pathogenic inclusions of α-syn was triggered only in the presence of pro-inflammatory mediators [63]. Given our evidence that PINK1 plays a key role in innate immune signaling, we were intrigued to decipher whether the secretome of human PINK1 KO macrophages may promote α-syn pathology in dopaminergic neurons. To assess this, we returned to our human model systems consisting of induced pluripotent stem cell-derived macrophages (hiPSC-MDM) and dopaminergic neurons (hiPSC-DA).

We first differentiated PINK1 KO and isogenic control hiPSC into monocyte-derived macrophages (hiPSC-MDM) adapted from the protocol developed by Armitage *et al.* [64]. Briefly, monocytes were generated weekly from CD34+ hematopoietic progenitor cells released from hemogenic endothelium feeder cells. These cells were then re-covered and plated at equal densities for further differentiation into macrophages for an additional 7 days (Fig. 2A). Both genotypes produced comparable yields, viability, and purity (Fig. 2B). Reminiscent of mouse macrophages, we found that PINK1 KO hiPSC-MDM displayed exaggerated responses to LPS/IL-1β stimulation, with enhanced cell surface MHC2 and TLR2 expression (Fig. 2C) and elevated secretion of cytokines and chemokines (Fig. 2D, Supp. Fig. 1). Among the 25 significantly upregulated inflammatory mediators were IL-1β, IL-6 and TNFα, previously implicated in driving PFF-induced pathology in DA neurons [63] and neurotoxicity [65–70].

**Figure 2:**
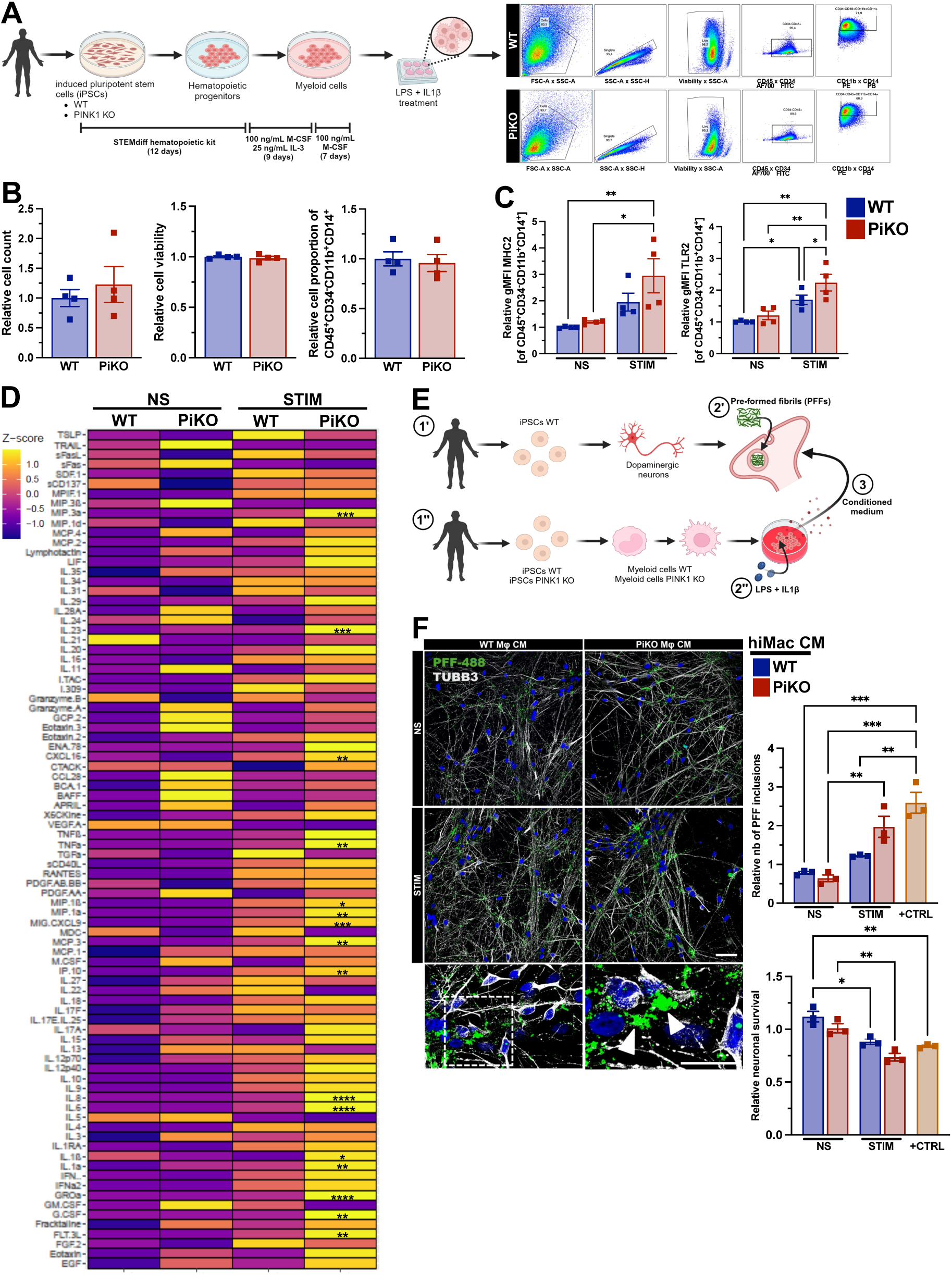
PINK1 loss in human iPSC-derived macrophages induces pro-inflammatory responses and dopaminergic neurodegeneration. **A)** Schematic diagram depicting the differentiation of peripheral myeloid cells derived from WT and PiKO human induced pluripotent stem cells (iPSC) using commercially available growth factors along with recombinant proteins, macrophage colony stimulating factor (M-CSF) and interleukin-3 (IL-3) for 30+ days. Following maturation of monocyte-derived macrophages (MDM), cells were treated with LPS and IL-1β for either 6 or 24 h depending on the downstream biochemical assays performed. Flow cytometric graphs of WT and PiKO human iPSC-derived MDM (hiPSC-MDM) represent gating strategies employed to identify live CD45^+^CD34^−^CD11b^+^CD14^+^ MDM. **B)** Displayed is the quantification of the relative cell count, viability and purity of hiPSC-MDM between genotypes at baseline, determined by flow cytometry. The cell proportions across all groups were normalized to the averaged proportion of WT cells. Two-tailed unpaired t-test was applied. Mean ± SEM, n=4 independent experiment per group. **C)** WT and PiKO hiPSC-MDM were cultured and stimulated for 24 h with LPS and human recombinant IL-1β. Cell surface expression of MHC2 and TLR2 were assessed by flow cytometry. Shown are relative mean fluorescence intensities (MFI) of inflammatory cell surface markers across all groups and normalized to the average MFI of non-stimulated (NS) WT cells. Two-way ANOVA followed by Tukey’s multiple comparison test was applied. Mean ± SEM, n=4 independent experiments per group. **D)** Heatmap illustrating the variation in the z-scores calculated from the mean and standard deviation of the inflammatory cytokines/chemokines measured from the supernatant of stimulated (STIM) WT and PiKO hiPSC-MDM, as well as non-stimulated (NS) control groups taken from three replicates. Two-way ANOVA followed by Tukey’s multiple comparison test was applied to the mean concentrations across all groups. Significant changes in inflammatory mediators between WT and PiKO stimulated hiPSC-MDM were indicated by an asterisk. **E)** Graphical representation of functional *in vitro* assay adapted from Bayati *et al.* [63] to evaluate neuronal α-synuclein pathology in human iPSC-derived dopaminergic neurons (hiPSC-DAn) upon sequential exposure to pre-formed fibrils (PFF) and conditioned media (CM) of activated hiPSC-MDM lacking PINK1. hiPSC-DAn from healthy donors (1’) as well as WT and PiKO hiPSC-MDM (1”) were cultured separately. Following differentiation, at 4 weeks, mature hiPSC-DAn were treated with green fluorescent protein (GFP)-labelled PFF for 2 days then fresh medium was added for an additional 3 days (2’). Concomitantly, MDM were stimulated with LPS and IL-1β for 24 h and CM was collected, including in non-stimulated (NS) control groups (2”). Finally, hiPSC-DAn receive either CM from WT or PiKO MDM of NS or STIM conditions for 2 days, which was then replenished with fresh medium for an additional 7 days before immunostaining was performed (3”). Neuronal treatment with recombinant IFNγ for 1 day after PFF exposure was used as positive control. **F)** Representative immunofluorescence staining of hiPSC-DAn using TUBB3 (white) and detection of PFF inclusions (green) indicative of synuclein pathology. A magnified view of the PFF inclusions found in the dotted white box and denoted by the arrowheads are shown in the inset. Nuclei are stained with DAPI (blue). Scale bars: 100 μm and 10 μm (for the inset). Quantifications of the relative neuronal density and number of PFF inclusions across all groups were conducted as established by Bayati *et al*. [63] and normalized to the neurons receiving MDM basal medium in the presence of stimuli (to account for possible effects from base medium). Two-way ANOVA followed by Tukey’s multiple comparison test was applied. Mean ± SEM, n=3 independent experiment with each experiment consisted of pooled CM from 3 independent MDM cultures and comprised of 5-6 images analyzed.

In accordance with the experimental design in our earlier study [63], we next treated hiPSC-DA neurons with GFP-labelled PFF for 2 days, after which the medium was freshly replenished allowing PFF internalization for 3 consecutive days. We then applied the conditioned medium from the unstimulated or stimulated hiPSC-MDM across genotypes (1:1) to neurons for an additional 2 days. Recombinant human interferon gamma (IFNγ) served as a positive control. Finally, we cultured the cells in fresh medium for an additional 7 days prior to immunostaining (Fig. 2E). We observed that conditioned medium from stimulated PINK1 KO hiPSC-MDM markedly increased α-syn inclusion burden in hiPSC-DA neurons, compared to unstimulated hiPSC-MDM conditions (Fig. 2F). Neuronal density was also reduced under stimulated conditions for both genotypes, suggesting cytotoxic effects of macrophage activation in general, albeit more pronounced with PINK1 deficiency. Of note, α-syn inclusion and neuronal counts were normalized to account for potential hiPSC-MDM base media effects. Collectively, our human *in vitro* studies reinforce that PINK1’s absence in human macrophages creates a neurotoxic environment that exacerbates PD-like neuronal pathology that places PINK1 and α-syn along a common pathway.

### Single-cell RNAseq unearths mitochondrial and immune functions of PINK1 in human iPSC-derived macrophages

To unveil the molecular mechanisms underpinning the pro-inflammatory characteristics of human PINK1 KO peripheral myeloid cells following activation by LPS/IL-1β, we carried out an unbiased single cell transcriptomics approach using 10X Genomics. In brief, we prepared gene expression libraries and employed standard *Seurat* parameters for quality control and integration of datasets (Fig. 3A). Subsequently, we identified cell clusters with high expression of canonical myeloid cell identity markers curated from the literature [20, 71, 72], excluding 8 out of 30 clusters (∼18% of total cells) which represent cells enriched in fibroblast-related genes (Fig. 3B, Supp. Table 1). The myeloid cell clusters were then segregated according to genotype and treatment. As expected, we detected distinct cell cluster separation between stimulated and non-stimulated groups (Fig. 3C). To define responsive populations, we assigned an IL-1β-responsive gene (IRG) score to each cell based on expression of IL-1β target genes identified in Fig. 1E. Cells with high IRG scores (determined as greater than the average score) localized almost exclusively to stimulated groups as expected (Clusters 2, 6, 8, 17, 22, 23, 24, 29) (Fig. 3D, E), with similar proportions in wild type and PINK1 KO cells (Fig. 3F). Differential expression analysis of high IRG-expressing stimulated myeloid cells elucidated substantial transcriptional remodeling, with 1,678 upregulated and 329 downregulated significant genes compared to wild type (adjusted p-values < 0.05 using Wilcoxon rank sum test with Bonferroni correction applied) (Fig. 2G). GO enrichment of upregulated genes pointed to two major categories: mitochondrial processes, particularly energy production and cellular respiration, and innate immune pathways, most prominently NF-κB signaling (Fig. 3H-J). Taken together, PINK1 deficiency alters transcriptional landscape of activated human peripheral myeloid cells possibly through a disrupted crosstalk between mitochondrial and inflammatory signaling that may prime these cells for pathogenic responses.

**Figure 3:**
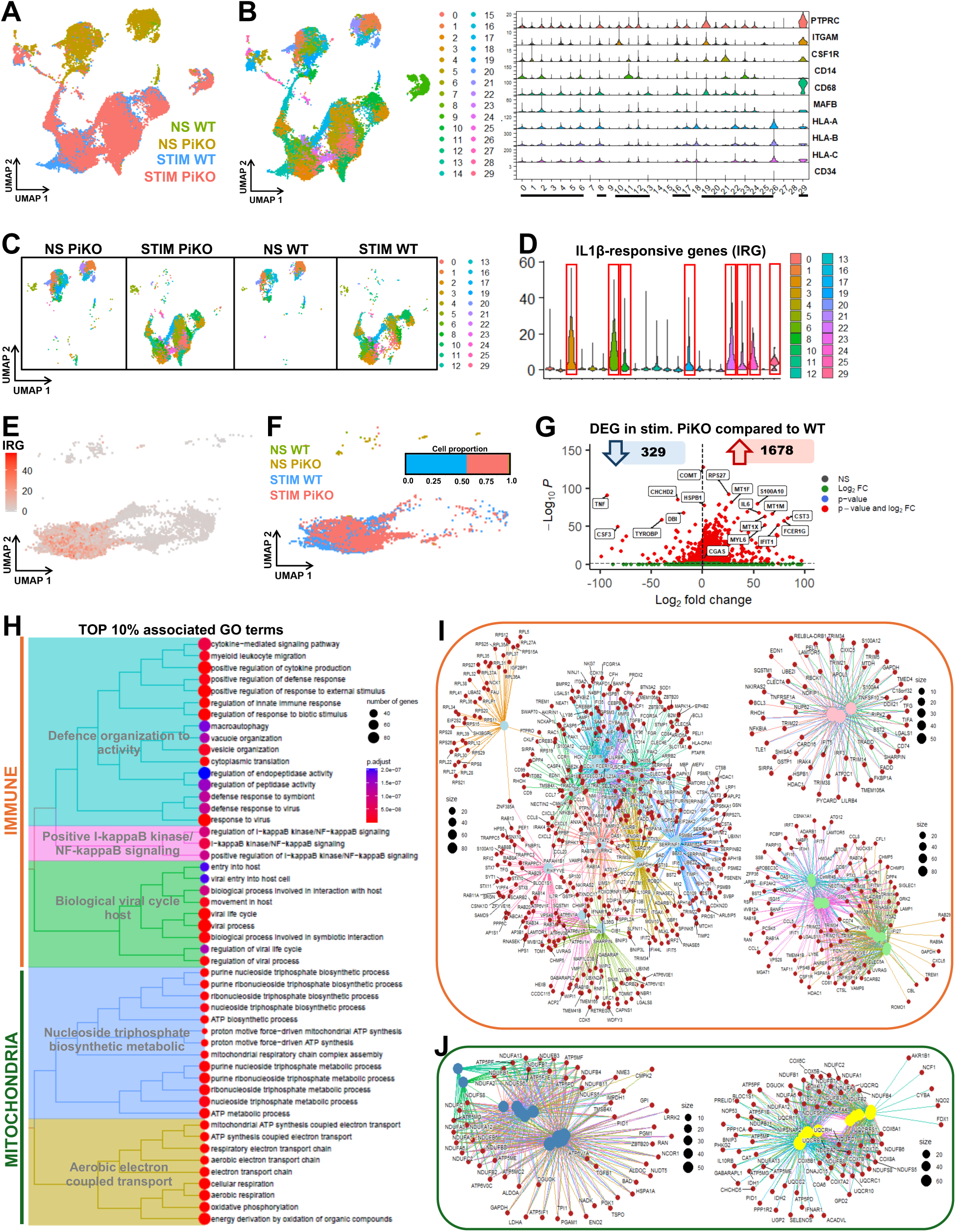
Single-cell RNAseq reveals PINK1-mediated regulation of mitochondrial and immune processes. **A)** Single-cell RNAseq performed using 10X Genomics on WT and PiKO hiPSC-MDM following LPS and IL-1β stimulation, and non-stimulated controls. Integration of all four conditions is displayed in a 2D UMAP plot. **B, C)** Multiple cell clusters across all groups identified by unsupervised hierarchical clustering using standart *Seurat* parameters. Violin plot displays the expression of specific myeloid cell canonical markers in each cell cluster (B). Those with low to no expression of identity genes were excluded from downstream analyses (not underlined) and UMAP plot of bona fide myeloid cell clusters segregated by conditions is rendered (C). **D)** Violin plot depicting the aggregated expression of IL-1β responsive genes (IRG) as determined in Fig. 1E, per cell cluster. This is termed IRG score, with myeloid cell subtypes bearing high IRG score as compared to averaged score (IRG > 2) highlighted in red boxes. **E, F)** UMAP plots illustrating cells with highest IRG score predominantly comprises of WT and PiKO stimulated groups, with equal proportions (blue and red, respectively). **G)** Differentially expressed genes (DEG) between stimulated WT and PiKO within the high IRG-expressing myeloid cell populations displayed in the volcano plot, with few of the top inflammatory-related genes highlighted in boxes. Compared to WT, PiKO stimulated cells have 1,678 significant upregulated and 329 downregulated DEG (red dots). P<0.05 and absolute log_2_foldChange>0.5 was considered significant. Wilcoxon rank sum test with Bonferroni correction was applied. **H)** The top 10% enriched GO terms associated with the significant upregulated genes in PiKO stimulated myeloid cells (determined in G) illustrated as a tree plot, categorizing immune- and mitochondria-related biological processes (orange and green, respectively). The number of DEG underlying a specified GO term is denoted by the size of the dots while the color scale represents the adj. p-value. False Discovery Rate (FDR) test was applied. **I, J)** Cnetplots indicating the genes involved in biological processes pertaining to immune (I) and mitochondrial functions (J). The number of DEG underlying a specified GO term is denoted by the size of the dots.

**Table 1:**
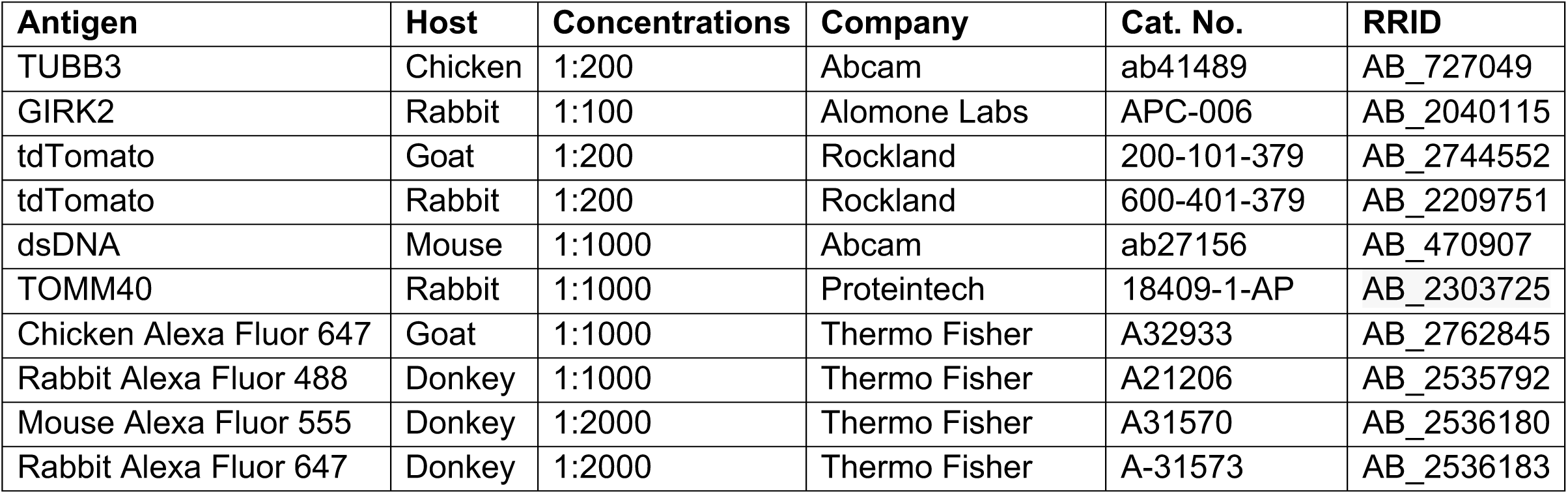
Primary and secondary antibodies for immunostaining.

| Antigen | Host | Concentrations | Company | Cat. No. | RRID |
| --- | --- | --- | --- | --- | --- |
| TUBB3 | Chicken | 1:200 | Abcam | ab41489 | AB_727049 |
| GIRK2 | Rabbit | 1:100 | Alomone Labs | APC-006 | AB_2040115 |
| tdTomato | Goat | 1:200 | Rockland | 200-101-379 | AB_2744552 |
| tdTomato | Rabbit | 1:200 | Rockland | 600-401-379 | AB_2209751 |
| dsDNA | Mouse | 1:1000 | Abcam | ab27156 | AB_470907 |
| TOMM40 | Rabbit | 1:1000 | Proteintech | 18409-1-AP | AB_2303725 |
| Chicken Alexa Fluor 647 | Goat | 1:1000 | Thermo Fisher | A32933 | AB_2762845 |
| Rabbit Alexa Fluor 488 | Donkey | 1:1000 | Thermo Fisher | A21206 | AB_2535792 |
| Mouse Alexa Fluor 555 | Donkey | 1:2000 | Thermo Fisher | A31570 | AB_2536180 |
| Rabbit Alexa Fluor 647 | Donkey | 1:2000 | Thermo Fisher | A-31573 | AB_2536183 |

### Absence of PINK1 in peripheral macrophages triggers mtDNA-dependent STING/NF-kB-driven inflammation following LPS and IL-1β stimulation

The enrichment of mitochondrial and NF-κB-related pathways in our single-cell RNAseq dataset from PINK1 KO hiPSC-MDM (Fig. 3H-J) suggests that PINK1 loss sensitizes macrophages to pro-inflammatory signaling relevant to PD. Given the emerging evidence for PD-related proteins playing roles in the release of mtDNA into the cytosol as a driver of cGAS/STING signaling pathways [73–77], we tested whether a specific STING-related gene signature emerged in PINK1 KO hiPCS-derived MDM. Gulen *et al.* previously used the STING inhibitor H-151 in irradiated human fibroblasts and mixed cortical cells from aging mice to generate a curated list of 108 STING responsive genes (SRG) [78] (Supp. Table 2, Fig. 4A). Analysis of high IRG-expressing myeloid cells from our hiPSC single-cell RNAseq dataset showcased increased SRG scores in PINK1 KO activated myeloid cells, particularly the expression of pro-inflammatory cytokines and chemokines, as well as type I interferons (Fig. 4B).

**Figure 4:**
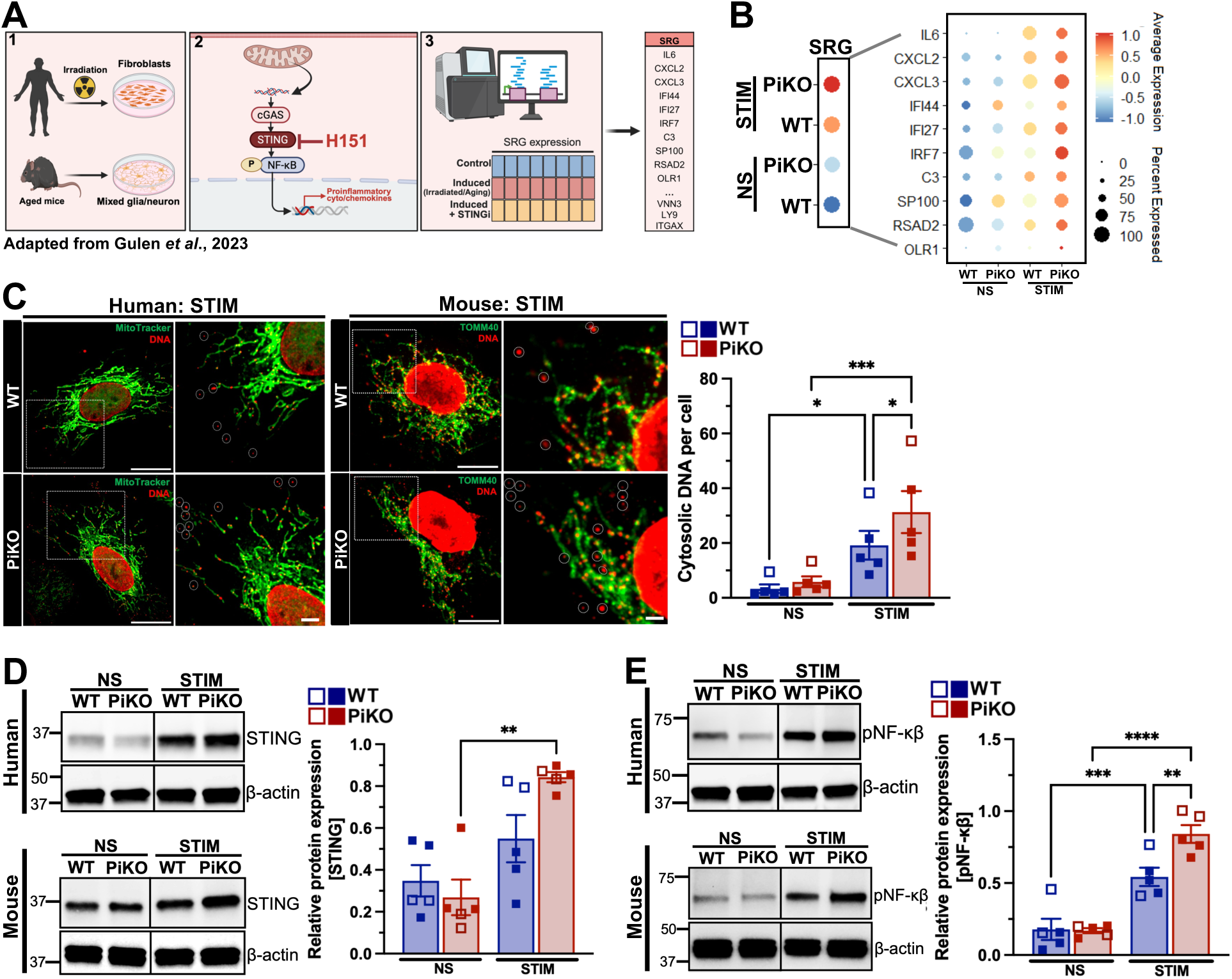
Absence of PINK1 in macrophages promotes mtDNA-dependent STING/NF-κβ driven inflammatory response across species. **A)** Graphical representation of methodology and findings adapted from Gulen *et al.* [78] to identify genes induced by the innate immune pathway involving STING, herein referred to as STING-responsive genes (SRG). Cells taken from either skin of human healthy donors, which are then irradiated to promote aging, or brains from old mice were treated with STING inhibitor (H-151). RNAseq (validated by qPCR) was then performed to interrogate the expression of inflammatory genes induced through the activation of the STING pathway during aging. Using their combined data, a list of SRG comprising of 108 genes was compiled (full list shown in Supp. Table 2). **B)** Dot plot indicating the average expression of SRG across all conditions from the single-cell RNAseq dataset of human iPSC-derived peripheral myeloid cells wherein a larger view showcases key proinflammatory cytokines/chemokines and type I interferons within the SRG panel. The size of the dot determines the proportion of cells expressing the particular gene/gene set while the color scale dictates the normalized average expression. **C)** Representative immunofluorescence staining of macrophages from human (hiPSC-MDM) and mouse (BMDM) origins for DNA (red) and mitochondria (green) using anti-DNA antibody and AlexaFluor 647-labelled MitoTracker or anti-TOMM40 antibody, respectively. Scale bars: 10 μm and 2 μm (inset). The red-positive puncta dissociated from the mitochondria (green signal) was defined as cytosolic DNA in WT and PiKO stimulated cells, as well as non-stimulated controls. Open and closed squares represent data point for human and mouse experiments, respectively. Two-way ANOVA followed by Holm-Sidak’s multiple comparison test was applied. Mean ± SEM, n=5 independent experiment with each experiment comprised of 20-25 images analyzed. **D, E)** Representative immunoblots of human and mouse macrophages stained with anti-STING and anti-phospho NF-κβ (pNF-κβ) antibodies. β-actin was used as loading control. The relative protein expression of STING and pNF-κβ in WT and PiKO stimulated cells, as well as non-stimulated controls, were quantified and normalized to β-actin. Open and closed squares represent data point for human and mouse experiments, respectively. Two-way ANOVA followed by Tukey’s multiple comparison test was applied. Mean ± SEM, n=5 independent experiments.

**Table 2:** Extracellular antibodies for flow cytometry.

| Species | Antigen | Fluorophore | Clone | Concentrations | Company | Cat. No. | RRID |
| --- | --- | --- | --- | --- | --- | --- | --- |
| Mouse | CD11b | PerCP-Cy5.5 | M1/70 | 1:300 | Invitrogen | 45-0112-82 | AB_953558 |
| Mouse | F4/80 | PE-Texas Red | BM8 | 1:300 | BioLegend | 123146 | AB_2564133 |
| Mouse | MHCII | PE-Cy7 | M5/114152 | 1:500 | Invitrogen | 25-5321-82 | AB_10870792 |
| Mouse | TLR2 | BV421 | CB225 | 1:200 | BD | 565908 | AB_2739387 |
| Human | CD34 | FITC | 561 | 1:200 | BioLegend | 343604 | AB_1732005 |
| Human | CD45 | Alexa Fluor 700 | HI30 | 1:200 | BioLegend | 304024 | AB_493760 |
| Human | CD11b | PE | LM2 | 1:200 | BioLegend | 393112 | AB_2750255 |
| Human | CD14 | Pacific Blue | M5E2 | 1:200 | BioLegend | 301815 | AB_2275670 |
| Human | HLA-DR | APC | L243 | 1:200 | BioLegend | 307610 | AB_314688 |
| Human | TLR2 | BV786 | 11G7 | 1:200 | BD | 742771 | AB_2741035 |

We previously documented a key role for PINK1 in suppressing the formation of immune-stimulated mitochondrial derived vesicles (MDV) that led to the presentation of mitochondrial peptides on MHC-I molecules to activate CD8+ T cells. Considering the heightened SRG expression in stimulated PINK1 KO hiPSC-MDM, we assessed the participation of cytosolic mtDNA release in engaging the innate immune STING pathway in our model systems. Indeed, both mouse BMDM and human hiPSC-MDM lacking PINK1 displayed increased cytosolic mtDNA foci following LPS/IL-1β treatments compared to wild type (Fig. 4C). This was accompanied by higher STING expression and NF-κB phosphorylation (Fig. 4D, E), indicative of PINK1 as a repressor of the pro-inflammatory mtDNA-dependent STING/NF-κβ axis. Critically, pharmacological blockade of STING with H-151 attenuated the exacerbated inflammatory response in stimulated PINK1 KO macrophages (Supp. Fig. 2). This inhibition did not reverse the increase in cytosolic mtDNA foci, thus we propose that PINK1 in macrophages acts upstream of STING/NF-κβ signaling by limiting mtDNA leakage into the cytosol.

### Pharmacological activation of PINK1 in patient monocyte-derived macrophages restores dysregulated immune functions in idiopathic PD

Our findings that PINK1 regulates mitochondrial and immune processes in our model systems prompted us to ask whether disrupted PINK1 signaling also contributes to disease etiology in idiopathic PD beyond familial genetic forms. To address if common pathways are observed in PD, we conducted single-cell RNAseq of PBMCs from eight PD (prodromal and clinically established) as well as age- and sex-matched heathy controls (Fig. 5A). Monocytic populations were annotated as CD14+ classical, CD16+ non-classical, transitional pre-monocytes, or hematopoietic stem/progenitor cells (HSCs) progenitors (Fig. 5B). All monocytic subtypes were observed across both conditions, but with higher proportions in PD compared to healthy controls, except for HSCs (Fig. 5C). In line with earlier studies using flow cytometry and bulk RNAseq [14, 17, 20, 27, 79], we discovered extensive regulation in CD14+ monocytes of PD patients (Fig. 5D). GO enrichment analysis of DEGs within this population suggested mitochondrial deficits, with 50% of enriched mitochondria-related processes linked to the *PINK1* gene (Fig. 5F). Despite minimal immune-related GO terms were detected (Fig. 5E), CD14+ monocytes displayed reduced *PINK1* expression concomitant with increased SRG score in PD *versus* controls (Fig. 5G, H), suggesting latent inflammatory potential.

**Figure 5:**
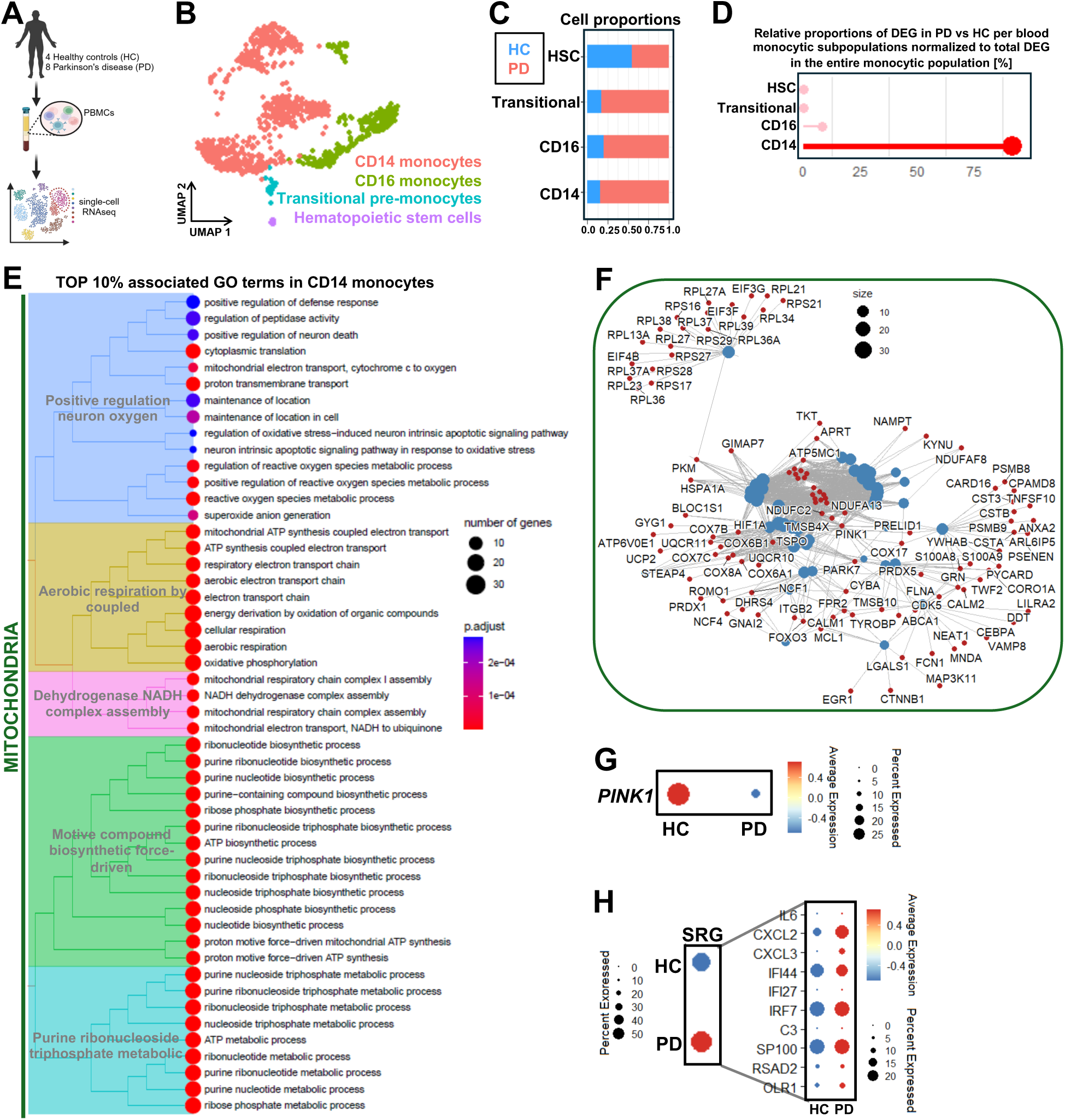
Idiopathic PD patients display PINK1 deficiency and dysregulated PINK1-related functions in blood monocytes. **A)** Graphical representation of single-cell transcriptomic analyses of peripheral blood mononuclear cells (PBMC) from healthy controls (HC) and Parkinson’s disease (PD) patients. **B)** UMAP plot showing monocytic cell populations from PBMC single-cell RNAseq dataset. **C, D)** Cell proportions and significant DEG between HC and PD normalized to total DEGs in the entire monocyte population are displayed per monocytic subtypes identified in the blood. **E)** The Top 10% enriched GO terms associated with the significant DEGs in PD CD14^+^ classical monocytes compared to HC are illustrated as a tree plot, implicating mitochondria-related biological processes. The number of DEG underlying a specified GO term is denoted by the size of the dots while the color scale represents the adj. p-value. False Discovery Rate (FDR) test was applied. **F)** Cnetplot indicating the genes underlying the Top 10% mitochondria-related biological processes. The number of DEG underlying a specified GO term is denoted by the size of the dots. **G, H)** Dot plots displaying the average expression of *PINK1* (G) and SRG (H) between HC and PD within CD14^+^ classical monocytes. A larger view of the SRG panel shows the changes in the expression of key proinflammatory cytokines/chemokines and type I interferons (H). The size of the dot determines the proportion of cells expressing the particular gene/gene set while the color scale dictates the normalized average expression.

We speculated that patient-derived monocytes require a stimulus and enhanced differentiation cues akin to the environment in our GI-targeted, infection-induced model to unleash disease-relevant inflammatory responses conducive to neurodegeneration. As such, monocytes were differentiated into macrophages (MDM) and stimulated with LPS/IL-1β (Fig. 6A). Consistent with observations in our PINK1 KO models, this approach triggered cytosolic mtDNA release and STING/NF-κB activation in PD-derived MDM (Fig. 6B-D). To test therapeutic modulation, we applied the PINK1 agonist, MTK-458, a kinetin analog with better potency and pharmacokinetics currently under evaluation for Phase I clinical trials [80, 81]. Following treatment of stimulated PD-derived MDM with MTK-458, we found a marked decrease in cytosolic mtDNA foci and suppression of STING/NF-κB activation (Fig. 6B-D). Multiparametric protein analyses of the cell supernatant ultimately showed a significant reduction in 32 out of the 92 secreted proteins (35%) assessed following PINK1 activation (Fig. 6E, Supp. Fig. 3). Roughly 40% of these were cytokines/chemokines (highlighted in Fig. 6E) previously reported induced after STING activation [78, 82–89]. Intriguingly, most STING-regulated proteins attenuated by the MTK-458 challenge (∼92%) were inversely affected by the absence of PINK1 in stimulated myeloid cells (Fig. 2C, Supp. Fig. 1), further underscoring the interplay between PINK1 and STING.

**Figure 6:**
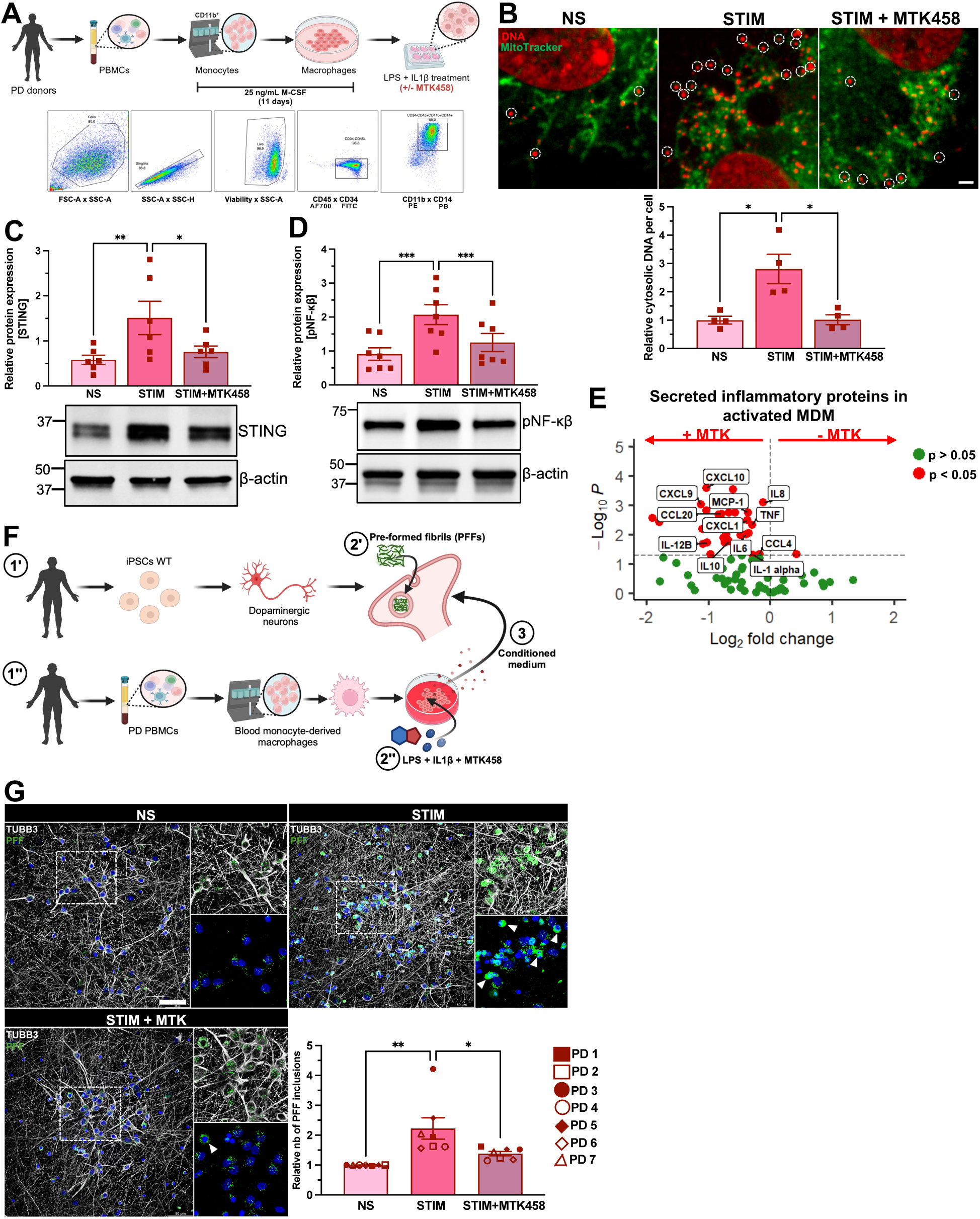
Pharmacological activation of PINK1 attenuates pro-inflammatory responses in idiopathic PD patient-derived macrophages. **A)** Schematic diagram depicting the isolation of blood monocytes from PD donors via CD11b-positive selection magnetic beads and differentiation to macrophages using recombinant M-CSF. Subsequent to maturation, MDM were treated with LPS and IL-1β in combination with ± PINK1 activator (MTK-458) for 6 h. Flow cytometric graph showing gating strategies employed to identify live CD45^+^CD34^−^CD11b^+^CD14^+^ diseased MDM. **B)** Representative immunofluorescence staining of PD blood MDM for DNA (red) and mitochondria (green) using anti-DNA antibody and AlexaFluor 647-labelled MitoTracker, respectively. Scale bar: 2 μm. The red-positive puncta dissociated from the mitochondria (green signal) was defined as cytosolic DNA in PD MDM across the three conditions. Paired one-way ANOVA followed by Tukey’s multiple comparison test was applied. Mean ± SEM, n=4 donors per group with each donor consisted of 20-25 images analyzed. **C, D)** Representative immunoblots of PD blood MDM stained with anti-STING and anti-pNF-κβ antibodies. β -actin was used as loading control. The relative protein expression of STING and pNF-κβ across conditions were quantified and normalized to β-actin. Paired one-way ANOVA followed by Tukey’s multiple comparison test was applied. Mean ± SEM, n=6-7 donors per group. **E)** Volcano plot depicting significant changes in the levels of inflammatory proteins (red dots) detected in the supernatant of stimulated patient-derived MDM in the presence (left) or absence (right) of MTK-458 using a 92-plex assay performed from 3 PD donors. Two-tailed paired t-test was applied. **F)** Graphical representation of functional *in vitro* assay adapted from Bayati *et al.* [63] to evaluate neuronal α-synuclein pathology in human iPSC-derived dopaminergic neurons (hiPSC-DAn) upon sequential exposure to pre-formed fibrils (PFF) and conditioned media (CM) of activated PD patient-derived MDM with or without MTK-458 co-treatment. hiPSC-DAn from healthy donors (1’) as well as blood MDM from PD donors (1”) were cultured separately. Following differentiation, at 4 weeks, mature hiPSC-DAn were treated with green fluorescent protein (GFP)-labelled PFF for 2 days then fresh medium was added for an additional 3 days (2’). Concomitantly, MDM were stimulated with LPS and IL-1β ± MTK-458 for 24 h, and CM was collected, including in non-stimulated (NS) control groups (2”). Finally, hiPSC-DAn receive CM from activated PD MDM either with or without co-treatment of MTK-458, or from non-stimulated control for 2 days, which was then replenished with fresh medium for an additional 7 days before immunostaining was performed (3”). **G)** Representative immunofluorescence staining of hiPSC-DAn using TUBB3 (white) and detection of PFF inclusions (green) indicative of synuclein pathology. A magnified view of the PFF inclusions found in the dotted white box and denoted by the arrowheads are shown in the inset. Nuclei are stained with DAPI (blue). Scale bars: 100 μm and 10 μm (for the inset). Quantifications of the relative number of PFF inclusions across all groups were conducted as established by Bayati *et al*. [63] and normalized to the neurons receiving MDM basal medium in the presence of stimuli (to account for possible effects from base medium). Paired one-way ANOVA followed by Tukey’s multiple comparison test was applied. Mean ± SEM, n=7 PD donors per group with each donor consisted of 7-8 images analyzed.

Finally, to determine whether activation of PD macrophages induces α-syn pathology in DA neurons, we performed exchange medium experiments using aforementioned treatment regime similar to Bayati *et al.* [63]. Conditioned medium from stimulated PD-derived MDM – either untreated or co-treated with MTK-458 –, or non-stimulated was applied to hiPSC-DA neurons following exposure to GFP-labelled α-syn fibrils (Fig. 6F). Consistent with our earlier observations using PINK1 KO hiPSC-MDM (Fig. 2F), conditioned medium from stimulated PD-derived MDM induced a higher burden of α-syn inclusions in DA neurons, compared to non-stimulated controls (Fig. 6G). Strikingly, this effect was markedly reduced when neurons were exposed to conditioned medium from MTK-458-treated, stimulated PD-derived MDM. Ultimately, these findings establish proof-of-concept that modulating PINK1-controlled innate immune signaling can mitigate macrophage-driven neurotoxic signaling and highlight its therapeutic potential in idiopathic PD.

## Discussion

Despite a growing recognition that a dysregulated immune response in PD contributes to disease processes [90–92], the evidence for immune events at early stages of neurodegeneration is less established. This is largely due to limitations in modeling genetic and environmental aspects of immunity in the context of disease initiation. As outlined here, we have put forward both rodent and human models, where early neurodegenerative features are apparent that link PD-related genetics and immune pathways activated through environmental cues. Our data demonstrate that PINK1 acts as a repressor of the mtDNA-dependent STING/NF-κβ pathway in rodent and human macrophages. We reveal that the enhanced innate immune signaling within PINK1 macrophages produces a secretome that drives neurodegeneration directly within the gut niche. The early loss of neurons is consistent with our previous observation that PINK1 KO mice develop constipation early after infection [39], which is a critical feature of prodromal PD. Interestingly, our analysis of DA neurons within the colon further highlighted the selective vulnerability of GIRK2+ neurons, consistent with recent data from primate and human models [57, 58].

In the context of α-syn pathology, the increased innate immune signaling upon loss of PINK1 was seen to promote the accumulation of α-syn aggregates, again converging multiple PD-related proteins along a common immune pathway. Indeed, a key finding is the impact of activated PINK1 KO macrophages on neuronal survival and α-syn pathology. Several inflammatory mediators secreted by activated human (and mouse) PINK1 KO macrophages identified here are also elevated in blood and CSF samples of PD patients based on a large meta-analysis [93]. We did not directly unravel which macrophage-secreted factor might drive α-synuclein pathology; however, in our previous study, we demonstrated that blocking the receptors for IL-1β or TNFα partially reversed the accumulation of neuronal α-syn inclusions after co-stimulation with preformed fibrils (PFF) and conditioned medium from activated hiPSC-derived microglia. Accordingly, administration of these recombinant proteins in this dual-hit model also induces neuronal α-syn pathology [63]. As such, we speculate that similar immune mechanisms might be at play where the joint deleterious effects of PFF and heightened immune signaling resulting from PINK1 loss might impair lysosomal function culminating in the build-up of pathogenic α-syn inclusions. Of note, our model also demonstrated that conditioned media from both activated wild type and PINK1 KO macrophages can trigger neuronal loss. Such observations are in keeping with previous work showing that activated wild type macrophages also secrete neurotoxic factors [65–69], albeit at lower levels. Notably, Sommer *et al.* elucidated a mechanism by which hiPSC-DA neuron underwent increased cell death following addition of pro-inflammatory cytokine, such as interleukin 17 (IL-17) [94]. Since only PINK1 KO macrophage conditioned media induced α-syn inclusions, recapitulating neuronal pathology in PD, PINK1-linked immune signals in this model are of particular relevance to probe immune-driven mechanisms of DA neurodegeneration, whether it be ENS or CNS. We argue that combining PD-related environmental cues, modelled here by PFF and inflammation, in addition to genetic predisposition in immune cells may be sufficient to engender neuronal α-syn pathology in patients at the earliest stages of disease.

Our previous work demonstrated that PINK1 loss promotes mitochondrial antigen presentation, leading to autoimmune mechanisms required for the later onset of PD-like motor symptoms in our mouse model [38]. Our present study underscores PINK1’s implication in repressing innate immune signaling and provides new evidence that mtDNA is packaged within the MDVs we previously reported to carry mitochondrial antigens [37]. In this way, we posit that PINK1 serves as a rheostat to bridge innate and adaptive immune system in both mouse and human models. Its depletion leads to the hyperactivation of these pathways, which has pathological consequences both locally at the site of infection, and in the longer-term wiring of the adaptive immune system.

Mechanistically, the data is consistent with our recent work denoting the importance of mtDNA+ MDV release triggered by inflammatory cues [95]. The release of MDVs was essential for pyroptotic cell death in macrophages, and a genome-wide CRISPR screen identified at least five distinct PD-related proteins that participate along this cell death pathway at either the mitochondria or at the lysosome, where MDVs are delivered. We elucidated that the immune-signaled cleavage of gasdermin E resulted in pore formation in mtDNA-containing lysosomes, leading to content leakage into the cytosol and activation of cGAS and STING. Our work here places PINK1 as an early repressor of mtDNA+ MDV formation, presenting additional PD-related proteins along this critical immune-mediated vesicle transport pathway. Furthermore, we previously reported that the kinase substrate of PINK1, the PD-related ubiquitin ligase Parkin, acted upon sorting nexin 9, culminating to its degradation and abrogation of vesicle formation [37]. Further work is required to detail the precise molecular events at mitochondria that selectively sort mtDNA and antigenic protein cargoes into MDVs, and the interplay of PD-related protein functions at mitochondria and lysosomes. With roles that extend well beyond energy metabolism, mitochondria have emerged as dynamic signaling hubs orchestrating a wide range of cellular processes. We have proposed that mitochondria sit at the nexus of immune signaling and cell death, positioning them as central players in both cellular homeostasis and disease. In our current work, we reveal a distinct, cell-type specific role of PINK1 in rodent and human macrophages, where it modulates mtDNA-dependent STING signaling to drive pro-inflammatory NF-κβ responses. Our findings align with prior reports suggesting enhanced PINK1 function dampens STING-driven inflammation [73–75]; albeit these earlier studies primarily relied on murine models and lacked focus on PD-relevant cell types, limiting their translational relevance.

Finally, our results provide compelling evidence that these pathways extend into sporadic forms of PD since the addition of a PINK1 activator (MTK-458) dampened immune signaling in patient macrophages and improved neurotoxic cell-cell interactions. Multiple earlier reports have underscored the association of familial PD susceptibility genes with immune functions and noted their elevated expression in various immune populations, including myeloid cells [96]. Nonetheless, direct evidence for the involvement of PINK1 in immune-mediated pathological processes potentially underlying idiopathic PD is sparse. Significantly elevated levels of phosphorylated ubiquitin at serine 65 residue (pUb), which is a known PINK1 substrate and indicator of mitochondrial damage, have been detected in postmortem brain tissue and plasma of idiopathic PD patients compared to healthy controls [80, 97]. Correlational analyses additionally denoted worse clinical progression scores with increasing plasma pUb [80]. Others further observed higher secreted PINK1 protein levels in the plasma of idiopathic PD patients [98, 99]. Williams *et al.* revealed that PD patients have more frequent and stronger T cell responses toward PINK1 peptides found in blood [100]. Taken together and given pharmacological activation of PINK1 could rescue the cytotoxic profiles of macrophages from idiopathic PD, it is warranted to further establish how therapeutic targeting of PINK1 can be leveraged to treat the immunological components of PD at early stages of the disease. As the field continues to define the origins and mechanisms of PD pathogenic immune responses, translating such insights into effective, disease-modifying therapies becomes increasingly pivotal. Our study ultimately contributes to that effort by integrating our findings from clinically relevant rodent models and human *in vitro* systems to advance an immune-centric framework for understanding and treating PD.

## Methods

### Study subjects

This study was reviewed and approved by the Montreal Neurological Institute and Hospital (MNI) Review Board. Informed consent was required for participation. Recruitment was conducted from Movement Disorders Clinic at the MNI and a participant was deemed to be living with Parkinson’s disease by our clinicians where metrics used were based on the recently published SynNeurGe research diagnostic criteria [101]. Participation in this study included inclusion criteria of reported age between 49 and 68 years old and a diagnosis of PD or no diagnosis in the case of the healthy control cohort. Blood was collected from 19 individuals with PD and 4 healthy controls identified biologically as male or females.

### Mice

DAT-Ires-Cre (RRID:IMSR_JAX:006660) mice were bred with Ai9 mice (RRID:IMSR_JAX:007909) to induce the expression of tdTomato in DAT-expressing neurons (DATtdTomato). These mice were crossed with PINK1 knockout (PINK1 KO) mice in a mixed B6.129 background (RRID:IMSR**_**JAX:017946) [80]. Adult DAT-Cre-tdTomato wild type and PINK1 KO mice (both sexes) were utilized in the present study. Mice were housed under specific pathogen-free conditions, given water and food ad libitum. All rodent experiments were performed in compliance to the guidelines and conditions specified by the Canadian Council on Animal Care and were approved by the animal care committee of Université de Montréal and McGill University (animal use protocol number MCGL-5009).

### Infection with C. rodentium

As described in Recinto *et al.* [39], mice were infected with chloramphenicol-resistant *C. rodentium* (4 × 10^9^ CFU, ATCC 51459) by oral gavage. To monitor burden of *C. rodentium*, faeces were collected from each mouse, weighed, dissociated in PBS (Thermo Fisher, 10010023), serial-diluted and plated on chloramphenicol-containing MacConkey agar Petri plates. Petri plates were incubated at 37°C overnight to allow colony growth, which were counted the following day. *C. rodentium* colonies were identified by their distinctive morphology on MacConkey agar. Final counts were measured as CFU/g of faeces. Faecal collection occurred on 1-, 2- and 4-weeks-post-infection (w.p.i.) to confirm 10^7^ CFU/g infection rates. Approximately 80% of infected mice achieved this rate and were included in the study.

### Preparation of colonic lamina propria cells

As reported in Recinto *et al.* [39], following mice euthanasia by CO_2_ asphyxiation and cervical dislocation, colons were dissected, and then fecal material was flushed with an 18G needle and adipose tissue was removed. The colon was then cut longitudinally and chopped into sections of 1-2 cm in length. To separate the mucosal epithelium, intestinal tissue sections were incubated in 2 mM EDTA (Thermo Fisher, AM9260G) at 37°C for 15 min followed by vigorous manual shaking three times until cellular suspension appeared dense and cloudy. Remaining intestinal tissue sections (consisting of the lamina propria cells) were then collected and enzymatically dissociated with 2:1 ratio of collagenase IV (Sigma, C5138) to DNAse I (Sigma, 10104159001) at 37°C for 30 min followed by vigorous vortexing. The cellular suspension was then collected through a 70 μm cell strainer. A subsequent similar enzymatic digestion was necessary to completely dissociate intestinal tissue and release the whole lamina propria, and the collected cells were then pooled with the first dissociation. Cells were spun down at 1500 rpm at 4°C for 7 min at the end of each enzymatic digestion, then the pellet was resuspended in PBS supplemented with 1% fetal bovine serum (FBS; Thermo Fisher, A5670801). The percentage of viable cells was assessed by flow cytometry; the average was 60%. Cells were used for cell surface and intracellular staining to acquire cell profile, and cell sorting for qPCR and single-cell RNAseq (whereby only live single cells were collected by FACS). Once separated, the mucosal epithelium was processed independently for single-cell RNAseq. Briefly, the cloudy cellular suspension containing the crypts was further dissociated into single cells by resuspending in TrypLe (Thermo Fisher, 12604013) using a 21G needle followed by 10 min incubation at 37°C. The reaction was neutralized with Advanced DMEM F/12 (Thermo Fisher, 12634010) supplemented with 10% FBS. The cellular suspension was then collected through a 40 μm cell strainer and spun down at 900 rpm at 4°C for 5 min. The pellet was resuspended in PBS supplemented with 1% FBS. The percentage of viable cells was assessed by flow cytometry; the average was 40%. Finally, only live single cells were collected by FACS.

### Preparation of bone marrow-derived macrophages

Sex-paired cohorts of male or female mice at 8 to 12 weeks of age were euthanized and bone marrow progenitor cells were flushed through a 25G needle from both femurs and tibia. Bone marrow cells were further dissociated by passing through an 18G needle and collected through a 70 μm cell strainer. Cellular suspensions were centrifuged at 300 g for 10 min. Supernatants were discarded and cells were resuspended in basal medium comprised of RPMI 1640 (Thermo Fisher, 11875093) supplemented with L-Glutamine (Thermo Fisher, 25030081), Penicillin/Streptomycin (Thermo Fisher, 15140122) and 10% FBS, and cultured in the presence of 20 ng/mL mouse recombinant macrophage colony-stimulating factor (M-CSF) (PeproTech, 315-02). Cells were incubated at 37°C, 5% CO_2_ for 7 days and media changes were completed every 2-3 days M-CSF. Matured macrophages were replated and co-stimulated with 500 ng/mL Ultrapure lipopolysaccharide (LPS) from *E.coli* O111:B4 (Invivogen, tlrl-3pelps) and 50 ng/mL mouse recombinant interleukin-1beta (IL-1β) (PeproTech, 211-11B) with or without 8 μg/mL small molecule STING antagonist H-151 (InvivoGen, inh-h151) for 6 h. For the exchange medium experiment, bone marrow-derived macrophages (BMDM) were treated with inflammatory stimuli for 24 h then cells were washed, and conditioned medium (CM) was collected 24 h later. CM was stored at -80°C for later use. Basal medium with differentiation and inflammatory cocktails alone was used as negative control to eliminate effects of growth medium and added factors.

### Preparation of primary mouse enteric neurons

Adapted from Smith *et al.* [61] and previously described in Recinto *et al.* [56], colons from adult wild type mice were dissected and washed following the aforementioned steps supplemented with 1% antibiotic/antimycotic (Thermo Fisher, 15240062). To pin the colon on a 4% agarose pad containing cold-HBSS (Thermo Fisher, 14175095) with intestinal lumen facing down, the tissue was cut open longitudinally. Under a microscope, the longitudinal muscle and myenteric plexus (LMMP) were then peeled off gently using fine-tip tweezers, which were then cut into smaller 1-2 cm pieces prior to collecting in 15 mL falcon tubes with ice-cold HBSS. Enzymatic dissociation of LMMP sections to release myenteric cells was conducted by submerging the tissue sections in prewarmed 1 mg/mL collagenase IV (Sigma, C5138) diluted in HBSS supplemented with 0.5 mM CaCl_2_ (Sigma, C34006) and 10 mM HEPES (Thermo Fisher, 15630080) for 15 min, at 37°C with gentle shaking every 5 min. The cellular suspension was spun down cells at 300 g, 5 min at 4°C. Then, the cell pellet was gently resuspended in pre-warmed 0.05% Trypsin (Sigma, T4049) diluted in HBSS supplemented with 5 mM EDTA using wide-bore P1000 pipette tips. Cells were then immediately incubated at 37°C for 7 min. The reaction was neutralized by adding ice-cold Advanced DMEM/F12 supplemented with 10% FBS and 1% antibiotic/antimycotic with gentle shaking. Subsequently, cells were spun down at 300 g, 5 min at 4°C and the cell pellet was triturated in complete neuronal media consisting of Neurobasal-A medium (Thermo Fisher, 10888022) supplemented with 2% B-27 (Thermo Fisher, 17504044), N-2 (Thermo Fisher, 17502048), 2 mM L-Glutamine, 10 ng/mL mouse recombinant glial derived neurotrophic factor (GDNF) (Thermo Fisher, 450-10), 1% FBS and 1% antibiotic/antimycotic. Cells were counted and plated on a glass-bottom black 96-well plate at cell density of 2-4 × 10^4^ per well. The plate was pre-coated with 15 µg/mL Poly-L-Ornithine (PLO, Sigma, A-004-C) and 2 µg/mL Laminin (Corning, 354232) at least 24 h prior. Neuronal cultures were maintained for 7 days with daily half-media changes and treated with conditioned medium from mouse BMDM for exchange medium experiments conducted for 48 h prior to immunostaining. Treatment with 1 µM staurosporine (Abcam, ab146588) for 24 h was used as positive control in the exchange medium experiment to induce apoptosis.

### Preparation of human iPSC-derived myeloid cells

The protocol was adapted from Armitage *et al.* [64]. Briefly, wild type and PINK1 KO AIW-002 human iPSC lines [102, 103] were cultured in mTeSR Plus basal medium (STEMCELL Technologies, 100-0276) supplemented with 10 mM ROCK inhibitor (Selleckem, S1049) at 50-70% confluency. At day 0, 20-40 undifferentiated clusters of ∼100 cells each were harvested and differentiated to hematopoietic progenitor cells (HPC) using Medium A from StemDiff Hematopoietic kit (STEMCELL, 05310) for two days which was then replaced with Medium B every two days until day 10 of differentiation. On day 12, CD34^+^ cells were harvested leaving behind a hemogenic endothelium which was then cultured in basal medium consisted of X-VIVO 15 Serum-free Hematopoietic Cell Medium (Lonza, 04-418Q) supplemented with 55 µM β-mercaptoethanol (Sigma, M6250), GlutaMAX (Thermo Fisher, 35050061) and 1% Penicillin/Streptomycin (Thermo Fisher, 15140122) in the presence of human recombinant proteins, 100 ng/mL M-CSF (Peprotech, 300-25) and 25 ng/mL IL-3 (Peprotech, 200-03). The hemogenic endothelium begins producing monocytes over the next 8-9 days and for 30+ weeks. On day 19, freshly harvested monocytes were assessed for purity by flow cytometry and replated in ultra-low attachment plates (Corning, 3471) cultured in basal medium in the presence of 100 ng/mL M-CSF for 10 days with half-media changes every 3-4 days. After maturation, cell viability and purity were evaluated by flow cytometry; on average, 95% viability and 70% CD45^+^CD34^−^CD11b^+^CD14^+^ monocyte-derived macrophages (MDM) were obtained. MDM were subsequently replated for downstream biochemical assays and co-stimulated with 500 ng/mL Ultrapure LPS from *E.coli* O111:B4 (Invivogen, tlrl-3pelps) and 50 ng/mL human recombinant IL-1β (PeproTech, 200-01B) for 6 h. For supernatant analyses and exchange medium experiments, treatment was done for 24 h prior to medium collection. CM was stored at -80°C for later use. Basal medium with differentiation and inflammatory cocktails alone was used as negative control to eliminate effects of growth medium and added factors. For single-cell RNAseq, human iPSC-derived wild type and PINK1 KO peripheral myeloid cells were collected at immature (monocytes) and mature (macrophages) stages following stimulation with LPS and IL-1β for 24 h, along with non-stimulated control groups. All datasets were integrated for downstream bioinformatic analyses. All human iPSC work was done under UdeM ethics approval CERSES-18-004-D.

### Peripheral blood mononuclear cells preparation

Protocols were approved by the McGill University Institutional Review Board of the Faculty of Medicine and Health Sciences (2021-6588). Whole blood from consented donors was collected in heparized tubes (BD Vacutainer, 366480) at the MNI, and processed to isolate peripheral blood mononuclear cells (PBMC) in accordance with the C-BIG repository standard protocol for PBMC isolation (https://www.mcgill.ca/neuro/files/neuro/cbig-02001_v1.0_pbmc_isolation_from_whole_blood_conventional_method.pdf). Briefly, up to 10 mL of whole blood diluted in 10 mL of PBS (Gibco, 311-010-CL), was carefully layered over 15 mL of Ficoll (STEMCELL, 17144003) and was spun down at 700 g for 30 min with brakes off. The buffy coat was collected and washed twice in PBS. PBMC were counted and were either immediately used for cell culture or cryopreserved in 1 mL of 90% human serum (Millipore Sigma, H4522) supplemented with 10% DMSO (ThermoScientific, J66650.AE) at a density of 10-20 million cells. Cryopreserved PBMC was stored long-term in liquid nitrogen.

For *in-vitro* assays, fresh PBMC from individuals with PD were immediately used to purify blood monocytes using CD11b-positive magnetic bead sorting (Miltenyi Biotec, 130-097-142) according to the manufacturer’s instructions. Briefly, PBMC were incubated with CD11b-positive magnetic beads in Miltenyi buffer consisting of PBS supplemented with 6.67% BSA (Gibco, 15260-037) and 4% EDTA (MilliporeAigma, E7889) for 15 mins at 4°C. Following wash, CD11b-labeled PBMC were resuspended in Miltenyi buffer and applied to LS column (Miltenyi Biotec, 30-042-401) equipped with the MidiMACS™ magnet separator (Miltenyi Biotec, 130-042-302) and MACS^®^ MultiStand (Miltenyi Biotec, 130-042-303). To remove any remaining CD11b-negative cells, the column was washed with Miltenyi buffer. CD11b-positive cells were then eluted and spun down at 300 g, 10 min at 4°C. The cell pellet was resuspended in basal media consisting of RPMI 1640 Medium (Gibco, 21870076) supplemented with 1% penicillin/streptomycin, 1% GlutaMAX (Gibco, 35050061) and 10% FBS (Thermo Fisher, A5670801) in the presence of 25 ng/mL human recombinant M-CSF (PeproTech, 300-25). Cells were counted, plated at a density of ∼ 3 × 10^5^ cells per well in a 12-well dish, and incubated at 37 °C in 5% CO₂ for ∼10 days. Half-media changes were performed every three days. Mature patient-derived MDM were co-stimulated with 500 ng/mL Ultrapure LPS from *E.coli* O111:B4 (Invivogen, tlrl-3pelps) and 50 ng/mL human recombinant IL-1β (PeproTech, 200-01B) for 6 h with and without 10 µM MTK-458 (MedChemExpress, HY-152943). For supernatant analyses, supernatant was collected and stored at -80°C for later use.

For single-cell RNAseq, patients comprised of 4 established PD and 4 REM Sleep Behaviour Disorder (RBD) of equal sex as well as age- and sex-matched heathy controls were recruited, and informed consent was obtained for blood collections. Cryopreserved PBMC from these donors were thawed by incubating cells for 90 s in a 37°C water bath and rapidly transferred to LONZA X-VIVO media (Thermo Fisher, BW04-418Q) at 4°C. Cells were pelleted by centrifugation at 300 g for 10 min at 4°C. The pellet was resuspended in dead cell removal microbeads (Miltenyi, 130-090-10) and dead cells were removed according to the manufacturer’s instructions. Cells were counted and resuspended in PBS supplemented with 0.04% BSA (Gibco, 15260-037) at a density of 1000 cells/µL. 30,000 cells collected from each sample were dedicated for single-cell RNAseq.

### Preparation of human iPSC-derived midbrain dopaminergic neurons

The protocol was adapted from Jefri *et al.* [104]. In brief, human iPSC line 3450 from healthy donors (Clinical Biospecimen Imaging and Genetic (C-BIG) repository) were cultured onto plates coated with Matrigel (Corning, 354277) in mTeSR1 medium (STEMCELL, 85857) at ∼80% confluency. Following differentiation, dopaminergic (DA) neural progenitor cells were harvested using StemPro Accutase Cell Dissociation Reagent (Thermo Fisher, A1110501) and plated on coverslips in neural progenitor medium consisting of DMEM/F12 (Thermo Fisher, A4192001) supplemented with N-2 (Thermo Fisher, A4192001) and B-27 (Thermo Fisher, 17504044). To promote maturation, the medium was then replaced with DA neural differentiation medium consisting of BrainPhys neuronal medium (STEMCELL, 05790) supplemented with N2A (STEMCELL, 07152), Neurocult SM1 (STEMCELL, 05711), 20 ng/mL human recombinant brain-derived neurotrophic factor (BDNF) (Sigma, SRP3014), 20 ng/mL GDNF (PeproTech, 450-10), 0.1 µM Compound E (STEMCELL, 73954), 0.5 mM db-cAMP (Sigma, D0260), 200 µM ascorbic acid (Sigma A5960) and 1 µg/mL laminin (Sigma, L2020) for 4 weeks.

As described in Bayati *et al.* for the immune and preformed fibril (PFF) treatment regime [63], 4-week differentiated DA neurons grown on 15-mm coverslips were administered with 1 µg/mL GFP-labelled PFF for 48 h in DA differentiation medium. Neurons were then given fresh media for an additional 72 h. Subsequently, cells were treated with CM from either wild type or PINK1 KO stimulated hiPSC-MDM or non-stimulated controls, or from PD patient-derived MDM (either pre-treated or not with IL-1β/LPS ± MTK-458) and incubated for 48 h. As positive and negative controls, neurons received 0.2 µg/mL IFNγ (Thermo Fisher, 300-02) for 24 h or basal medium in the presence of inflammatory stimuli, respectively. Neurons were then given fresh media for 8 days to allow the formation of PFF-positive inclusions by which time cells were fixed and stained.

### Alpha-synuclein preformed fibrils (PFF) generation

Adapted from the protocol by Feller *et al.* [105], the human wild-type α-synuclein (α-syn) gene was cloned into a pGEX-6P-1 plasmid (University of Dundee MRC Protein Phosphorylation and Ubiquitination Unit, DU30005) and transformed to BL-21(DE3) *E.coli.* (New England Biolabs, C2527H). The protein was over-expressed as GST-tagged α-syn in the presence of the inducer IPTG and was isolated through affinity binding to Glutathione Sepharose® 4B resin (GE Healthcare, 17075601). The GST tag was cleaved off with GST-HRV 3C protease, and α-syn was purified using a GSTrap 4B column (GE Healthcare, 28401748). The purified α-syn was adjusted to a concentration of 5 mg/mL with PBS and sterilized through a 0.22-μm PVDF syringe filter (MilliporeSigma, SLGV033RS). 0.5 mL of purified α-synuclein was shook on a ThermoMixer set at 1000 rpm and 37°C for 5 days to create preformed fibrils (PFF). Then, PFF were placed in a Bioruptor® Pico sonication unit (Diagenode, B01020001) for 40 cycles (30-sec on/30-sec off). Samples (∼20 μL) underwent quality control by conducting thioflavin T assays, dynamic light scattering (using Zetasizer Nano S, Malvern, DLS assay) and electron microscopy imaging.

### Mitochondria/Cytosolic DNA extraction

Subsequent to the activation and pharmacological treatment (if applicable) of BMDM as described above (at a density of 1 × 10^6^ per well of 6-well plate), cells were harvested in 2 mM EDTA diluted in PBS using cell scraper on ice. Cells were pelleted by centrifugation at 400 xg, 5 mins at 4°C. The supernatant was discarded, and the cell pellet was frozen at -80°C until ready for fractionation and DNA extraction.

For mitochondria/cytosol fractionation, commercially available kit was used (Abcam, ab65320). Briefly, frozen cell pellets were resuspended in cold cytosol extraction buffer supplemented with protease inhibitor cocktail (1:500) and 1 mM dithiothreitol (DTT) and incubated on ice for 10 min. Cells are then homogenized gently for ∼40 passes in Cytosolic extraction buffer using ice-cold Dounce tissue grinder (tight). To assess the efficiency of homogenization, ∼2-3 μL of cellular suspension was observed under a microscope to determine whether 70-80% of cells lack a shiny ring around them (indicative of ruptured cell membrane). Homogenates were collected and first spun down at 1000 x g for 10 min at 4°C then the supernatant was transferred to a new tube and pelleted at 10,000 g for 30 min at 4°C. The supernatant was collected, which consisted of the cytosolic fraction while the cell pellet was resuspended and vortexed in mitochondrial extraction buffer supplemented with protease inhibitor cocktail (1:500), serving as the mitochondrial fraction.

DNA from both the cytosolic and mitochondrial fractions were extracted according to the manufacturer’s instructions (Qiagen, 51104). Briefly, proteinase K was added to all samples followed by vigorous mixing. Samples were then lysed and mixed vigorously in Buffer AL then incubated at 56°C for 10 min. DNA was precipitated with 100% ethanol and pulse-vortexed for 15 s before applying to the QIAamp Mini spin column. The samples were spun down then the filtrates were discarded, and the spin columns were washed twice with Buffer AW1 and AW2 respectively. Following washes and centrifugation, DNA was eluted using Buffer AE. DNA concentration was measured using a NanoDrop and was normalized to a final concentration of 2 ng/μL in nuclease-free water.

Cytosolic and mitochondrial fractions were validated by running protein lysates on SDS-PAGE and performing western blotting. Anti-GAPDH and TOMM20 antibodies were used to visualize each subcellular fraction respectively (Table 3). β-actin was used as loading control.

**Table 3:** Primary antibodies for western blot.

| **Antigen** | **Host** | **Clone** | **Concentrations** | **Company** | **Cat. No.** | **RRID** |
| --- | --- | --- | --- | --- | --- | --- |
| pNF-κB | Rabbit | 93H1 | 1:1000 | Cell Signaling | 3033S | AB_331284 |
| STING | Rabbit | D2P2F | 1:1000 | Cell Signaling | 13647S | AB_2943235 |
| β-actin | Rabbit | 13E5 | 1:5000 | Cell Signaling | 4970S | AB_2223172 |
| TOM20 | Rabbit | D8T4N | 1:1000 | Cell Signaling | 42406 | AB_2687663 |
| GAPDH | Rabbit | 14C10 | 1:5000 | Cell Signaling | 2118 | AB_561053 |

### Immunohistochemistry

As described in Recinto *et al.* for immunostaining of mouse colonic tissues [56], colons from wild type and PINK1 KO infected as well as uninfected mice were first cleared fecal materials and adipose tissues following several rounds of washes with PBS. Colons were then cut open longitudinally and coiled around themselves, resembling a *swiss-roll*. Subsequently, the tissue was fixed in 4% paraformaldehyde (PFA) for 24 h then dehydrated with a 30% sucrose solution for an additional 48 h. It was then embedded and cryosectioned serially to obtain coronal sections at 15 μm, then stored at -80°C. Following thawing, tissue sections were permeabilized and blocked in 10% donkey serum (Sigma, D9663) supplemented with 0.5% Triton X-100 (Sigma, 93443) for 2 h. Primary antibody staining was performed overnight at room temperature (RT) using chicken anti-TUBB3, goat anti-tdTomato and rabbit anti-GIRK2 antibodies in blocking solution followed by appropriate secondary antibodies staining for 2 h at RT (Table 1). Hoechst 33258 was used for nuclear staining (1:5000, Thermo Fisher H3569, RRID:AB_2651133). Slides were mounted in Permafluor Mounting media (Thermo Fisher, TA030FM).

### Immunocytochemistry

For enteric neuronal staining in 96-well black glass-bottom plates, cells were fixed in 4% PFA for 15 min at RT then washed thrice with PBS and permeabilized with 0.3% Triton X-100 for 15 min at RT while shaking. Following three washes in PBS every 5 min at RT, cells were blocked in 3% bovine serum albumin (BSA) for 1 h at RT. Subsequently, staining was performed with chicken anti-TUBB3 and rabbit anti-tdTomato antibodies in blocking solution supplemented with 0.3% Triton X-100 overnight at 4°C followed by corresponding secondary antibodies in similar antibody dilution buffer for 1 h at RT while shaking protected from light (Table 1). Cells were washed thrice with PBS every 5 min at RT and nuclei were stained using DAPI (Invitrogen, D1306) for 10 min at RT (1:1000) then washed once.

For DA neuronal staining on coverslips, cells were fixed in 4% PFA for 20 min at RT then blocked and permeabilized in 5% BSA supplemented with 0.05% Triton X-100 for 30 min at RT while shaking protected from light. Cells were subsequently incubated in primary anti-TUBB3 antibody diluted in 5% BSA with 0.01% Triton X-100 for 2 h at RT (Table 1). Subsequent to two washes in PBS every 5 min at RT, cells were stained with appropriate secondary antibody diluted in 5% BSA with 0.01% Triton X-100 for 1 h at RT (Table 1). Cells were washed twice and nuclei were stained with DAPI (Invitrogen, D1306) for 10 min at RT (1:1000). Finally, cells were washed once, and the coverslips were mounted onto microscope slides in mounting media (Dako, S302380-2).

For myeloid cell staining on glass coverslips (Thermo Fisher, 12-545-81), cells were fixed in 5% PFA (Sigma, P6148) for 15 min at 37°C then washed three times in PBS followed by PFA quenching with 50 mM NH_4_Cl (Sigma, A9434) for 10 min at RT. Following three times, cells were permeabilized in 0.1% Triton-X100 for 10 min then washed again three times. Cells were incubated in 5% FBS for 10 min to block then in primary antibodies (diluted in 5% FBS in PBS) for 1 h at RT (Table 1). Either rabbit anti-TOMM40 antibody or Mitotracker dye conjugated with Alexa Fluor 647 (Thermo Fisher, M22426) at 200 nM instead was used to label mitochondria along with mouse anti-dsDNA. Between primary antibodies, three washes in PBS were performed every 5 mins. Subsequently, cells were incubated in the appropriate secondary antibodies for 1 h at RT (Table 1). Cells were then washed twice in PBS, before addition of DAPI (Invitrogen, D1306) for 10 min to stain nuclei. After one further wash in PBS, coverslips were mounted onto microscope slides (VWR, 82003-412) in mounting media (Dako, S302380-2).

### Imaging acquisition and quantification

For colonic tissues, immunostained slices were imaged using a Leica TCS SP8 Confocal microscope (Leica microsystems, RRID:SCR_018169) and quantified with the Las X Leica software. Laser lines corresponding to 405, 488, 555 and 647 nm were used, with excitation and exposure times maintained across samples. Images were taken at 40X magnification with Z-stacks (step size of 0.9). Each ganglion was identified in the myenteric plexus based on morphology as shown in earlier publications [106, 107]. Similar methodology for neuronal density quantification was employed as shown in Recinto *et al.* 2024 [56]. An unbiased assessor defined a nucleus within the ganglia, then scrolled through the Z-stack to ensure that the stains were consistently expressed in and around the cell body surrounding the nuclei. Pan-neuronal marker TUBB3 has clear cytoplasmic signal abutting the nuclei, which made this approach ideal for ensuring accurate quantification. All quantitative analysis at the ganglia was normalized to an area of 900 mm^2^ (calculated as the average size of a ganglion) and 5 ganglia per mouse were analyzed.

For primary mouse enteric neurons, high-content imaging was performed using ImageXpress Micro Confocal (Molecular Devices, RRID:SCR_020294). Laser lines corresponding to 405 and 647 nm were used, with excitation and exposure times maintained across samples for each independent experiment. Each well consisted of 45 sites/images acquired. Images were processed and quantified using the MetaXpress High Content Image acquisition & Analysis software. The application module “Multi Wavelength Cell Scoring” was chosen to obtain an automated quantification of the number or proportion of cells or nuclei in an image. The same parameters of nuclei and cell body sizes as well as intensity threshold per channel were applied across all images in all wells within the same independent experiment.

For human iPSC-derived DA neurons, images were obtained using a Leica TCS SP8 confocal microscope (Leica microsystems, RRID:SCR_018169). Laser lines corresponding to 405, 488 and 647 nm were used, with excitation and exposure times maintained across samples for each independent experiment. Images were processed using FIJI Image J software. For quantification of cell count and inclusions, an unbiased assessor loaded single-channel images into ImageJ and manually counted the number of either nucleus or puncta/particles with a diameter of greater than 2 μm (anything less are considered lysosomal PFF). For cell count, the nuclear staining channel (DAPI) was used, and, for inclusions, the PFF channel (GFP) was used. To ensure PFF signal is observed within neurons, cells were stained with anti-TUBB3 antibody (Table 1). The same threshold was used for all the conditions within each experiment.

For peripheral myeloid cells, images were acquired using an Olympus IX83 confocal microscope containing a spinning disk system (Andor/Yokogawa CSU-X) and Andor Neo sCMOS camera and MetaMorph software. Laser lines corresponding to 405, 555 and 647 nm were used, with excitation and exposure times maintained across samples for each independent experiment. Z-stacks of 0.2 μm per stack were used. Images were processed using FIJI Image J software. For counting the number of cytosolic mtDNA foci, a maximum intensity projection was generated, before automated image analysis macros enabled unbiased quantification as employed by Nguyen *et al.* [108]. First, a mask of the nucleus (DAPI signal), mitochondria (TOMM40 or MitoTracker dye), and mtDNA (DNA signal) were generated. mtDNA foci outside of mitochondria were calculated by subtracting the nuclear and mitochondrial masks from the mtDNA mask, leaving only non-nuclear and non-mitochondrial foci. Cell perimeter was drawn by hand for each cell, and foci within the perimeter were counted.

### Flow cytometry

Peripheral myeloid cells were collected in a 96-well round bottom plates and initially stained for viability using eFluor506 (Invitrogen, 65-0866-14) for 20 min on ice away from light. Prior to washes with FACS buffer (PBS supplemented with 2% FBS and 1 mM EDTA), cells were spun down at 400 g at 4°C for 5 min. For human cells, Fc receptors were blocked with Human TruStain FcX (BioLegend, 422301) for 15 min at RT while mouse cells were blocked with anti-mouse CD16/CD32 (Invitrogen, 14-01061-86) in combination with extracellular staining cocktail for 30 min on ice. Cells were incubated in extracellular fluorescently conjugated antibodies diluted in FACS buffer for 30 mins on ice, in the dark. Table 2 displays the primary antibodies used for both mouse and human cells. After two washes, cells were resuspended in FACS buffer and acquired using Attune NxT flow cytometer (Thermo Fisher, RRID:SCR_019590). Appropriate controls were applied such as single-stained using UltraComp eBeads (Invitrogen, 01-2222-42) and fluorescence minus one (FMO) using corresponding cells.

### Multiplex secreted protein analysis

Supernatant from human iPSC-derived wild type and PINK1 KO monocyte-derived macrophages (MDM) following activation and non-stimulated controls were collected and spun down to get rid of dead cells/debris, which were then sent to Eve Technologies for quantification of inflammatory cytokines/chemokines (Human Cytokine/Chemokine 96-Plex Discovery Assay Array). Three replicates per experimental group were evaluated.

Supernatant from blood PD patient-derived MDM following activation in the presence or absence of MTK-458 were collected and spun down to get rid of dead cells/debris, which were then sent to Olink Technology for quantification of inflammatory cytokines/chemokines (Human 92-Plex Inflammation panel). Three independent donors per experimental group were evaluated.

### SDS-PAGE and Western Blot

Cells were lysed in RIPA buffer (50 mM Tris-HCl, pH 7.4, 150 mM NaCl, 1% NP-40, 0.1% SDS) supplemented with EDTA-containing protease and phosphatase inhibitors (Thermo Fisher, A32959) on ice. Total cell lysates were collected by scraping and were vigorously vortexed for 15 s every 10 min for 30 min. Solubilized proteins are then spun down at maximum speed for 30 min and supernatant was collected. Protein concentration was quantified and normalized using BCA Pierce Quantification Kit (Thermo Fisher, 23225). Laemmli sample buffer (BioRad, 1610747) supplemented with β-mercaptoethanol was added and protein was boiled at 95°C for 5 min. Equal protein amounts were loaded and separated on 4-15% Mini-PROTEAN TGX Precast gels (BioRad, 4561084) and finally transferred to 0.2 μm nitrocellulose membranes using a Trans-Blot turbo system (BioRad, 1704270). Membranes were blocked in 5% BSA in Tris-buffered saline (TBS) for 30 min at RT and incubated with the primary antibodies diluted in 5% BSA TBS overnight at 4°C while shaking (Table 3). On the subsequent day, membranes were washed thrice in TBS with 0.05% Tween20 (TBS-T), incubated with the appropriate secondary antibody (diluted in 5% BSA in TBS-T) for 1 h at RT and finally washed thrice in TBS-T. The membrane was then developed in Clarity Max Western ECL Substrate (BioRad, 1705062) and visualized using the ChemiDoc imaging system (BioRad, RRID:SCR_019037). Images were processed and quantified using FIJI Image J software.

### Quantitative reverse transcription PCR

RNA was extracted using the RNeasy mini kit (Qiagen, 74104) as instructed by the manufacturer. High-Capacity cDNA Reverse Transcription Kit (Thermo Fisher, 4368814) was then used for reverse transcription to generate cDNA. Either cDNA or DNA (from subcellular fractions) was used for real-time quantitative PCR (qPCR) via TaqMan assays on a QuantStudio^TM^ 5 real-time PCR system (Thermo Fisher). The 2^−βΔCt^ method was used to analyse the data using *β-ACTIN* as control. Table 4 illustrates the probes used.

**Table 4:** TaqMan probes for qPCR.

| **Species** | **Probes** | **Assay ID** | **Company** | **Cat. No.** |
| --- | --- | --- | --- | --- |
| Human | MT-ND1 | Hs02596873_s1 | Thermo Fisher | 4331182 |
| Human | ACTB | Hs01060665_g1 | Thermo Fisher | 4331182 |
| Mouse | Il6 | Mm01211445_m1 | Thermo Fisher | 4331182 |
| Mouse | Ifnb1 | Mm00439552_s1 | Thermo Fisher | 4331182 |
| Mouse | Actb | Mm02619580_g1 | Thermo Fisher | 4331182 |

### Single-cell library preparation and analysis

Live cells were prepared for single-cell RNAseq at final concentration of 1000 cells/μL. All cells were processed according to 10X Genomics Chromium Single Cell 3’ Reagent Guidelines (https://support.10xgenomics.com/single-cell-gene-expression). Briefly, cells were partitioned into nanoliter-scale Gel Bead-In-EMulsions (GEMs) using 10X GemCode Technology. Primers containing (i) an Illumina R1 sequence, (ii) a 16 bp 10x barcode, (iii) a 10 bp Unique Molecular Identifier (UMI) and (iv) a poly-dT primer sequence were incubated with partitioned cells resulting in barcoded, full-length cDNA from poly-adenylated mRNA. Silane magnetic beads were used to remove leftover biochemical reagents/primers, then cDNA was amplified by PCR. Enzymatic fragmentation and size selection was used to optimize cDNA amplicon size prior to library construction. R1 (read 1 primer sequence) were added during GEM incubation, whereas P5, P7, a sample index (i7), and R2 (read 2 primer sequence) were added during library construction via end repair, A-tailing, adaptor ligation and PCR. Quality control and quantification was performed using an Agilent Bioanalyzer High Sensitivity chip. Sequencing was performed using NovaSeq 6000 S4 PE 100bp, which resulted in a read depth of ∼20 000 reads/cell. Reads were processed using the 10X Genomics Cell Ranger Single Cell 2.0.0 pipeline (RRID:SCR_017344, https://support.10xgenomics.com/single-cell-gene-expression/software/pipelines/latest/what-is-cell-ranger) with default and recommended parameters, as previously described by Zheng *et al.,* 2017 [109]. FASTQs generated from sequencing output were aligned to either the mouse GRCm38 or human GRCh38 reference genomes using the STAR algorithm 2.7.3a (RRID:SCR_004463, http://code.google.com/p/rna-star/). Next, gene-barcode matrices were generated for each individual sample by counting unique molecular identifiers (UMIs) and filtering non-cell associated barcodes. This output was then imported into the Seurat v4.9.9.9039 R toolkit v.4.2.1 (for quality control and downstream analysis of our single-cell RNA-seq experiment [110]. All functions were run with default parameters, unless specified otherwise. Low quality cells (<200 genes/cell and >5 % of mitochondrial genes) were excluded from the overall experiment. Gene expression was log normalized to a scale factor of 10 000.

### Statistical analysis

Data were analyzed using GraphPad Prism 9.0. Statistical comparisons were performed as noted in the Figure legends using either Student’s *t*-test when only two conditions were being compared, one-way ANOVA for multiple conditions or two-way ANOVA when comparing multiple independent variables and multiple dependent variables. When performing multiple comparisons, p-values were adjusted using post-hoc multiple comparison tests indicated in the Figure legends. Mean and standard error of the mean (SEM) of biological replicates (*n*) are plotted in all graphs and shown in the Figure legends. P-value < 0.05 was considered statistically significant.

### Illustrations

Graphical abstract and representation of treatments and models were created using BioRender.

## Acknowledgment

We would like to thank Christina Gavino and Vanessa Omana for the technical support in inoculating mice and bacterial analyses as well as preparation of gene expression libraries, respectively. We would also like to thank all members of the Desjardins ASAP team for the insightful feedback throughout the course of this study. The study was funded by the joint efforts of The Michael J. Fox Foundation for Parkinson’s Research (MJFF) and the Aligning Science Across Parkinson’s (ASAP) initiative. MJFF administers the grant ASAP 000525 on behalf of ASAP and itself. SJR received a CIHR-Vanier Canada Graduate Scholarship. The Trudeau lab also received support from the Canadian Institutes of Health Research (CIHR) (grant PJT-165928).

## Author Contributions

SJR led all experiments including cellular, biochemical, imaging and bioinformatics work. AM, SP, LL supported primary rodent and human monocyte-derived macrophages cultures, immunohistochemistry and imaging. AM generated the PBMC single-cell RNAseq dataset. AB maintained hiPSC dopaminergic neuronal cultures and supported imaging and analysis of pre-formed fibril experiments. LR and FP optimized and generated hiPSC myeloid cell cultures. MN supported imaging and analysis of mitochondrial DNA. SM bred and maintained animal colony. AA created the website allowing access to and user-friendly analysis of the single-cell RNAseq data. MY ensured open science access to the single-cell datasets. PM and TMD provided technical support and materials. SJR and JAS conceived the project, prepared figures, and drafted the manuscript. JAS, HM, LET, JDO and SG provided funding, supervised the work, and revised the final manuscript. All authors contributed to the article and approved submitted version.

## Data availability

All protocols are available at https://www.protocols.io/view/protocols-for-34-myeloid-pink1-represses-mtdna-rel-gzdrbx257. Raw and processed single-cell data is deposited in a public repository at GEO (GSE313883). While source code and raw tabular data is available at https://doi.org/10.5281/zenodo.15213551. Single-cell RNAseq data is accessible through a user-interface website available for further analyses at https://singlocell.openscience.mcgill.ca/display?dataset=RNA_Hu_CulturedCell_PiKO_Stimulation_2025.

---

**Supplementary Figure 1:**
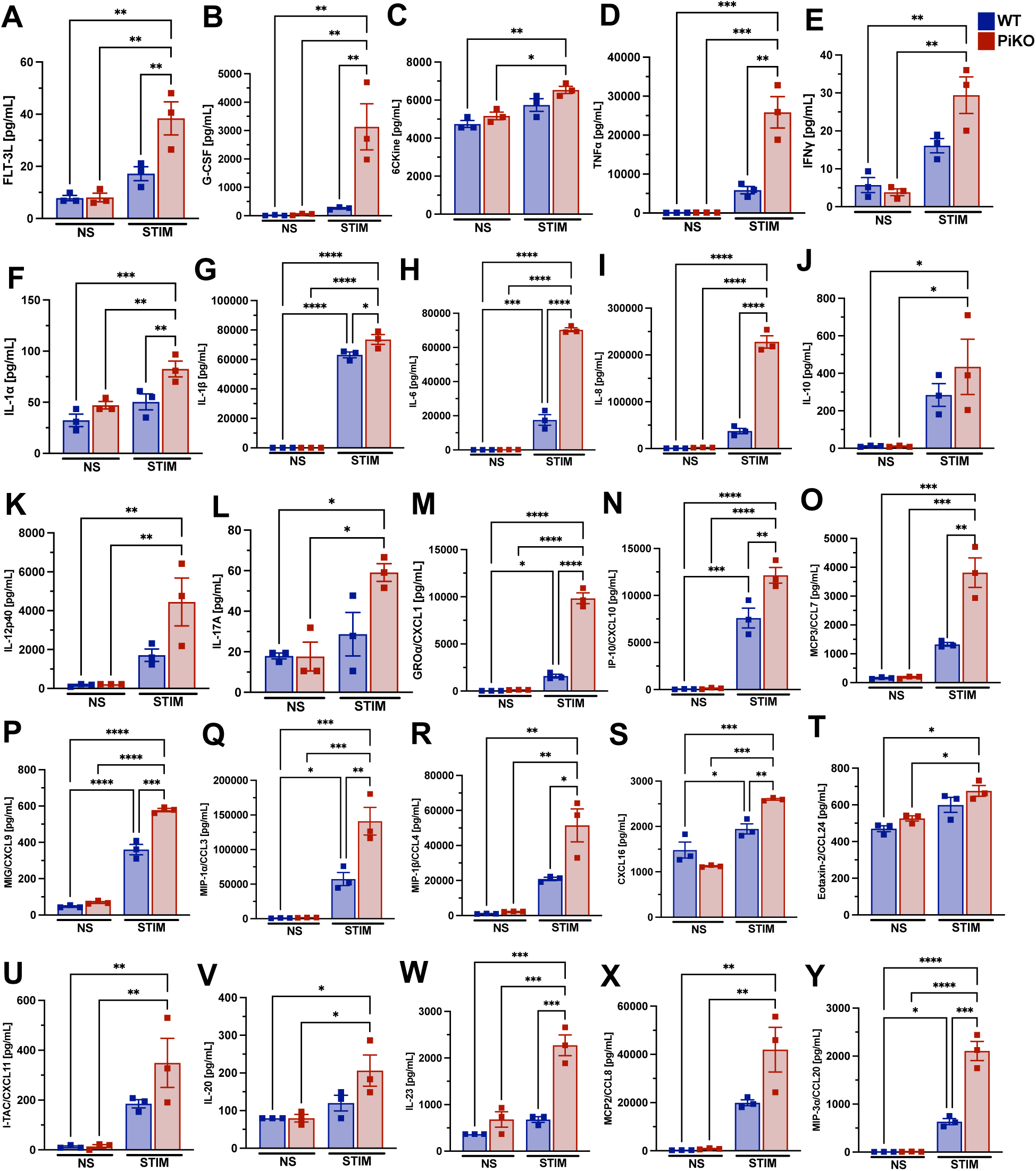
PINK1 KO macrophages secrete higher levels of proinflammatory cytokines following stimulation with lipopolysaccharide and interleukin-1β. Using a 96-plex ELISA, shown are individual inflammatory cyto/chemokine that were significantly higher in the supernatant of stimulated (STIM) PiKO hiPSC-MDM compared to WT and/or non-stimulated (NS) controls. Two-way ANOVA followed by Tukey’s multiple comparison test was applied. Mean ± SEM, n=3 independent experiments.

**Supplementary Figure 2:**
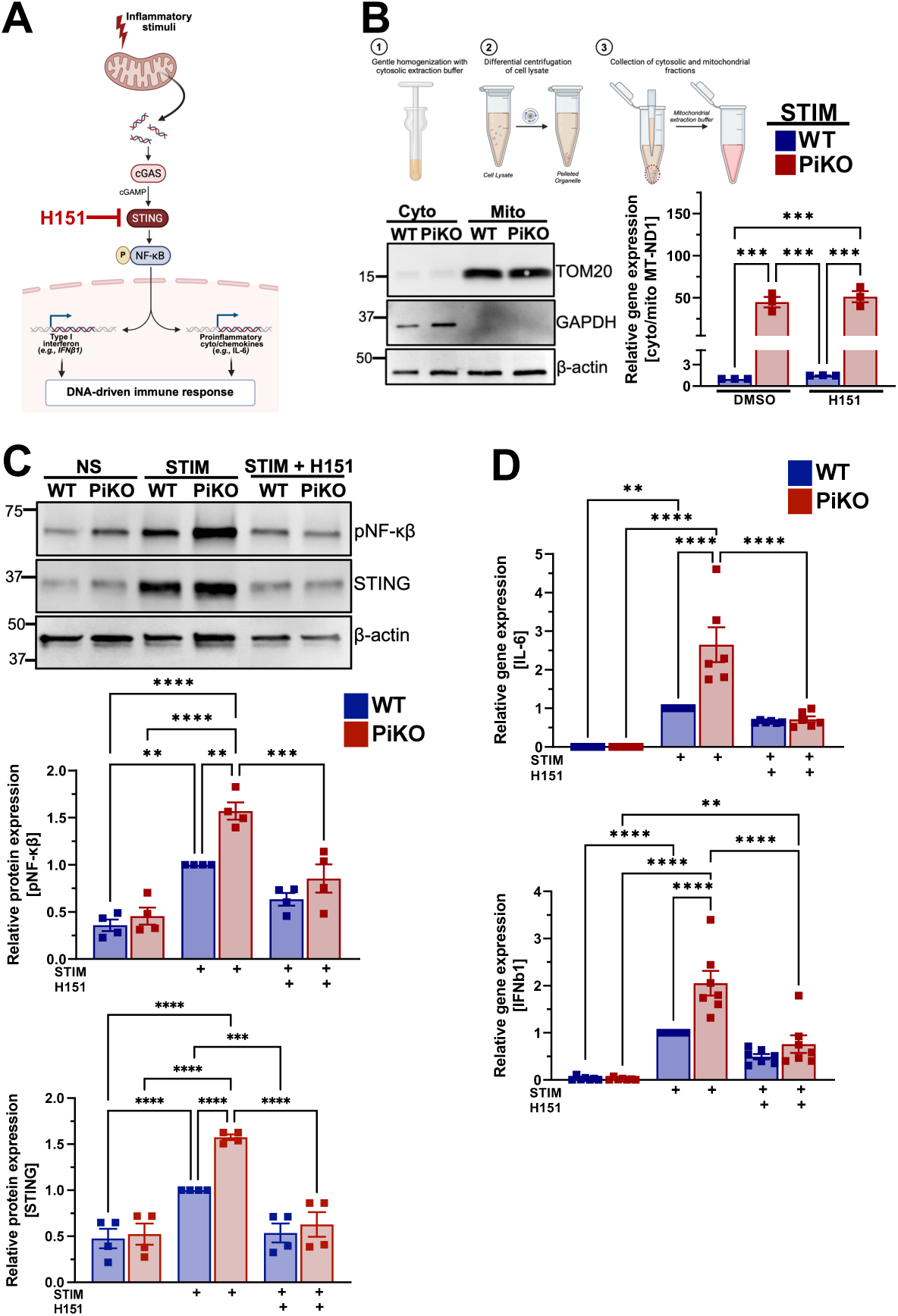
STING inhibition mitigated mtDNA-dependent STING/NF-κβ activation triggered by PINK1 deficiency. **A)** Graphical representation of mtDNA-dependent STING/NF-κβ signaling pathway resulting in inflammation, underscoring H-151 as a STING inhibitor. **B)** Schematic diagram to describe cell fractionation using commercially available buffers and differential centrifugation. Shown are representative immunoblots of cytosolic (cyto) and mitochondrial (mito) fractions isolated from stimulated WT and PiKO BMDM indicating purity of subcellular fractions according to the expression of mitochondrial protein TOMM20 and cytosolic protein GAPDH. β-actin was used as loading control. Both fractions from stimulated BMDM of WT and PiKO mice were assessed for relative gene expression of mtDNA (MT-ND1). Relative ratios of cyto-to-mito MT-ND1 (cyto/mito) in stimulated PiKO with and without H-151 were normalized to corresponding WT groups. Two-way ANOVA followed by Tukey’s multiple comparison test was applied. Mean ± SEM, n=3 mice per group. **C)** Representative immunoblots of WT and PiKO BMDM stained with anti-pNF-κβ and anti-STING antibodies. β-actin was used as loading control. The relative protein expression of pNF-κβ and STING in stimulated cells with or without H-151, as well as non-stimulated controls, were quantified and normalized to β-actin. Two-way ANOVA followed by Tukey’s multiple comparison test was applied. Mean ± SEM, n=4 mice per group. **D)** Quantification of inflammatory target genes downstream of STING innate immune pathway (IL6 and IFNB1) by qPCR across all conditions. Twoway ANOVA followed by Tukey’s multiple comparison test was applied. Mean ± SEM, n=6-7 mice per group.

**Supplementary Figure 3:**
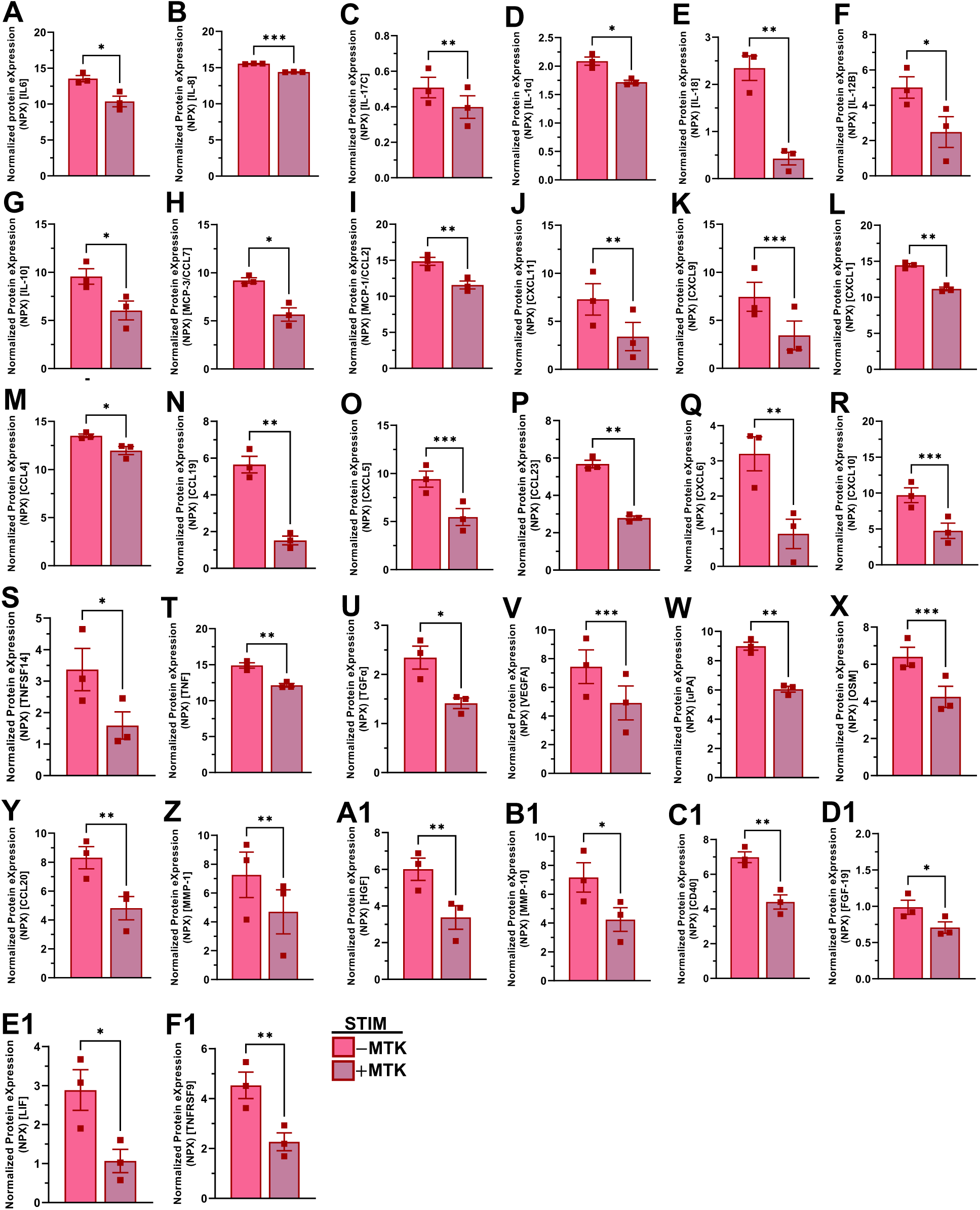
MTK-458 treatment of idiopathic PD patient-derived macrophages restricts the secretion of inflammatory mediators following stimulation with lipopolysaccharide and interleukin-1β. Using a 92-plex Olink technology, shown are individual inflammatory cyto/chemokines that were significantly lower in the supernatant of stimulated blood monocyte-derived macrophages from idiopathic PD donors treated with PINK1 activator/stabilizer (MTK-458) compared to non-treated cells. Two-tailed paired t-test was applied. Mean ± SEM, n=3 donors per group.

**Supplementary Table 1**: Full list of the genes enriched in non-bona fide myeloid cells identified in hiPSC-derived peripheral myeloid cell dataset, representing Cluster 7, 9, 14, 15, 18, 27 and 28 (in Fig. 3B).

**Supplementary Table 2**: Full list of STING-responsive genes (SRG) determined experimentally from the study conducted by Gulen *et al.,* 2023 using STING inhibitor H-151 in irradiated human fibroblasts and primary mixed cortical cells from old mice.

