## Supplemental Table 1 for "Myeloid PINK1 represses mtDNA release and immune signaling that impacts neuronal pathology in patient-derived idiopathic PD models"

**Supplementary Table 1**

| Gene | p_val | avg_log2FC | pct.1 | pct.2 | p_val_adj |
|---|---|---|---|---|---|
| ATOX1 | 0 | 10.8477477 | 0.427 | 0.78 | 0 |
| SLC16A3 | 0 | 18.0168415 | 0.319 | 0.672 | 0 |
| RAB13 | 0 | 29.5293327 | 0.42 | 0.766 | 0 |
| HCLS1 | 0 | 3.52482133 | 0.257 | 0.6 | 0 |
| BID | 0 | 3.25046399 | 0.351 | 0.683 | 0 |
| UQCR11 | 0 | 15.7806893 | 0.418 | 0.744 | 0 |
| SERPINA1 | 0 | 5.13739574 | 0.127 | 0.44 | 0 |
| ATP5MF | 0 | 5.00226543 | 0.417 | 0.728 | 0 |
| MINOS1 | 0 | 1.28370401 | 0.366 | 0.676 | 0 |
| UQCRB | 0 | 13.6993562 | 0.405 | 0.714 | 0 |
| AP2S1 | 0 | 42.5164334 | 0.479 | 0.787 | 0 |
| HM13 | 0 | 28.3437576 | 0.445 | 0.753 | 0 |
| COX6C | 0 | 11.8784034 | 0.402 | 0.71 | 0 |
| RAC1 | 0 | 20.8672389 | 0.49 | 0.797 | 0 |
| BRK1 | 0 | 10.3966588 | 0.393 | 0.698 | 0 |
| POMP | 0 | 72.6729155 | 0.422 | 0.724 | 0 |
| ATP5PF | 0 | 26.6988798 | 0.371 | 0.672 | 0 |
| EDF1 | 0 | 24.6497222 | 0.413 | 0.714 | 0 |
| ATP5MD | 0 | 23.8203407 | 0.366 | 0.665 | 0 |
| NDUFB2 | 0 | 20.5963879 | 0.386 | 0.683 | 0 |
| H3F3A | 0 | 32.3232901 | 0.47 | 0.763 | 0 |
| NDUFS5 | 0 | 28.2634067 | 0.417 | 0.71 | 0 |
| BST2 | 0 | 82.5232477 | 0.429 | 0.722 | 0 |
| SELENOW | 0 | 12.3518768 | 0.333 | 0.626 | 0 |
| ARL6IP1 | 0 | 35.0778261 | 0.371 | 0.663 | 0 |
| HIGD2A | 0 | 3.64410866 | 0.317 | 0.608 | 0 |
| TOMM7 | 0 | 4.96315846 | 0.398 | 0.688 | 0 |
| UBL5 | 0 | 57.4414278 | 0.425 | 0.712 | 0 |
| COX7B | 0 | 25.2871905 | 0.37 | 0.656 | 0 |
| CLTA | 0 | 7.96645039 | 0.478 | 0.764 | 0 |
| COX7A2 | 0 | 18.0612517 | 0.48 | 0.765 | 0 |
| NDUFA4 | 0 | 51.2445083 | 0.474 | 0.759 | 0 |
| NEDD8 | 0 | 9.40667958 | 0.412 | 0.697 | 0 |
| TMA7 | 0 | 22.5644207 | 0.438 | 0.72 | 0 |
| OST4 | 0 | 61.7307599 | 0.47 | 0.751 | 0 |
| GRINA | 0 | 27.0880283 | 0.437 | 0.718 | 0 |
| PSME2 | 0 | 20.9411394 | 0.446 | 0.726 | 0 |
| LAMTOR2 | 0 | 0.49002128 | 0.286 | 0.565 | 0 |
| HINT1 | 0 | 55.5713266 | 0.534 | 0.812 | 0 |
| SLC25A3 | 0 | 49.6759799 | 0.476 | 0.754 | 0 |

| | | | | | |
|---|---|---|---|---|---|
| COX6B1 | 0 | 20.3985189 | 0.516 | 0.794 | 0 |
| ATP5MG | 0 | 0.73428456 | 0.519 | 0.796 | 0 |
| PGAM1 | 0 | 48.4279762 | 0.462 | 0.739 | 0 |
| BRI3 | 0 | 4.62955503 | 0.245 | 0.521 | 0 |
| TCIRG1 | 0 | 2.58741962 | 0.167 | 0.442 | 0 |
| RAP1B | 0 | 12.2777223 | 0.469 | 0.744 | 0 |
| COX8A | 0 | 17.8549838 | 0.508 | 0.781 | 0 |
| PFDN5 | 0 | 1.43114292 | 0.515 | 0.785 | 0 |
| PRR13 | 0 | 14.7252085 | 0.402 | 0.671 | 0 |
| NINJ1 | 0 | 39.5459278 | 0.419 | 0.688 | 0 |
| SIRPA | 0 | 9.82762316 | 0.293 | 0.558 | 0 |
| RPL22 | 0 | 80.9443147 | 0.543 | 0.807 | 0 |
| PSMA6 | 0 | 20.946155 | 0.475 | 0.739 | 0 |
| SUMO2 | 0 | 25.3016731 | 0.489 | 0.753 | 0 |
| RPS11 | 0 | 41.1348222 | 0.506 | 0.77 | 0 |
| COX7C | 0 | 27.7135445 | 0.528 | 0.791 | 0 |
| RPL38 | 0 | 49.5585896 | 0.503 | 0.765 | 0 |
| GNG5 | 0 | 55.5713036 | 0.549 | 0.81 | 0 |
| COX6A1 | 0 | 52.6817631 | 0.533 | 0.794 | 0 |
| ATP6V1G1 | 0 | 7.77931535 | 0.493 | 0.753 | 0 |
| RPL36AL | 0 | 31.0352849 | 0.529 | 0.788 | 0 |
| YWHAB | 0 | 13.7299884 | 0.511 | 0.768 | 0 |
| BTF3 | 0 | 99.0205531 | 0.523 | 0.779 | 0 |
| HMGN2 | 0 | 36.8760752 | 0.472 | 0.723 | 0 |
| OAZ1 | 0 | 29.1220619 | 0.587 | 0.837 | 0 |
| CLU | 0 | 144.593085 | 0.294 | 0.045 | 0 |
| RPS21 | 0 | 90.1957525 | 0.548 | 0.797 | 0 |
| PGK1 | 0 | 69.9980736 | 0.55 | 0.796 | 0 |
| PPDPF | 0 | 183.944708 | 0.48 | 0.724 | 0 |
| SRP14 | 0 | 128.642543 | 0.568 | 0.806 | 0 |
| RPL37A | 0 | 113.269408 | 0.614 | 0.85 | 0 |
| NME2 | 0 | 114.721979 | 0.591 | 0.827 | 0 |
| PRDX1 | 0 | 45.4735727 | 0.626 | 0.858 | 0 |
| MIF | 0 | 98.8681617 | 0.608 | 0.838 | 0 |
| RPL27 | 0 | 106.065336 | 0.568 | 0.796 | 0 |
| ANXA2 | 0 | 56.0840951 | 0.664 | 0.892 | 0 |
| ATP5F1E | 0 | 20.9429791 | 0.631 | 0.859 | 0 |
| RPL39 | 0 | 200.634021 | 0.608 | 0.835 | 0 |
| RPL41 | 0 | 75.5031694 | 0.61 | 0.835 | 0 |
| TPI1 | 0 | 65.6963518 | 0.6 | 0.824 | 0 |
| RPL9 | 0 | 110.255269 | 0.606 | 0.828 | 0 |
| RPL24 | 0 | 206.583152 | 0.599 | 0.817 | 0 |
| RPS9 | 0 | 101.471787 | 0.644 | 0.861 | 0 |

| | | | | | |
|---|---|---|---|---|---|
| RPS2 | 0 | 185.179935 | 0.604 | 0.819 | 0 |
| RPL36 | 0 | 123.365866 | 0.637 | 0.851 | 0 |
| ENO1 | 0 | 71.4409493 | 0.642 | 0.854 | 0 |
| HLA-A | 0 | 92.6292856 | 0.602 | 0.814 | 0 |
| RPS16 | 0 | 238.79357 | 0.62 | 0.831 | 0 |
| RPL21 | 0 | 180.558879 | 0.636 | 0.838 | 0 |
| RPL3 | 0 | 142.128858 | 0.64 | 0.84 | 0 |
| UBA52 | 0 | 104.368214 | 0.672 | 0.87 | 0 |
| RPL37 | 0 | 260.362632 | 0.675 | 0.87 | 0 |
| TUBA1B | 0 | 9.40829886 | 0.635 | 0.829 | 0 |
| SERF2 | 0 | 54.1284302 | 0.706 | 0.897 | 0 |
| RACK1 | 0 | 166.658127 | 0.663 | 0.854 | 0 |
| RPS26 | 0 | 247.449744 | 0.643 | 0.834 | 0 |
| RPS25 | 0 | 121.863744 | 0.659 | 0.844 | 0 |
| RPS28 | 0 | 117.602325 | 0.686 | 0.871 | 0 |
| NACA | 0 | 74.3263943 | 0.665 | 0.846 | 0 |
| PKM | 0 | 22.3893163 | 0.738 | 0.916 | 0 |
| RPL35 | 0 | 286.392788 | 0.69 | 0.867 | 0 |
| RPLP2 | 0 | 351.296044 | 0.695 | 0.872 | 0 |
| RPL35A | 0 | 183.971162 | 0.677 | 0.854 | 0 |
| RPL6 | 0 | 101.294915 | 0.689 | 0.865 | 0 |
| HLA-B | 0 | 296.501375 | 0.691 | 0.857 | 0 |
| RPS13 | 0 | 74.3263389 | 0.708 | 0.872 | 0 |
| RPL18 | 0 | 261.876694 | 0.701 | 0.865 | 0 |
| FAU | 0 | 97.4094596 | 0.701 | 0.864 | 0 |
| HLA-C | 0 | 182.528467 | 0.661 | 0.824 | 0 |
| RPS3A | 0 | 98.856209 | 0.699 | 0.86 | 0 |
| RPL15 | 0 | 134.923102 | 0.711 | 0.871 | 0 |
| RPL29 | 0 | 142.132804 | 0.705 | 0.865 | 0 |
| ALDOA | 0 | 42.5870375 | 0.705 | 0.86 | 0 |
| RPS7 | 0 | 132.034141 | 0.726 | 0.879 | 0 |
| RPL18A | 0 | 316.699102 | 0.702 | 0.855 | 0 |
| PPIA | 0 | 144.992151 | 0.754 | 0.906 | 0 |
| RPL11 | 0 | 65.6714837 | 0.746 | 0.891 | 0 |
| RPL14 | 0 | 155.091076 | 0.728 | 0.873 | 0 |
| RPL26 | 0 | 143.574504 | 0.737 | 0.882 | 0 |
| RPS15A | 0 | 248.822279 | 0.741 | 0.881 | 0 |
| H3F3B | 0 | 230.33647 | 0.736 | 0.875 | 0 |
| RPL7A | 0 | 88.936834 | 0.738 | 0.875 | 0 |
| RPL30 | 0 | 243.121659 | 0.746 | 0.882 | 0 |
| UBC | 0 | 25.2747731 | 0.737 | 0.873 | 0 |
| RPS24 | 0 | 106.065654 | 0.76 | 0.893 | 0 |
| PTMA | 0 | 251.777829 | 0.77 | 0.901 | 0 |

| | | | | | |
|---|---|---|---|---|---|
| RPS27A | 0 | 155.117252 | 0.761 | 0.891 | 0 |
| RPS8 | 0 | 257.548609 | 0.775 | 0.904 | 0 |
| RPS18 | 0 | 225.287278 | 0.76 | 0.885 | 0 |
| RPL8 | 0 | 163.773497 | 0.761 | 0.886 | 0 |
| RPS15 | 0 | 251.776448 | 0.762 | 0.887 | 0 |
| RPS14 | 0 | 217.136651 | 0.774 | 0.896 | 0 |
| RPL32 | 0 | 287.841633 | 0.769 | 0.889 | 0 |
| RPS23 | 0 | 172.429602 | 0.775 | 0.893 | 0 |
| CFL1 | 0 | 50.0599145 | 0.787 | 0.902 | 0 |
| RPL19 | 0 | 175.314992 | 0.781 | 0.894 | 0 |
| RPS19 | 0 | 432.114709 | 0.803 | 0.914 | 0 |
| RPL13 | 0 | 257.549093 | 0.798 | 0.906 | 0 |
| RPL28 | 0 | 335.454141 | 0.824 | 0.914 | 0 |
| RPL10 | 0 | 133.928777 | 0.846 | 0.928 | 0 |
| TMSB10 | 0 | 173.872297 | 0.883 | 0.959 | 0 |
| RPS12 | 0 | 654.219649 | 0.843 | 0.917 | 0 |
| MALAT1 | 0 | 20.9466229 | 0.982 | 0.994 | 0 |
| RPL13A | 7.46E-308 | 129.122596 | 0.651 | 0.845 | 1.697E-303 |
| EEF1A1 | 1.483E-306 | 238.793574 | 0.885 | 0.942 | 3.374E-302 |
| RPS6 | 5.1E-306 | 188.299247 | 0.742 | 0.877 | 1.16E-301 |
| EIF1 | 5.008E-305 | 178.200377 | 0.747 | 0.872 | 1.139E-300 |
| MGLL | 6.079E-305 | 11.7960085 | 0.298 | 0.549 | 1.383E-300 |
| TBCA | 2.113E-304 | 35.30045 | 0.386 | 0.675 | 4.808E-300 |
| RNF181 | 4.166E-303 | 21.3913479 | 0.331 | 0.621 | 9.478E-299 |
| SNX3 | 1.454E-299 | 30.4578102 | 0.356 | 0.638 | 3.308E-295 |
| TUBA1C | 1.71E-299 | 39.7016572 | 0.53 | 0.75 | 3.89E-295 |
| RPL34 | 1.468E-298 | 100.224815 | 0.653 | 0.852 | 3.34E-294 |
| MTDH | 6.571E-298 | 81.2576709 | 0.37 | 0.679 | 1.495E-293 |
| NDUFB4 | 1.222E-297 | 11.8657874 | 0.297 | 0.584 | 2.781E-293 |
| RPS29 | 3.798E-297 | 31.2485321 | 0.422 | 0.694 | 8.641E-293 |
| ATP5MPL | 6.848E-297 | 16.3894785 | 0.361 | 0.643 | 1.558E-292 |
| SEC61B | 1.859E-296 | 277.267921 | 0.441 | 0.719 | 4.229E-292 |
| SELENOH | 1.828E-295 | 7.76097928 | 0.372 | 0.653 | 4.159E-291 |
| TUBA1A | 2.754E-294 | 40.5488112 | 0.264 | 0.54 | 6.265E-290 |
| ELOC | 4.443E-293 | 18.0610972 | 0.33 | 0.617 | 1.011E-288 |
| TNFSF13B | 8.926E-293 | 18.2194068 | 0.154 | 0.406 | 2.031E-288 |
| YBX1 | 1.127E-292 | 36.3742305 | 0.72 | 0.882 | 2.564E-288 |
| DAD1 | 3.072E-291 | 65.3615678 | 0.382 | 0.669 | 6.989E-287 |
| HSP90AA1 | 3.202E-291 | 296.501375 | 0.726 | 0.868 | 7.283E-287 |
| KLF6 | 7.642E-290 | 13.7329692 | 0.362 | 0.596 | 1.738E-285 |
| SERP1 | 8.108E-289 | 116.164495 | 0.401 | 0.678 | 1.845E-284 |
| SLC25A6 | 6.428E-288 | 30.5371138 | 0.502 | 0.759 | 1.462E-283 |
| CHI3L1 | 3.208E-287 | 19.5039274 | 0.38 | 0.605 | 7.298E-283 |

| | | | | | |
|---|---|---|---|---|---|
| PNRC1 | 1.084E-285 | 35.2319408 | 0.374 | 0.635 | 2.466E-281 |
| LAP3 | 3.007E-283 | 3.68082555 | 0.415 | 0.677 | 6.841E-279 |
| NDUFA13 | 8.763E-282 | 5.33387241 | 0.355 | 0.632 | 1.994E-277 |
| ABRACL | 3.295E-281 | 1.91497044 | 0.28 | 0.552 | 7.495E-277 |
| UQCRQ | 9.089E-281 | 26.9496622 | 0.361 | 0.639 | 2.068E-276 |
| RPL27A | 1.91E-280 | 53.440897 | 0.534 | 0.774 | 4.346E-276 |
| HSBP1 | 9.376E-280 | 32.4607365 | 0.341 | 0.632 | 2.133E-275 |
| RPL23 | 1.539E-279 | 37.2643375 | 0.416 | 0.696 | 3.502E-275 |
| RPLP0 | 2.111E-279 | 362.865347 | 0.802 | 0.9 | 4.802E-275 |
| RPL7 | 1.874E-278 | 94.5501793 | 0.521 | 0.763 | 4.263E-274 |
| AC004448.2 | 2.063E-278 | 139.064047 | 0.617 | 0.812 | 4.693E-274 |
| PLTP | 2.741E-278 | 42.3425826 | 0.244 | 0.5 | 6.237E-274 |
| RPS4X | 2.87E-278 | 121.935278 | 0.658 | 0.842 | 6.53E-274 |
| RPS20 | 5.665E-277 | 20.936579 | 0.471 | 0.714 | 1.289E-272 |
| RPL31 | 3.213E-276 | 139.219673 | 0.549 | 0.782 | 7.309E-272 |
| CD99 | 9.786E-276 | 22.3291664 | 0.391 | 0.681 | 2.226E-271 |
| RPS5 | 1.215E-274 | 235.744204 | 0.635 | 0.821 | 2.763E-270 |
| RTN4 | 3.072E-274 | 20.7885283 | 0.413 | 0.707 | 6.99E-270 |
| PPIB | 2.292E-273 | 349.881092 | 0.562 | 0.786 | 5.214E-269 |
| RPL10A | 8.821E-273 | 63.2930515 | 0.582 | 0.798 | 2.007E-268 |
| UQCRH | 2.181E-270 | 29.1839369 | 0.447 | 0.711 | 4.962E-266 |
| FYB1 | 1.082E-269 | 0.98913636 | 0.171 | 0.414 | 2.46E-265 |
| FKBP1A | 6.756E-269 | 29.4987793 | 0.407 | 0.681 | 1.537E-264 |
| MMP24OS | 2.764E-267 | 1.69726008 | 0.15 | 0.39 | 6.289E-263 |
| TUBB | 5.633E-266 | 44.0297431 | 0.585 | 0.792 | 1.281E-261 |
| RPL23A | 1.117E-263 | 96.1638453 | 0.555 | 0.784 | 2.541E-259 |
| ZNF706 | 4.191E-262 | 12.2865155 | 0.305 | 0.58 | 9.534E-258 |
| RAB5IF | 9.543E-262 | 3.20404722 | 0.339 | 0.599 | 2.171E-257 |
| ROMO1 | 4.159E-261 | 5.25788192 | 0.29 | 0.55 | 9.463E-257 |
| RPS3 | 4.714E-261 | 142.133006 | 0.77 | 0.885 | 1.072E-256 |
| EIF3K | 2.08E-260 | 11.2966879 | 0.461 | 0.72 | 4.732E-256 |
| ARL6IP5 | 6.427E-260 | 3.6334195 | 0.364 | 0.614 | 1.462E-255 |
| CYSTM1 | 2.922E-259 | 17.6305235 | 0.229 | 0.481 | 6.648E-255 |
| MYL12B | 2.306E-258 | 25.2035109 | 0.498 | 0.714 | 5.246E-254 |
| ELOB | 9.279E-258 | 23.8493066 | 0.474 | 0.73 | 2.111E-253 |
| GSTO1 | 3.407E-257 | 50.5788383 | 0.508 | 0.748 | 7.752E-253 |
| PSMB6 | 1.47E-255 | 6.51034869 | 0.376 | 0.646 | 3.345E-251 |
| PRRC2C | 2.209E-254 | 52.6847492 | 0.493 | 0.753 | 5.026E-250 |
| ATP5F1B | 2.587E-253 | 38.186868 | 0.533 | 0.765 | 5.885E-249 |
| RBX1 | 1.208E-252 | 0.56122729 | 0.351 | 0.615 | 2.749E-248 |
| ACAP2 | 3.219E-252 | 2.65690668 | 0.361 | 0.613 | 7.323E-248 |
| PTPN1 | 1.862E-251 | 34.3149483 | 0.278 | 0.525 | 4.236E-247 |
| KRTCAP2 | 9.558E-251 | 37.1251831 | 0.379 | 0.641 | 2.175E-246 |

| | | | | | |
|---|---|---|---|---|---|
| ACTG1 | 5.236E-249 | 312.371021 | 0.81 | 0.895 | 1.191E-244 |
| SAP18 | 8.403E-249 | 55.1090642 | 0.389 | 0.657 | 1.912E-244 |
| RPL5 | 6.271E-248 | 93.0813757 | 0.635 | 0.827 | 1.427E-243 |
| DYNLL1 | 5.341E-247 | 314.255477 | 0.58 | 0.769 | 1.215E-242 |
| GSTP1 | 6.074E-245 | 34.1139962 | 0.503 | 0.753 | 1.382E-240 |
| GUK1 | 1.292E-244 | 28.0499259 | 0.335 | 0.607 | 2.94E-240 |
| ANAPC11 | 1.079E-241 | 11.0563517 | 0.377 | 0.636 | 2.454E-237 |
| POLR2L | 1.8E-241 | 62.5541537 | 0.38 | 0.644 | 4.095E-237 |
| H2AFZ | 6.031E-241 | 106.065146 | 0.377 | 0.645 | 1.372E-236 |
| PARK7 | 9.611E-241 | 30.6025613 | 0.378 | 0.649 | 2.187E-236 |
| MT-ATP6 | 5.733E-240 | 2.53695487 | 0.524 | 0.768 | 1.304E-235 |
| RAN | 3.007E-238 | 121.761682 | 0.532 | 0.753 | 6.841E-234 |
| NDUFA2 | 4.59E-238 | 4.9550422 | 0.244 | 0.488 | 1.044E-233 |
| GNAS | 1.252E-237 | 12.8426902 | 0.298 | 0.553 | 2.848E-233 |
| PSMB3 | 1.891E-237 | 33.5225774 | 0.385 | 0.649 | 4.303E-233 |
| SQSTM1 | 4.93E-235 | 82.9789375 | 0.671 | 0.837 | 1.122E-230 |
| ISCU | 7.751E-233 | 10.383658 | 0.254 | 0.502 | 1.763E-228 |
| FKBP2 | 9.967E-233 | 33.4926785 | 0.295 | 0.565 | 2.268E-228 |
| SKP1 | 1.323E-232 | 36.8162674 | 0.439 | 0.686 | 3.009E-228 |
| TMEM50A | 2.313E-232 | 15.0116297 | 0.362 | 0.609 | 5.263E-228 |
| SEM1 | 1.792E-231 | 27.9807624 | 0.346 | 0.611 | 4.076E-227 |
| EIF5 | 2.032E-230 | 84.378652 | 0.459 | 0.691 | 4.623E-226 |
| MYDGF | 2.104E-230 | 84.9308155 | 0.366 | 0.638 | 4.787E-226 |
| ZFAS1 | 7.756E-229 | 71.8752799 | 0.431 | 0.695 | 1.765E-224 |
| C19orf53 | 3.607E-228 | 33.8238759 | 0.35 | 0.614 | 8.206E-224 |
| LAMTOR5 | 9.58E-228 | 6.56865724 | 0.369 | 0.621 | 2.179E-223 |
| CALM2 | 1.368E-227 | 23.8317209 | 0.484 | 0.718 | 3.113E-223 |
| EEF1B2 | 4.546E-227 | 65.6701589 | 0.514 | 0.744 | 1.034E-222 |
| CD82 | 2.701E-226 | 15.1665102 | 0.395 | 0.619 | 6.144E-222 |
| CHMP4B | 2.133E-225 | 14.3601942 | 0.258 | 0.512 | 4.851E-221 |
| SCARB2 | 8.907E-224 | 10.2412459 | 0.283 | 0.531 | 2.026E-219 |
| NDUFB11 | 6.875E-223 | 7.91951885 | 0.34 | 0.595 | 1.564E-218 |
| TSPAN3 | 1.351E-222 | 11.3056844 | 0.3 | 0.554 | 3.073E-218 |
| UBB | 1.573E-222 | 92.2798749 | 0.501 | 0.716 | 3.579E-218 |
| PPP1CA | 7.542E-221 | 2.93768281 | 0.346 | 0.589 | 1.716E-216 |
| PCBP2 | 2.404E-220 | 32.6202745 | 0.559 | 0.786 | 5.468E-216 |
| ATP5F1A | 5.763E-219 | 5.8755587 | 0.372 | 0.622 | 1.311E-214 |
| WTAP | 8.542E-219 | 8.60062323 | 0.366 | 0.586 | 1.943E-214 |
| C4orf48 | 9.561E-219 | 6.41484479 | 0.231 | 0.463 | 2.175E-214 |
| PSMB1 | 2.166E-218 | 40.6896913 | 0.375 | 0.633 | 4.928E-214 |
| S100A10 | 6.77E-218 | 207.054283 | 0.482 | 0.706 | 1.54E-213 |
| NDUFC1 | 9.367E-217 | 27.0148038 | 0.262 | 0.506 | 2.131E-212 |
| LAMTOR4 | 3.757E-216 | 3.4785533 | 0.318 | 0.552 | 8.547E-212 |

| | | | | | |
|---|---|---|---|---|---|
| NDUFAB1 | 7.636E-215 | 17.3073201 | 0.313 | 0.566 | 1.737E-210 |
| H2AFJ | 1.838E-214 | 8.56255649 | 0.146 | 0.363 | 4.181E-210 |
| NAP1L1 | 2.698E-214 | 16.882079 | 0.386 | 0.643 | 6.138E-210 |
| SEC62 | 1.26E-213 | 26.3934816 | 0.315 | 0.568 | 2.868E-209 |
| PCBD1 | 3.729E-212 | 8.62901915 | 0.181 | 0.412 | 8.484E-208 |
| RAD23A | 1.538E-210 | 0.5278345 | 0.325 | 0.565 | 3.499E-206 |
| PGLS | 2.232E-210 | 5.46029525 | 0.276 | 0.52 | 5.079E-206 |
| SRSF5 | 5.121E-210 | 7.92534435 | 0.309 | 0.549 | 1.165E-205 |
| ITGB1 | 1.566E-209 | 60.8071643 | 0.405 | 0.668 | 3.562E-205 |
| RNF7 | 5.082E-209 | 27.8088635 | 0.288 | 0.552 | 1.156E-204 |
| H2AFY | 1.372E-208 | 2.1813493 | 0.371 | 0.609 | 3.122E-204 |
| RPS17 | 3.605E-208 | 93.0813745 | 0.428 | 0.669 | 8.202E-204 |
| ATP5F1D | 3.652E-208 | 19.7392488 | 0.362 | 0.611 | 8.308E-204 |
| PABPC1 | 1.606E-206 | 35.3734371 | 0.563 | 0.776 | 3.653E-202 |
| CYP1B1 | 1.897E-206 | 32.3049135 | 0.413 | 0.611 | 4.316E-202 |
| RPS27 | 7.252E-206 | 105.444601 | 0.599 | 0.807 | 1.65E-201 |
| UQCRFS1 | 4.952E-205 | 6.17097876 | 0.318 | 0.564 | 1.127E-200 |
| ATP5ME | 6.083E-205 | 6.37348633 | 0.226 | 0.456 | 1.384E-200 |
| PRDX3 | 7.645E-205 | 6.66357566 | 0.301 | 0.549 | 1.739E-200 |
| KPNB1 | 1.642E-204 | 13.202482 | 0.446 | 0.679 | 3.735E-200 |
| LDHA | 1.629E-203 | 44.0297284 | 0.532 | 0.724 | 3.707E-199 |
| TMEM258 | 3.186E-203 | 28.6612473 | 0.313 | 0.558 | 7.249E-199 |
| ARF1 | 3.443E-203 | 62.7842667 | 0.474 | 0.696 | 7.833E-199 |
| C3 | 4.025E-203 | 13.7283882 | 0.164 | 0.369 | 9.156E-199 |
| SCAND1 | 2.145E-202 | 12.4214546 | 0.283 | 0.529 | 4.881E-198 |
| HSPA8 | 2.303E-202 | 371.521517 | 0.73 | 0.856 | 5.238E-198 |
| SNRPD2 | 3.598E-202 | 56.8083479 | 0.398 | 0.647 | 8.185E-198 |
| C1orf43 | 6.877E-202 | 0.718045 | 0.265 | 0.495 | 1.565E-197 |
| MRPL14 | 1.264E-201 | 5.98328101 | 0.251 | 0.487 | 2.876E-197 |
| ATP5IF1 | 1.627E-200 | 6.55558478 | 0.273 | 0.499 | 3.702E-196 |
| NPM1 | 6.571E-200 | 178.200349 | 0.664 | 0.824 | 1.495E-195 |
| PRDX5 | 8.746E-200 | 19.4242312 | 0.348 | 0.584 | 1.99E-195 |
| AP2M1 | 3.278E-199 | 6.52508752 | 0.406 | 0.643 | 7.458E-195 |
| TRIR | 3.321E-199 | 16.5952022 | 0.371 | 0.608 | 7.556E-195 |
| MORF4L1 | 1.606E-198 | 27.0219948 | 0.435 | 0.672 | 3.653E-194 |
| RAB2A | 1.804E-198 | 13.7561346 | 0.367 | 0.609 | 4.104E-194 |
| HMGN1 | 1.9E-198 | 25.1450517 | 0.357 | 0.606 | 4.323E-194 |
| ARF3 | 9.586E-198 | 6.09013802 | 0.234 | 0.457 | 2.181E-193 |
| ALKBH7 | 1.303E-197 | 2.35407321 | 0.159 | 0.366 | 2.965E-193 |
| SLC43A2 | 3.134E-197 | 1.56982752 | 0.145 | 0.348 | 7.129E-193 |
| SEC61G | 6.693E-197 | 123.376287 | 0.349 | 0.591 | 1.523E-192 |
| COPE | 2.453E-196 | 14.5067638 | 0.348 | 0.585 | 5.58E-192 |
| MMP14 | 6.083E-196 | 13.8461728 | 0.41 | 0.62 | 1.384E-191 |

| | | | | | |
|---|---|---|---|---|---|
| HNRNPA2B1 | 1.218E-195 | 139.248095 | 0.691 | 0.856 | 2.771E-191 |
| UBE2D3 | 4.509E-195 | 49.7108343 | 0.453 | 0.694 | 1.026E-190 |
| NDUFB7 | 4.854E-195 | 9.03966963 | 0.295 | 0.528 | 1.104E-190 |
| TMEM179B | 9.132E-195 | 4.56414799 | 0.264 | 0.488 | 2.078E-190 |
| SNRPG | 1.012E-194 | 28.632507 | 0.335 | 0.582 | 2.302E-190 |
| LAMTOR1 | 1.073E-194 | 2.13359106 | 0.358 | 0.594 | 2.44E-190 |
| MARCH1 | 2.911E-194 | 1.13486195 | 0.116 | 0.312 | 6.623E-190 |
| UBE2L3 | 7.429E-194 | 4.1804152 | 0.327 | 0.567 | 1.69E-189 |
| REEP5 | 1.266E-193 | 4.76757205 | 0.305 | 0.537 | 2.881E-189 |
| SPG21 | 1.316E-193 | 0.8751358 | 0.275 | 0.501 | 2.995E-189 |
| MRPL57 | 7.163E-193 | 7.18902085 | 0.27 | 0.506 | 1.63E-188 |
| P2RY6 | 3.275E-192 | 6.55993766 | 0.079 | 0.259 | 7.451E-188 |
| FAM162A | 1.011E-191 | 6.5337552 | 0.188 | 0.409 | 2.301E-187 |
| ZYX | 1.65E-191 | 9.0229729 | 0.328 | 0.56 | 3.755E-187 |
| LIMS1 | 6.156E-191 | 2.21315579 | 0.41 | 0.643 | 1.4E-186 |
| MTPN | 1.547E-190 | 3.40607361 | 0.334 | 0.556 | 3.52E-186 |
| CORO1B | 7.507E-190 | 3.20332065 | 0.252 | 0.467 | 1.708E-185 |
| SDHC | 1.089E-189 | 1.87590075 | 0.287 | 0.52 | 2.477E-185 |
| NDUFS6 | 1.185E-189 | 10.4330772 | 0.306 | 0.544 | 2.696E-185 |
| ZFP36L1 | 7.446E-188 | 64.2274459 | 0.339 | 0.574 | 1.694E-183 |
| DNAJC15 | 1.067E-187 | 5.50991492 | 0.179 | 0.387 | 2.428E-183 |
| NDUFB9 | 3.662E-187 | 27.710445 | 0.405 | 0.642 | 8.331E-183 |
| ATP6V0D1 | 9.782E-187 | 10.0412814 | 0.269 | 0.492 | 2.225E-182 |
| ATP5PB | 2.336E-186 | 5.90370172 | 0.373 | 0.612 | 5.314E-182 |
| NDUFAF3 | 4.057E-186 | 15.9093221 | 0.252 | 0.481 | 9.23E-182 |
| SLC43A3 | 8.853E-186 | 10.6005275 | 0.218 | 0.423 | 2.014E-181 |
| TCF25 | 1.93E-185 | 20.2156393 | 0.328 | 0.569 | 4.391E-181 |
| EXOC4 | 1.979E-185 | 0.74564784 | 0.275 | 0.481 | 4.502E-181 |
| ENY2 | 9.062E-185 | 22.3265345 | 0.316 | 0.555 | 2.062E-180 |
| PDXK | 1.75E-183 | 3.38573887 | 0.222 | 0.434 | 3.981E-179 |
| TRAPPC1 | 9.987E-183 | 8.1179611 | 0.282 | 0.503 | 2.272E-178 |
| UFC1 | 5.554E-182 | 5.83569162 | 0.297 | 0.522 | 1.264E-177 |
| HCFC1R1 | 7.334E-182 | 2.44851223 | 0.119 | 0.306 | 1.668E-177 |
| ENG | 2.051E-181 | 3.68456275 | 0.235 | 0.454 | 4.666E-177 |
| MRPS21 | 6.512E-181 | 15.8286179 | 0.265 | 0.5 | 1.481E-176 |
| DYNLT1 | 9.686E-181 | 24.2772668 | 0.307 | 0.532 | 2.204E-176 |
| C11orf58 | 1.193E-180 | 28.1643405 | 0.38 | 0.618 | 2.714E-176 |
| UQCRC1 | 3.684E-180 | 4.83804868 | 0.29 | 0.522 | 8.381E-176 |
| GPI | 4.681E-180 | 13.5407109 | 0.383 | 0.604 | 1.065E-175 |
| STMP1 | 1.843E-179 | 4.2627954 | 0.27 | 0.483 | 4.193E-175 |
| MRPL41 | 5.947E-179 | 6.47203391 | 0.312 | 0.533 | 1.353E-174 |
| GABARAPL2 | 2.534E-178 | 21.1532223 | 0.27 | 0.496 | 5.765E-174 |
| HMGN3 | 1.343E-177 | 10.2287746 | 0.213 | 0.42 | 3.054E-173 |

| | | | | | |
|---|---|---|---|---|---|
| JTB | 3.955E-177 | 1.43704958 | 0.291 | 0.5 | 8.998E-173 |
| MTHFD2 | 5.944E-177 | 0.83966265 | 0.386 | 0.592 | 1.352E-172 |
| UBE2D2 | 4.893E-176 | 12.0271122 | 0.343 | 0.576 | 1.113E-171 |
| NBDY | 1.233E-175 | 8.2566168 | 0.242 | 0.464 | 2.804E-171 |
| SUMO3 | 2.069E-175 | 4.59221166 | 0.243 | 0.451 | 4.707E-171 |
| PDE4DIP | 7.255E-175 | 0.56453053 | 0.322 | 0.513 | 1.651E-170 |
| TMEM14C | 1.074E-174 | 28.9104913 | 0.288 | 0.506 | 2.444E-170 |
| YWHAZ | 2.155E-174 | 12.12377 | 0.578 | 0.754 | 4.904E-170 |
| NDUFB10 | 3.744E-174 | 5.25429851 | 0.301 | 0.525 | 8.518E-170 |
| BNIP3L | 1.171E-173 | 25.9560173 | 0.305 | 0.516 | 2.664E-169 |
| FAM96B | 7.289E-173 | 19.058791 | 0.331 | 0.557 | 1.658E-168 |
| GADD45GIP1 | 9.512E-173 | 19.1254149 | 0.297 | 0.532 | 2.164E-168 |
| PDCD5 | 3.417E-172 | 46.9274552 | 0.313 | 0.552 | 7.773E-168 |
| NDUFS8 | 3.75E-172 | 12.9340466 | 0.325 | 0.547 | 8.531E-168 |
| POU2F2 | 1.203E-171 | 0.49433347 | 0.148 | 0.335 | 2.738E-167 |
| KTN1 | 1.311E-171 | 11.538161 | 0.295 | 0.521 | 2.984E-167 |
| TPT1 | 2.161E-171 | 140.690311 | 0.879 | 0.927 | 4.916E-167 |
| TIMM8B | 1.589E-170 | 23.2565047 | 0.266 | 0.493 | 3.615E-166 |
| PCBP1 | 1.923E-170 | 10.8083212 | 0.439 | 0.651 | 4.376E-166 |
| VDAC2 | 3.776E-170 | 22.3221928 | 0.357 | 0.586 | 8.591E-166 |
| COX14 | 4.48E-170 | 1.87010715 | 0.182 | 0.381 | 1.019E-165 |
| PSMD4 | 6.393E-170 | 9.20247357 | 0.312 | 0.533 | 1.454E-165 |
| TOMM5 | 1.094E-169 | 55.9464345 | 0.3 | 0.539 | 2.488E-165 |
| C18orf32 | 2.309E-169 | 0.5528268 | 0.144 | 0.329 | 5.253E-165 |
| DEK | 1.102E-168 | 8.40076391 | 0.314 | 0.524 | 2.508E-164 |
| PRELID1 | 1.774E-168 | 19.9569041 | 0.379 | 0.605 | 4.035E-164 |
| RPL17 | 2.136E-168 | 57.4211968 | 0.489 | 0.706 | 4.86E-164 |
| NRP2 | 2.779E-168 | 32.4843804 | 0.351 | 0.574 | 6.321E-164 |
| GLA | 2.908E-168 | 7.17641737 | 0.146 | 0.33 | 6.617E-164 |
| CD59 | 3.455E-168 | 7.45966227 | 0.341 | 0.573 | 7.861E-164 |
| SRRM1 | 5.116E-168 | 3.81516027 | 0.345 | 0.568 | 1.164E-163 |
| TRAPPC2L | 1.456E-167 | 21.9576489 | 0.225 | 0.435 | 3.312E-163 |
| MESD | 4.344E-167 | 7.35377701 | 0.269 | 0.481 | 9.882E-163 |
| PSMA1 | 1.901E-166 | 13.709535 | 0.307 | 0.537 | 4.324E-162 |
| RHEB | 3.15E-166 | 0.95705388 | 0.298 | 0.518 | 7.167E-162 |
| ERH | 7.052E-166 | 59.3208372 | 0.355 | 0.589 | 1.604E-161 |
| HSPE1 | 5.327E-165 | 137.804784 | 0.357 | 0.59 | 1.212E-160 |
| MTCH2 | 7.629E-165 | 1.04210809 | 0.266 | 0.487 | 1.736E-160 |
| TOM1 | 1.172E-164 | 1.17873994 | 0.25 | 0.449 | 2.666E-160 |
| PSENEN | 5.324E-164 | 3.59515508 | 0.203 | 0.407 | 1.211E-159 |
| CCNI | 2.874E-163 | 22.1468065 | 0.42 | 0.641 | 6.539E-159 |
| APRT | 7.787E-162 | 27.6993146 | 0.308 | 0.539 | 1.772E-157 |
| NDUFB3 | 6.372E-161 | 5.52054709 | 0.231 | 0.427 | 1.45E-156 |

| | | | | | |
|---|---|---|---|---|---|
| CDC37 | 6.842E-161 | 20.9136088 | 0.432 | 0.649 | 1.557E-156 |
| TMEM14B | 2.224E-160 | 6.77063718 | 0.224 | 0.426 | 5.059E-156 |
| PTGES3 | 6.486E-160 | 50.8009653 | 0.448 | 0.666 | 1.476E-155 |
| HNRNPC | 7.202E-160 | 41.0741285 | 0.485 | 0.712 | 1.639E-155 |
| DECR1 | 5.909E-159 | 0.92296502 | 0.193 | 0.387 | 1.344E-154 |
| ST13 | 6.607E-158 | 10.8555065 | 0.398 | 0.618 | 1.503E-153 |
| AURKAIP1 | 7.348E-158 | 18.7050073 | 0.287 | 0.505 | 1.672E-153 |
| RPS19BP1 | 2.609E-157 | 4.7880104 | 0.263 | 0.47 | 5.934E-153 |
| RAB1A | 1.202E-156 | 9.34909638 | 0.333 | 0.55 | 2.734E-152 |
| XRCC5 | 1.378E-156 | 6.04049155 | 0.435 | 0.661 | 3.136E-152 |
| UBXN4 | 1.751E-155 | 15.1848504 | 0.328 | 0.553 | 3.983E-151 |
| MYO9B | 2.653E-155 | 1.45702339 | 0.291 | 0.488 | 6.037E-151 |
| PSMB5 | 1.581E-153 | 5.54417785 | 0.3 | 0.528 | 3.596E-149 |
| SEC11A | 2.511E-153 | 28.9935799 | 0.364 | 0.595 | 5.713E-149 |
| ARF5 | 3.282E-153 | 7.2106607 | 0.252 | 0.458 | 7.466E-149 |
| CALR | 1.099E-152 | 149.346481 | 0.571 | 0.772 | 2.499E-148 |
| HSPB1 | 3.422E-151 | 153.674566 | 0.334 | 0.538 | 7.785E-147 |
| CNIH4 | 4.86E-151 | 6.14979427 | 0.213 | 0.413 | 1.106E-146 |
| WDR83OS | 5.806E-151 | 33.9308452 | 0.361 | 0.585 | 1.321E-146 |
| RGL1 | 6.666E-151 | 12.0723045 | 0.255 | 0.422 | 1.517E-146 |
| CHMP5 | 9.954E-151 | 20.1818301 | 0.326 | 0.526 | 2.264E-146 |
| KDELR1 | 1.178E-150 | 58.45743 | 0.331 | 0.557 | 2.68E-146 |
| RALA | 1.983E-149 | 27.6141555 | 0.319 | 0.524 | 4.512E-145 |
| RER1 | 3.058E-149 | 12.6337685 | 0.26 | 0.474 | 6.956E-145 |
| SSR4 | 9.413E-149 | 91.221407 | 0.381 | 0.606 | 2.141E-144 |
| ROCK1 | 3.582E-148 | 0.92426771 | 0.295 | 0.485 | 8.149E-144 |
| DPP7 | 3.595E-148 | 2.41300631 | 0.2 | 0.385 | 8.179E-144 |
| MDH2 | 4.391E-148 | 13.8268013 | 0.357 | 0.577 | 9.99E-144 |
| HSP90B1 | 7.009E-148 | 393.161943 | 0.579 | 0.774 | 1.595E-143 |
| CAST | 1.909E-147 | 3.63403047 | 0.337 | 0.537 | 4.344E-143 |
| TPM4 | 2.718E-147 | 116.081089 | 0.45 | 0.661 | 6.184E-143 |
| SOD1 | 2.821E-147 | 52.3049445 | 0.338 | 0.549 | 6.419E-143 |
| SRSF9 | 2.909E-147 | 20.2762828 | 0.255 | 0.469 | 6.617E-143 |
| MRPL51 | 1.878E-146 | 34.7719896 | 0.32 | 0.534 | 4.271E-142 |
| HSP90AB1 | 2.118E-146 | 169.544215 | 0.742 | 0.864 | 4.818E-142 |
| TMED10 | 7.648E-146 | 31.0381077 | 0.299 | 0.527 | 1.74E-141 |
| PPP4C | 1.886E-145 | 2.81146868 | 0.296 | 0.501 | 4.291E-141 |
| MRPL52 | 2.292E-145 | 10.1998087 | 0.312 | 0.513 | 5.213E-141 |
| SERBP1 | 2.417E-145 | 80.7884103 | 0.424 | 0.646 | 5.498E-141 |
| ATP1A1 | 3.698E-145 | 10.1486015 | 0.237 | 0.428 | 8.413E-141 |
| CEBPB | 1.061E-144 | 15.6372751 | 0.172 | 0.35 | 2.413E-140 |
| SFT2D1 | 1.13E-144 | 3.4272288 | 0.242 | 0.439 | 2.571E-140 |
| DUSP23 | 1.588E-144 | 0.9215918 | 0.146 | 0.316 | 3.613E-140 |

| | | | | | |
|---|---|---|---|---|---|
| RAB14 | 4.697E-144 | 1.10180192 | 0.29 | 0.486 | 1.068E-139 |
| NDUFB5 | 9.378E-144 | 0.65668978 | 0.223 | 0.41 | 2.134E-139 |
| TOMM20 | 1.448E-143 | 11.0349923 | 0.291 | 0.506 | 3.294E-139 |
| PTTG1IP | 1.851E-143 | 28.5814668 | 0.347 | 0.544 | 4.211E-139 |
| EIF4G2 | 2.879E-143 | 38.1275229 | 0.48 | 0.673 | 6.551E-139 |
| VPS28 | 2.933E-143 | 3.75848154 | 0.323 | 0.531 | 6.673E-139 |
| PSMD7 | 3.64E-143 | 19.2160041 | 0.331 | 0.546 | 8.282E-139 |
| PARP1 | 5.854E-143 | 8.2836945 | 0.237 | 0.428 | 1.332E-138 |
| TMEM165 | 1.642E-142 | 24.9303467 | 0.308 | 0.518 | 3.737E-138 |
| YWHAE | 4.139E-142 | 9.40354975 | 0.403 | 0.615 | 9.416E-138 |
| SON | 5.476E-142 | 31.9797225 | 0.443 | 0.643 | 1.246E-137 |
| PDCD6 | 9.83E-142 | 12.4399615 | 0.288 | 0.504 | 2.236E-137 |
| SRRM2 | 1.252E-141 | 15.4668497 | 0.36 | 0.564 | 2.849E-137 |
| GNB2 | 2.077E-141 | 8.74605007 | 0.287 | 0.483 | 4.726E-137 |
| GHITM | 7.807E-141 | 8.86398707 | 0.331 | 0.539 | 1.776E-136 |
| P4HB | 8.335E-141 | 233.370347 | 0.436 | 0.646 | 1.896E-136 |
| TGM2 | 1.151E-140 | 78.1237484 | 0.101 | 0.256 | 2.618E-136 |
| EID1 | 3.049E-140 | 34.8500837 | 0.298 | 0.519 | 6.936E-136 |
| TAX1BP1 | 3.677E-140 | 0.65435368 | 0.339 | 0.533 | 8.364E-136 |
| PSMA5 | 7.156E-140 | 8.02538825 | 0.317 | 0.532 | 1.628E-135 |
| TNFAIP2 | 7.421E-140 | 0.70526765 | 0.137 | 0.298 | 1.688E-135 |
| FAM177A1 | 8.933E-140 | 13.7636578 | 0.194 | 0.371 | 2.032E-135 |
| APIP | 5.676E-139 | 3.05401798 | 0.169 | 0.34 | 1.291E-134 |
| PTPN2 | 2.376E-138 | 0.57971732 | 0.239 | 0.416 | 5.406E-134 |
| C1QBP | 3.428E-138 | 56.4195522 | 0.313 | 0.529 | 7.798E-134 |
| JOSD2 | 1.648E-137 | 1.93826698 | 0.151 | 0.318 | 3.75E-133 |
| CANX | 4.186E-137 | 89.741299 | 0.507 | 0.71 | 9.523E-133 |
| CYCS | 6.538E-137 | 10.177035 | 0.315 | 0.52 | 1.487E-132 |
| SNRPB | 7.817E-137 | 27.9791928 | 0.356 | 0.565 | 1.778E-132 |
| CYC1 | 9.687E-137 | 12.2719988 | 0.271 | 0.481 | 2.204E-132 |
| FKBP8 | 3.333E-136 | 4.11570319 | 0.229 | 0.418 | 7.583E-132 |
| RPL4 | 7.798E-136 | 90.226977 | 0.606 | 0.776 | 1.774E-131 |
| RBM3 | 1.355E-135 | 9.20602938 | 0.368 | 0.561 | 3.083E-131 |
| SSBP1 | 7.733E-134 | 11.4696987 | 0.321 | 0.532 | 1.759E-129 |
| EIF2S2 | 1.019E-133 | 10.8070712 | 0.299 | 0.5 | 2.319E-129 |
| SLIRP | 1.097E-133 | 11.1142676 | 0.242 | 0.442 | 2.495E-129 |
| SP100 | 1.189E-133 | 12.373107 | 0.39 | 0.572 | 2.706E-129 |
| CDC123 | 1.911E-133 | 1.72971244 | 0.295 | 0.497 | 4.348E-129 |
| CSNK1A1 | 3.966E-133 | 24.1828366 | 0.37 | 0.583 | 9.022E-129 |
| UBXN1 | 4.268E-133 | 10.4327994 | 0.301 | 0.508 | 9.71E-129 |
| CLTC | 1.814E-132 | 20.3367773 | 0.421 | 0.612 | 4.126E-128 |
| EEF1D | 2.565E-131 | 45.5517928 | 0.487 | 0.676 | 5.835E-127 |
| SUPT4H1 | 3.456E-131 | 6.1298868 | 0.261 | 0.45 | 7.863E-127 |

| | | | | | |
|---|---|---|---|---|---|
| SF3B5 | 5.636E-131 | 62.1984167 | 0.276 | 0.49 | 1.282E-126 |
| ATP1B3 | 7.253E-131 | 12.2269166 | 0.284 | 0.496 | 1.65E-126 |
| MRPL54 | 1.922E-130 | 6.17699515 | 0.214 | 0.397 | 4.374E-126 |
| ST3GAL1 | 2.057E-130 | 25.2352097 | 0.244 | 0.415 | 4.679E-126 |
| NDUFS2 | 2.585E-130 | 9.6243436 | 0.253 | 0.442 | 5.88E-126 |
| HSPD1 | 3.208E-130 | 60.0941926 | 0.47 | 0.677 | 7.298E-126 |
| COMMD6 | 7.102E-130 | 6.10075526 | 0.282 | 0.467 | 1.616E-125 |
| RPS4Y1 | 1.852E-129 | 51.2474178 | 0.438 | 0.653 | 4.213E-125 |
| MRPS36 | 2.777E-129 | 1.50117184 | 0.192 | 0.364 | 6.318E-125 |
| MT-ND6 | 2.967E-129 | 93.0813745 | 0.469 | 0.669 | 6.749E-125 |
| ZFAND6 | 3.03E-129 | 1.97661356 | 0.244 | 0.419 | 6.893E-125 |
| VCP | 3.587E-129 | 32.8534108 | 0.461 | 0.65 | 8.161E-125 |
| SQLE | 3.761E-129 | 14.4076579 | 0.215 | 0.397 | 8.557E-125 |
| CIRBP | 5.01E-129 | 39.4388971 | 0.307 | 0.503 | 1.14E-124 |
| HMGB1 | 5.935E-129 | 33.930879 | 0.461 | 0.663 | 1.35E-124 |
| PSMD8 | 9.335E-129 | 24.0289074 | 0.417 | 0.611 | 2.124E-124 |
| RPSA | 1.005E-128 | 98.8521546 | 0.581 | 0.754 | 2.287E-124 |
| SARAF | 1.077E-128 | 41.1427533 | 0.311 | 0.504 | 2.449E-124 |
| SNU13 | 1.685E-128 | 50.1204313 | 0.328 | 0.541 | 3.834E-124 |
| SSB | 2.901E-128 | 7.95018936 | 0.34 | 0.537 | 6.599E-124 |
| TMED9 | 5.546E-128 | 58.2444 | 0.359 | 0.566 | 1.262E-123 |
| PSMB10 | 5.598E-128 | 13.0810908 | 0.271 | 0.457 | 1.274E-123 |
| SPTAN1 | 9.065E-128 | 4.98973024 | 0.277 | 0.455 | 2.062E-123 |
| CAPNS1 | 1.059E-127 | 0.25073229 | 0.159 | 0.32 | 2.41E-123 |
| ZC3H12A | 1.193E-127 | 10.6738567 | 0.142 | 0.301 | 2.713E-123 |
| LSM4 | 1.355E-127 | 0.28086841 | 0.248 | 0.432 | 3.083E-123 |
| NDUFA12 | 1.69E-127 | 18.1543874 | 0.273 | 0.472 | 3.846E-123 |
| PSMA3 | 1.831E-127 | 6.88563695 | 0.295 | 0.498 | 4.165E-123 |
| GTF3C6 | 6.663E-127 | 8.04440357 | 0.27 | 0.462 | 1.516E-122 |
| HIF1A | 4.537E-126 | 45.3769334 | 0.378 | 0.565 | 1.032E-121 |
| EIF5A | 1.26E-125 | 136.369555 | 0.492 | 0.67 | 2.867E-121 |
| WDFY3 | 1.602E-125 | 0.46253568 | 0.171 | 0.329 | 3.644E-121 |
| LSM6 | 1.224E-124 | 8.84042521 | 0.204 | 0.378 | 2.785E-120 |
| FXYD5 | 2.975E-124 | 9.40992156 | 0.487 | 0.682 | 6.769E-120 |
| RPS27L | 4.255E-124 | 29.5493446 | 0.364 | 0.565 | 9.681E-120 |
| NAA20 | 6.651E-124 | 5.61532014 | 0.251 | 0.443 | 1.513E-119 |
| ADGRE2 | 1.497E-123 | 1.07462303 | 0.142 | 0.295 | 3.405E-119 |
| NUCKS1 | 1.566E-123 | 6.25292705 | 0.318 | 0.524 | 3.563E-119 |
| DNAJA1 | 2.3E-123 | 163.74176 | 0.437 | 0.629 | 5.234E-119 |
| FDPS | 2.387E-123 | 14.9830284 | 0.214 | 0.39 | 5.43E-119 |
| ADAM9 | 4.002E-123 | 0.46314667 | 0.215 | 0.38 | 9.104E-119 |
| POLR2J | 4.099E-123 | 10.8477314 | 0.232 | 0.41 | 9.324E-119 |
| DNAJC7 | 8.387E-123 | 4.70674139 | 0.279 | 0.462 | 1.908E-118 |

| | | | | | |
|---|---|---|---|---|---|
| NDUFS3 | 1.801E-122 | 5.16193991 | 0.229 | 0.412 | 4.097E-118 |
| ZFAND3 | 2.724E-122 | 9.88427072 | 0.338 | 0.511 | 6.196E-118 |
| TCEA1 | 3.106E-122 | 30.7997743 | 0.275 | 0.473 | 7.066E-118 |
| ANP32A | 3.53E-122 | 16.6151354 | 0.361 | 0.537 | 8.03E-118 |
| FUS | 3.517E-121 | 135.903064 | 0.448 | 0.655 | 8.002E-117 |
| EIF3A | 8.733E-121 | 52.8169806 | 0.416 | 0.621 | 1.987E-116 |
| JUNB | 2.644E-120 | 38.4330691 | 0.322 | 0.524 | 6.015E-116 |
| BTF3L4 | 2.772E-120 | 7.3112065 | 0.215 | 0.392 | 6.307E-116 |
| TRMT112 | 2.978E-120 | 29.4854161 | 0.353 | 0.557 | 6.775E-116 |
| ATP5MC1 | 8.355E-120 | 12.2920699 | 0.264 | 0.457 | 1.901E-115 |
| COPS9 | 9.22E-120 | 29.0319276 | 0.251 | 0.436 | 2.098E-115 |
| PRKAR1A | 3.41E-119 | 13.7259814 | 0.375 | 0.551 | 7.758E-115 |
| EEF1G | 6.996E-119 | 25.3990168 | 0.452 | 0.649 | 1.592E-114 |
| FNDC3B | 1.02E-118 | 57.0138795 | 0.415 | 0.598 | 2.321E-114 |
| PICALM | 1.457E-118 | 3.76847019 | 0.259 | 0.427 | 3.315E-114 |
| RBM39 | 1.735E-118 | 1.97982688 | 0.358 | 0.552 | 3.948E-114 |
| RHOC | 3.908E-118 | 33.9310081 | 0.32 | 0.526 | 8.89E-114 |
| HNRNPU | 4.996E-118 | 32.4283225 | 0.54 | 0.715 | 1.137E-113 |
| IFI35 | 7.353E-118 | 17.9391707 | 0.291 | 0.447 | 1.673E-113 |
| MRPS16 | 8.436E-118 | 1.99795185 | 0.217 | 0.391 | 1.919E-113 |
| CHURC1 | 9.358E-118 | 9.28394244 | 0.248 | 0.426 | 2.129E-113 |
| ARHGEF2 | 1.05E-117 | 7.98558272 | 0.207 | 0.371 | 2.39E-113 |
| PRDX4 | 1.134E-117 | 31.5858878 | 0.235 | 0.433 | 2.58E-113 |
| LDHB | 1.973E-117 | 57.1180924 | 0.418 | 0.616 | 4.489E-113 |
| LEPROT | 3.377E-117 | 3.98477858 | 0.2 | 0.367 | 7.682E-113 |
| SPCS1 | 9.361E-117 | 25.4627148 | 0.356 | 0.557 | 2.13E-112 |
| SELENOS | 3.328E-116 | 57.4749669 | 0.235 | 0.413 | 7.57E-112 |
| ATRAID | 4.314E-116 | 11.3306412 | 0.223 | 0.397 | 9.814E-112 |
| FLNA | 2.88E-115 | 19.5043807 | 0.409 | 0.576 | 6.551E-111 |
| PRDX6 | 3.619E-115 | 31.0433221 | 0.368 | 0.568 | 8.232E-111 |
| SF3B2 | 8.867E-115 | 28.3035213 | 0.376 | 0.575 | 2.017E-110 |
| TBRG1 | 1.573E-114 | 1.85487278 | 0.211 | 0.378 | 3.579E-110 |
| SCNM1 | 1.979E-114 | 0.51763073 | 0.195 | 0.356 | 4.501E-110 |
| EIF3I | 9.894E-114 | 30.6638543 | 0.333 | 0.53 | 2.251E-109 |
| NDUFS7 | 1.672E-113 | 3.41458252 | 0.263 | 0.431 | 3.805E-109 |
| RBM8A | 1.983E-113 | 45.4723795 | 0.357 | 0.549 | 4.511E-109 |
| ADRM1 | 2.544E-112 | 20.016125 | 0.296 | 0.483 | 5.787E-108 |
| SMDT1 | 3.961E-112 | 13.6188359 | 0.195 | 0.354 | 9.011E-108 |
| AL365205.1 | 9.068E-112 | 11.5739751 | 0.188 | 0.355 | 2.063E-107 |
| HDLBP | 1.036E-111 | 34.2655663 | 0.377 | 0.577 | 2.358E-107 |
| CSDE1 | 1.13E-111 | 20.7069855 | 0.4 | 0.589 | 2.57E-107 |
| EEF2 | 1.683E-111 | 133.476836 | 0.647 | 0.789 | 3.828E-107 |
| GTF2A2 | 1.961E-111 | 24.1269956 | 0.257 | 0.438 | 4.461E-107 |

| | | | | | |
|---|---|---|---|---|---|
| YWHAG | 2.041E-111 | 6.58891303 | 0.341 | 0.511 | 4.644E-107 |
| ARL6IP4 | 2.184E-111 | 5.44259386 | 0.28 | 0.459 | 4.968E-107 |
| PHPT1 | 3.144E-111 | 10.9597084 | 0.246 | 0.422 | 7.151E-107 |
| CCT6A | 4.976E-111 | 14.3412824 | 0.418 | 0.609 | 1.132E-106 |
| BCAT1 | 6.873E-111 | 9.4828796 | 0.236 | 0.404 | 1.564E-106 |
| CYB5R3 | 7.997E-111 | 8.86168213 | 0.258 | 0.434 | 1.819E-106 |
| KIF5B | 8.51E-111 | 16.6185373 | 0.343 | 0.521 | 1.936E-106 |
| CHMP2A | 1.477E-110 | 1.16942243 | 0.237 | 0.403 | 3.36E-106 |
| OSTC | 1.547E-110 | 65.4235223 | 0.292 | 0.495 | 3.519E-106 |
| PPIG | 1.65E-110 | 6.14144675 | 0.248 | 0.418 | 3.755E-106 |
| UBE2D1 | 1.774E-110 | 1.09919502 | 0.192 | 0.35 | 4.036E-106 |
| UPP1 | 2.938E-110 | 63.4129329 | 0.247 | 0.414 | 6.684E-106 |
| RBM25 | 4.001E-110 | 17.0677781 | 0.324 | 0.51 | 9.102E-106 |
| CCT3 | 6.086E-110 | 44.0184252 | 0.39 | 0.586 | 1.385E-105 |
| MEA1 | 8.913E-110 | 15.2619045 | 0.252 | 0.434 | 2.028E-105 |
| ATP6V1A | 1.341E-109 | 0.6165763 | 0.188 | 0.341 | 3.05E-105 |
| NAA38 | 1.435E-109 | 0.67892347 | 0.191 | 0.344 | 3.264E-105 |
| B4GALT1 | 1.825E-109 | 2.40729786 | 0.231 | 0.39 | 4.153E-105 |
| BIRC6 | 3.771E-109 | 1.08719248 | 0.282 | 0.445 | 8.578E-105 |
| MLF2 | 3.975E-109 | 12.2936141 | 0.3 | 0.494 | 9.043E-105 |
| MRPL20 | 4.841E-109 | 34.6905556 | 0.279 | 0.469 | 1.101E-104 |
| JAK1 | 6.241E-109 | 28.7168695 | 0.319 | 0.498 | 1.42E-104 |
| SNRPD3 | 1.371E-108 | 28.4758439 | 0.304 | 0.495 | 3.118E-104 |
| SLC3A2 | 5.288E-108 | 136.362225 | 0.348 | 0.538 | 1.203E-103 |
| CNPY3 | 5.289E-108 | 0.71178957 | 0.181 | 0.333 | 1.203E-103 |
| MMADHC | 6.368E-108 | 48.120255 | 0.272 | 0.464 | 1.449E-103 |
| PSMB8 | 2.11E-107 | 16.5879118 | 0.242 | 0.415 | 4.801E-103 |
| NDFIP1 | 2.843E-107 | 3.16367844 | 0.26 | 0.428 | 6.467E-103 |
| SSR3 | 3.653E-107 | 55.5910908 | 0.31 | 0.507 | 8.31E-103 |
| PSMA4 | 5.157E-107 | 10.7911868 | 0.261 | 0.438 | 1.173E-102 |
| SET | 8.728E-107 | 74.8948029 | 0.351 | 0.541 | 1.986E-102 |
| TMEM167A | 1.015E-106 | 15.433696 | 0.224 | 0.391 | 2.308E-102 |
| LMAN2 | 1.229E-106 | 10.2691008 | 0.291 | 0.478 | 2.796E-102 |
| ZNFX1 | 1.853E-106 | 4.23363854 | 0.259 | 0.411 | 4.215E-102 |
| MPC2 | 1.934E-106 | 3.65005163 | 0.202 | 0.364 | 4.4E-102 |
| SRSF11 | 2.405E-106 | 6.73127249 | 0.333 | 0.525 | 5.471E-102 |
| VTI1B | 2.986E-106 | 2.12460761 | 0.206 | 0.366 | 6.793E-102 |
| RPS10 | 3.372E-106 | 33.4560763 | 0.329 | 0.523 | 7.672E-102 |
| RNH1 | 5.067E-106 | 9.05083141 | 0.298 | 0.481 | 1.153E-101 |
| PFKL | 7.584E-106 | 6.47031287 | 0.19 | 0.346 | 1.725E-101 |
| ZSWIM6 | 9.689E-106 | 0.8190176 | 0.209 | 0.357 | 2.204E-101 |
| TMEM59 | 1.288E-105 | 53.1579996 | 0.283 | 0.461 | 2.929E-101 |
| BCKDK | 1.298E-105 | 0.79783656 | 0.158 | 0.305 | 2.953E-101 |

| | | | | | |
|---|---|---|---|---|---|
| PFDN2 | 1.408E-105 | 24.4997408 | 0.308 | 0.497 | 3.203E-101 |
| USO1 | 2.621E-105 | 4.16299222 | 0.237 | 0.403 | 5.962E-101 |
| SNRPC | 2.743E-105 | 16.4608556 | 0.301 | 0.488 | 6.24E-101 |
| TGOLN2 | 3.545E-105 | 1.75357733 | 0.311 | 0.476 | 8.064E-101 |
| WDR1 | 4.505E-105 | 23.8953428 | 0.359 | 0.535 | 1.025E-100 |
| UBE2B | 4.551E-105 | 26.636896 | 0.255 | 0.426 | 1.035E-100 |
| HK2 | 7.865E-105 | 10.611284 | 0.188 | 0.329 | 1.789E-100 |
| LAMP2 | 1.028E-104 | 2.03734197 | 0.183 | 0.333 | 2.339E-100 |
| VAPA | 1.374E-104 | 4.9940367 | 0.372 | 0.553 | 3.125E-100 |
| JPT1 | 1.771E-104 | 36.1079993 | 0.247 | 0.428 | 4.03E-100 |
| NDUFA8 | 1.786E-104 | 7.01057733 | 0.199 | 0.363 | 4.062E-100 |
| GLIPR1 | 7.658E-104 | 20.8601294 | 0.175 | 0.328 | 1.742E-99 |
| DDRGK1 | 8.26E-104 | 8.34812686 | 0.218 | 0.378 | 1.879E-99 |
| DPH3 | 1.37E-103 | 22.5951027 | 0.239 | 0.4 | 3.116E-99 |
| UBE2I | 4.947E-103 | 17.2252531 | 0.336 | 0.519 | 1.1255E-98 |
| OTUB1 | 1.224E-102 | 7.09635066 | 0.252 | 0.424 | 2.7853E-98 |
| RTL8C | 1.31E-102 | 10.8217222 | 0.217 | 0.387 | 2.9814E-98 |
| MCL1 | 1.699E-102 | 17.0544026 | 0.295 | 0.458 | 3.8656E-98 |
| MRPS34 | 1.97E-102 | 11.7381191 | 0.253 | 0.426 | 4.4816E-98 |
| LYRM2 | 2.063E-102 | 1.92957715 | 0.181 | 0.336 | 4.6933E-98 |
| NOL7 | 2.674E-102 | 29.5993282 | 0.311 | 0.495 | 6.083E-98 |
| SUMO1 | 3.774E-102 | 11.6635612 | 0.259 | 0.437 | 8.5851E-98 |
| MRPS24 | 6.63E-102 | 22.1127463 | 0.244 | 0.425 | 1.5084E-97 |
| ERP29 | 7.029E-102 | 6.85117569 | 0.278 | 0.442 | 1.5991E-97 |
| CHCHD1 | 1.448E-101 | 1.94717796 | 0.181 | 0.331 | 3.2949E-97 |
| TEX264 | 2.114E-101 | 0.6005209 | 0.153 | 0.294 | 4.8093E-97 |
| SDF4 | 3.15E-101 | 36.2314082 | 0.292 | 0.482 | 7.166E-97 |
| EIF6 | 3.941E-101 | 16.6054352 | 0.306 | 0.487 | 8.966E-97 |
| BLOC1S2 | 4.608E-101 | 3.05280884 | 0.206 | 0.358 | 1.0484E-96 |
| TMEM147 | 9.951E-101 | 14.2106042 | 0.266 | 0.448 | 2.2638E-96 |
| CTSC | 1.776E-100 | 153.674554 | 0.458 | 0.627 | 4.0406E-96 |
| LAPTM4A | 2.931E-100 | 35.3735717 | 0.323 | 0.513 | 6.6683E-96 |
| TKT | 3.648E-100 | 76.1991494 | 0.442 | 0.624 | 8.2984E-96 |
| COX7A2L | 7.477E-100 | 22.400074 | 0.275 | 0.454 | 1.701E-95 |
| CORO1C | 7.635E-100 | 42.6310086 | 0.272 | 0.433 | 1.737E-95 |
| SF3B6 | 1.249E-99 | 16.7045537 | 0.259 | 0.443 | 2.842E-95 |
| ANP32B | 1.817E-99 | 42.5440826 | 0.371 | 0.555 | 4.1347E-95 |
| EIF1B | 2.276E-99 | 0.55429492 | 0.258 | 0.422 | 5.1774E-95 |
| DRAP1 | 5.634E-99 | 54.5427173 | 0.346 | 0.526 | 1.2817E-94 |
| BANF1 | 5.657E-99 | 21.9640003 | 0.321 | 0.509 | 1.2869E-94 |
| EIF3G | 1.5946E-98 | 9.09059196 | 0.336 | 0.515 | 3.6277E-94 |
| OCIAD1 | 2.2186E-98 | 17.561714 | 0.287 | 0.466 | 5.0473E-94 |
| PSMB2 | 2.951E-98 | 6.41222243 | 0.334 | 0.516 | 6.7135E-94 |

| | | | | | |
|---|---|---|---|---|---|
| RNF149 | 3.0337E-98 | 5.23985434 | 0.263 | 0.432 | 6.9017E-94 |
| IFIH1 | 4.5714E-98 | 18.3470523 | 0.212 | 0.344 | 1.04E-93 |
| SNX8 | 4.7828E-98 | 7.42722696 | 0.177 | 0.313 | 1.0881E-93 |
| CTSA | 1.575E-97 | 3.29464938 | 0.273 | 0.424 | 3.5832E-93 |
| BAZ1A | 1.6058E-97 | 3.61982004 | 0.26 | 0.419 | 3.6532E-93 |
| PAFAH1B1 | 1.8586E-97 | 2.46503816 | 0.306 | 0.466 | 4.2284E-93 |
| NFKB2 | 2.0399E-97 | 6.16008218 | 0.178 | 0.317 | 4.6407E-93 |
| TMBIM1 | 2.4247E-97 | 1.40415214 | 0.195 | 0.336 | 5.5162E-93 |
| PPP1R2 | 3.623E-97 | 1.82452651 | 0.181 | 0.324 | 8.2424E-93 |
| DDT | 5.0271E-97 | 3.9174507 | 0.172 | 0.314 | 1.1437E-92 |
| SEC14L1 | 5.718E-97 | 1.97359337 | 0.223 | 0.373 | 1.3009E-92 |
| DDX5 | 6.1951E-97 | 75.769034 | 0.585 | 0.736 | 1.4094E-92 |
| RSRC1 | 9.7627E-97 | 7.49449685 | 0.207 | 0.351 | 2.221E-92 |
| CHD4 | 2.2097E-96 | 26.4814971 | 0.269 | 0.435 | 5.027E-92 |
| PPCS | 2.4829E-96 | 0.53472442 | 0.167 | 0.309 | 5.6486E-92 |
| GTF2I | 3.1123E-96 | 0.60794536 | 0.278 | 0.441 | 7.0806E-92 |
| BNIP3 | 4.0427E-96 | 3.63897147 | 0.212 | 0.358 | 9.1972E-92 |
| MRPS18C | 4.3816E-96 | 16.3483581 | 0.231 | 0.395 | 9.9682E-92 |
| LAGE3 | 5.1028E-96 | 0.87141968 | 0.171 | 0.313 | 1.1609E-91 |
| ADIPOR1 | 5.313E-96 | 0.27384179 | 0.246 | 0.403 | 1.2087E-91 |
| RPN2 | 1.0738E-95 | 24.5760501 | 0.299 | 0.488 | 2.4429E-91 |
| GNG10 | 1.5627E-95 | 1.11459588 | 0.187 | 0.336 | 3.5552E-91 |
| ANAPC16 | 2.5535E-95 | 4.09986078 | 0.258 | 0.413 | 5.8091E-91 |
| MSN | 3.1049E-95 | 32.0295198 | 0.53 | 0.677 | 7.0637E-91 |
| FAM213A | 3.4532E-95 | 2.16954514 | 0.124 | 0.252 | 7.8561E-91 |
| CHCHD5 | 7.3939E-95 | 1.18108301 | 0.138 | 0.273 | 1.6821E-90 |
| SRA1 | 1.6843E-94 | 5.58467726 | 0.2 | 0.353 | 3.8318E-90 |
| TIMM17A | 2.6597E-94 | 48.224433 | 0.248 | 0.423 | 6.0508E-90 |
| TXN2 | 2.8507E-94 | 5.73879196 | 0.195 | 0.347 | 6.4853E-90 |
| HNRNPA3 | 3.2865E-94 | 77.3941379 | 0.451 | 0.628 | 7.4768E-90 |
| PA2G4 | 3.763E-94 | 10.554789 | 0.328 | 0.511 | 8.5609E-90 |
| AP1S2 | 4.079E-94 | 8.23978495 | 0.212 | 0.359 | 9.2796E-90 |
| UQCRC2 | 4.8222E-94 | 1.29318751 | 0.262 | 0.423 | 1.0971E-89 |
| TM2D2 | 5.2479E-94 | 2.2671336 | 0.189 | 0.337 | 1.1939E-89 |
| BAG1 | 5.3103E-94 | 8.91384042 | 0.247 | 0.409 | 1.2081E-89 |
| RRBP1 | 5.9232E-94 | 70.8030121 | 0.342 | 0.525 | 1.3475E-89 |
| TOP1 | 6.4063E-94 | 22.3813488 | 0.316 | 0.495 | 1.4574E-89 |
| MRPL34 | 1.3426E-93 | 0.86980702 | 0.185 | 0.33 | 3.0544E-89 |
| PPM1G | 4.0967E-93 | 22.1607503 | 0.309 | 0.488 | 9.3201E-89 |
| NECTIN2 | 4.2359E-93 | 29.598657 | 0.175 | 0.319 | 9.6366E-89 |
| LYPLA1 | 7.1031E-93 | 4.83867583 | 0.25 | 0.411 | 1.6159E-88 |
| ADPGK | 8.8261E-93 | 1.40303659 | 0.248 | 0.397 | 2.0079E-88 |
| EIF3D | 1.0228E-92 | 21.6316581 | 0.304 | 0.474 | 2.3269E-88 |

| | | | | | |
|---|---|---|---|---|---|
| COPZ1 | 1.6861E-92 | 15.837092 | 0.246 | 0.414 | 3.8358E-88 |
| HP1BP3 | 2.0233E-92 | 8.70454548 | 0.263 | 0.424 | 4.6031E-88 |
| COMMD1 | 2.7895E-92 | 2.76114272 | 0.231 | 0.386 | 6.3461E-88 |
| BOLA3 | 3.927E-92 | 4.38826786 | 0.156 | 0.296 | 8.934E-88 |
| RTRAF | 5.8876E-92 | 17.8388366 | 0.288 | 0.469 | 1.3394E-87 |
| TIMM13 | 6.4002E-92 | 17.0332866 | 0.243 | 0.418 | 1.4561E-87 |
| MKKS | 6.5018E-92 | 2.30360458 | 0.153 | 0.289 | 1.4792E-87 |
| CIB1 | 7.4202E-92 | 8.37946323 | 0.198 | 0.351 | 1.6881E-87 |
| RPL22L1 | 8.0565E-92 | 71.8359313 | 0.238 | 0.407 | 1.8329E-87 |
| CYP51A1 | 1.0392E-91 | 7.94661257 | 0.173 | 0.311 | 2.3641E-87 |
| FADS1 | 1.0779E-91 | 3.779083 | 0.175 | 0.316 | 2.4523E-87 |
| PRPF40A | 1.6215E-91 | 20.7749417 | 0.351 | 0.524 | 3.6889E-87 |
| RALY | 2.9045E-91 | 11.2227101 | 0.258 | 0.413 | 6.6078E-87 |
| PSMB7 | 8.896E-91 | 19.8217287 | 0.296 | 0.473 | 2.0238E-86 |
| ARHGDIA | 1.1372E-90 | 1.15403434 | 0.373 | 0.531 | 2.5872E-86 |
| MYH9 | 1.2283E-90 | 69.1676055 | 0.45 | 0.612 | 2.7944E-86 |
| PNP | 1.6835E-90 | 9.84366421 | 0.261 | 0.409 | 3.83E-86 |
| EIF4G3 | 2.0175E-90 | 4.53345396 | 0.276 | 0.428 | 4.5897E-86 |
| APP | 2.5952E-90 | 107.50832 | 0.353 | 0.547 | 5.904E-86 |
| MX2 | 4.7635E-90 | 18.4013179 | 0.227 | 0.35 | 1.0837E-85 |
| ZNF638 | 1.1968E-89 | 2.71842538 | 0.286 | 0.438 | 2.7228E-85 |
| TUFM | 1.2461E-89 | 29.5741159 | 0.274 | 0.452 | 2.8348E-85 |
| EMC6 | 1.4723E-89 | 15.9887838 | 0.237 | 0.402 | 3.3494E-85 |
| FRMD4A | 3.1697E-89 | 3.6550194 | 0.215 | 0.356 | 7.2111E-85 |
| MTCH1 | 5.4277E-89 | 43.7236025 | 0.292 | 0.469 | 1.2348E-84 |
| PFDN1 | 6.2135E-89 | 0.32042235 | 0.167 | 0.304 | 1.4136E-84 |
| BSG | 7.8424E-89 | 45.8945776 | 0.343 | 0.527 | 1.7841E-84 |
| ARF4 | 1.3208E-88 | 64.0403531 | 0.325 | 0.5 | 3.0048E-84 |
| AKR7A2 | 1.8061E-88 | 2.97539042 | 0.139 | 0.272 | 4.1088E-84 |
| CISD2 | 9.345E-88 | 14.227807 | 0.2 | 0.347 | 2.126E-83 |
| ORMDL1 | 1.0213E-87 | 15.4742016 | 0.212 | 0.361 | 2.3235E-83 |
| NUTF2 | 1.0961E-87 | 18.9533992 | 0.268 | 0.443 | 2.4936E-83 |
| KDELR2 | 1.4451E-87 | 168.509344 | 0.269 | 0.444 | 3.2876E-83 |
| DNAJC8 | 1.858E-87 | 5.39390887 | 0.307 | 0.469 | 4.2271E-83 |
| ZNHIT1 | 2.3158E-87 | 5.87375743 | 0.213 | 0.362 | 5.2684E-83 |
| HNRNPA1 | 2.4223E-87 | 94.3769474 | 0.398 | 0.576 | 5.5106E-83 |
| SPCS3 | 3.0395E-87 | 28.1583612 | 0.268 | 0.427 | 6.9148E-83 |
| ZCRB1 | 3.8264E-87 | 0.62630281 | 0.156 | 0.289 | 8.705E-83 |
| LIMD2 | 7.375E-87 | 3.95407823 | 0.257 | 0.4 | 1.6778E-82 |
| GLRX2 | 1.048E-86 | 4.83923203 | 0.168 | 0.306 | 2.3843E-82 |
| PSMC5 | 1.5906E-86 | 18.8006256 | 0.282 | 0.449 | 3.6185E-82 |
| LRP1 | 3.723E-86 | 54.2078879 | 0.254 | 0.408 | 8.4699E-82 |
| ETFA | 5.9179E-86 | 6.96041473 | 0.246 | 0.397 | 1.3463E-81 |

| | | | | | |
|---|---|---|---|---|---|
| SSU72 | 6.2891E-86 | 17.305088 | 0.26 | 0.423 | 1.4308E-81 |
| ZC3H15 | 6.3617E-86 | 18.3051769 | 0.286 | 0.453 | 1.4473E-81 |
| ITGB1BP1 | 9.2319E-86 | 5.14313951 | 0.205 | 0.35 | 2.1003E-81 |
| OS9 | 1.2694E-85 | 2.54233508 | 0.293 | 0.446 | 2.888E-81 |
| KXD1 | 2.8386E-85 | 5.90317981 | 0.219 | 0.371 | 6.4579E-81 |
| NEU1 | 2.9909E-85 | 60.7015656 | 0.245 | 0.392 | 6.8042E-81 |
| NSF | 3.0926E-85 | 2.19085042 | 0.216 | 0.351 | 7.0356E-81 |
| S100A4 | 3.757E-85 | 89.1953268 | 0.425 | 0.563 | 8.5472E-81 |
| MLX | 3.7597E-85 | 2.65510596 | 0.217 | 0.354 | 8.5534E-81 |
| TBCB | 4.2637E-85 | 10.5314548 | 0.238 | 0.387 | 9.6999E-81 |
| RALBP1 | 5.6834E-85 | 1.36330521 | 0.159 | 0.29 | 1.293E-80 |
| ECH1 | 7.8977E-85 | 4.93480813 | 0.172 | 0.301 | 1.7967E-80 |
| ACO2 | 1.2575E-84 | 2.01557086 | 0.201 | 0.343 | 2.8607E-80 |
| SEC11C | 1.5219E-84 | 8.9672796 | 0.222 | 0.377 | 3.4624E-80 |
| AUP1 | 1.6133E-84 | 25.1956322 | 0.287 | 0.464 | 3.6703E-80 |
| NEMF | 2.2527E-84 | 2.23454335 | 0.255 | 0.402 | 5.1248E-80 |
| UBE2K | 2.7309E-84 | 4.92242351 | 0.282 | 0.44 | 6.2128E-80 |
| CCT2 | 2.8995E-84 | 49.0174551 | 0.33 | 0.506 | 6.5965E-80 |
| MACF1 | 4.0965E-84 | 2.17093125 | 0.393 | 0.542 | 9.3196E-80 |
| GTF2H5 | 9.1465E-84 | 4.1874026 | 0.182 | 0.32 | 2.0808E-79 |
| CXXC5 | 1.1515E-83 | 9.79152085 | 0.14 | 0.266 | 2.6196E-79 |
| TRAM1 | 1.2894E-83 | 12.4866053 | 0.284 | 0.439 | 2.9333E-79 |
| TMEM230 | 2.0624E-83 | 5.79824826 | 0.235 | 0.383 | 4.692E-79 |
| MRPL27 | 4.1649E-83 | 7.29333875 | 0.209 | 0.358 | 9.475E-79 |
| MRFAP1 | 7.2518E-83 | 17.8914046 | 0.274 | 0.44 | 1.6498E-78 |
| ACP1 | 1.0296E-82 | 14.0883128 | 0.233 | 0.386 | 2.3424E-78 |
| SDHA | 1.2981E-82 | 3.67663606 | 0.185 | 0.32 | 2.9531E-78 |
| SEPHS2 | 1.4039E-82 | 3.51965375 | 0.181 | 0.312 | 3.1939E-78 |
| AHR | 1.9652E-82 | 3.10622447 | 0.15 | 0.271 | 4.4707E-78 |
| PLP2 | 4.0989E-82 | 10.5005833 | 0.341 | 0.472 | 9.3249E-78 |
| SELENOF | 8.4813E-82 | 15.2011545 | 0.306 | 0.48 | 1.9295E-77 |
| MIEN1 | 9.8067E-82 | 2.17601888 | 0.183 | 0.318 | 2.231E-77 |
| PIK3R1 | 1.1158E-81 | 1.493566 | 0.195 | 0.323 | 2.5384E-77 |
| MRPS7 | 1.4676E-81 | 4.44012712 | 0.217 | 0.37 | 3.3388E-77 |
| SSR2 | 1.7075E-81 | 81.9916065 | 0.355 | 0.533 | 3.8846E-77 |
| TIMM17B | 1.955E-81 | 4.1633441 | 0.199 | 0.338 | 4.4476E-77 |
| RABEP1 | 2.0575E-81 | 1.44545853 | 0.213 | 0.35 | 4.6809E-77 |
| SSNA1 | 2.8327E-81 | 3.24451325 | 0.211 | 0.357 | 6.4445E-77 |
| HNRNPDL | 3.5849E-81 | 46.9153023 | 0.429 | 0.592 | 8.1557E-77 |
| RTCB | 4.7661E-81 | 6.00835099 | 0.249 | 0.392 | 1.0843E-76 |
| CCT4 | 6.0657E-81 | 46.3462483 | 0.343 | 0.518 | 1.38E-76 |
| PSMC4 | 8.8846E-81 | 8.89159005 | 0.252 | 0.405 | 2.0213E-76 |
| SUMF2 | 9.639E-81 | 0.33660576 | 0.139 | 0.261 | 2.1929E-76 |

| | | | | | |
|---|---|---|---|---|---|
| FEZ2 | 1.1713E-80 | 0.91526901 | 0.178 | 0.31 | 2.6647E-76 |
| XRN2 | 1.691E-80 | 5.14109559 | 0.311 | 0.476 | 3.847E-76 |
| ARPC5L | 1.7874E-80 | 24.5183243 | 0.242 | 0.401 | 4.0663E-76 |
| TECR | 1.8692E-80 | 6.4301707 | 0.186 | 0.322 | 4.2525E-76 |
| HADHB | 1.9626E-80 | 1.51428088 | 0.204 | 0.344 | 4.465E-76 |
| TIMM10 | 2.3034E-80 | 6.72659285 | 0.195 | 0.338 | 5.2402E-76 |
| PAIP2 | 3.4795E-80 | 2.21359224 | 0.2 | 0.335 | 7.916E-76 |
| DMAC1 | 4.5231E-80 | 12.6496351 | 0.17 | 0.302 | 1.029E-75 |
| ITPA | 4.6207E-80 | 6.36657958 | 0.216 | 0.362 | 1.0512E-75 |
| EMC4 | 5.2054E-80 | 26.2679746 | 0.241 | 0.403 | 1.1842E-75 |
| SRI | 5.9958E-80 | 6.33518377 | 0.326 | 0.476 | 1.364E-75 |
| AKAP9 | 6.5459E-80 | 0.73794445 | 0.205 | 0.337 | 1.4892E-75 |
| H2AFV | 9.7135E-80 | 15.0937856 | 0.27 | 0.421 | 2.2098E-75 |
| RTF2 | 1.3119E-79 | 3.25420686 | 0.261 | 0.409 | 2.9845E-75 |
| LSM8 | 1.96E-79 | 8.14320411 | 0.207 | 0.345 | 4.4591E-75 |
| TRIP12 | 1.9985E-79 | 3.2610765 | 0.311 | 0.45 | 4.5466E-75 |
| RAB6A | 2.4269E-79 | 2.65380754 | 0.284 | 0.426 | 5.5212E-75 |
| XRCC6 | 3.7616E-79 | 25.3937804 | 0.357 | 0.524 | 8.5577E-75 |
| NCL | 1.1796E-78 | 384.322471 | 0.616 | 0.755 | 2.6836E-74 |
| RSU1 | 1.1929E-78 | 0.90079432 | 0.226 | 0.362 | 2.7138E-74 |
| HAX1 | 1.3525E-78 | 2.01625972 | 0.21 | 0.35 | 3.0769E-74 |
| SNRPD1 | 1.4958E-78 | 35.7027822 | 0.309 | 0.474 | 3.4029E-74 |
| POLR2K | 1.5253E-78 | 20.3681712 | 0.219 | 0.364 | 3.47E-74 |
| CAB39 | 1.7583E-78 | 1.13564965 | 0.154 | 0.271 | 4.0002E-74 |
| ACTN1 | 1.9561E-78 | 72.3100091 | 0.292 | 0.451 | 4.4502E-74 |
| CAMTA1 | 2.6247E-78 | 1.27976237 | 0.182 | 0.312 | 5.9712E-74 |
| EIF3L | 4.5087E-78 | 17.5461065 | 0.297 | 0.462 | 1.0257E-73 |
| PTP4A2 | 5.5219E-78 | 4.75108786 | 0.206 | 0.337 | 1.2562E-73 |
| COA3 | 5.7021E-78 | 3.36303638 | 0.2 | 0.342 | 1.2972E-73 |
| SIL1 | 8.7528E-78 | 3.02040187 | 0.201 | 0.337 | 1.9913E-73 |
| FUNDC2 | 9.7615E-78 | 7.1745465 | 0.232 | 0.379 | 2.2207E-73 |
| ACAA2 | 1.7854E-77 | 0.44578081 | 0.205 | 0.342 | 4.0618E-73 |
| CACYBP | 1.796E-77 | 8.26828339 | 0.277 | 0.431 | 4.0858E-73 |
| PHB | 1.882E-77 | 35.2817047 | 0.297 | 0.467 | 4.2816E-73 |
| PTMS | 1.9612E-77 | 19.716671 | 0.207 | 0.346 | 4.4618E-73 |
| ESD | 2.8806E-77 | 4.40175911 | 0.213 | 0.354 | 6.5534E-73 |
| CUTA | 3.812E-77 | 31.5602275 | 0.27 | 0.418 | 8.6722E-73 |
| RWDD1 | 7.7857E-77 | 9.80294972 | 0.233 | 0.383 | 1.7712E-72 |
| C7orf50 | 8.0782E-77 | 10.8111371 | 0.175 | 0.302 | 1.8378E-72 |
| AKIRIN1 | 8.5145E-77 | 3.71254337 | 0.215 | 0.352 | 1.937E-72 |
| MRPL17 | 3.8989E-76 | 6.72475858 | 0.192 | 0.331 | 8.8701E-72 |
| VPS36 | 5.3661E-76 | 1.11248673 | 0.188 | 0.318 | 1.2208E-71 |
| NRDC | 7.3473E-76 | 8.8729994 | 0.298 | 0.443 | 1.6715E-71 |

| | | | | | |
|---|---|---|---|---|---|
| PPP1R12A | 1.038E-75 | 0.3831322 | 0.313 | 0.456 | 2.3614E-71 |
| UQCC2 | 1.2993E-75 | 5.38623853 | 0.232 | 0.378 | 2.9558E-71 |
| OXA1L | 1.5939E-75 | 5.79112437 | 0.224 | 0.364 | 3.6261E-71 |
| G3BP1 | 2.0506E-75 | 15.6308154 | 0.374 | 0.542 | 4.6651E-71 |
| FLOT1 | 5.5884E-75 | 2.23417741 | 0.206 | 0.33 | 1.2714E-70 |
| SDHD | 6.5077E-75 | 5.85202067 | 0.233 | 0.373 | 1.4805E-70 |
| TOMM22 | 6.576E-75 | 22.2053183 | 0.248 | 0.404 | 1.496E-70 |
| TTC3 | 8.0852E-75 | 21.8714447 | 0.267 | 0.425 | 1.8394E-70 |
| G3BP2 | 8.552E-75 | 19.8821784 | 0.288 | 0.434 | 1.9456E-70 |
| TMED5 | 9.7771E-75 | 18.5461187 | 0.332 | 0.475 | 2.2243E-70 |
| EIF3E | 1.3487E-74 | 27.4466039 | 0.289 | 0.451 | 3.0683E-70 |
| STAT1 | 2.126E-74 | 32.4881821 | 0.479 | 0.592 | 4.8366E-70 |
| HIST1H2BK | 2.7536E-74 | 12.2827889 | 0.137 | 0.257 | 6.2644E-70 |
| DESI1 | 5.8003E-74 | 1.09084824 | 0.16 | 0.281 | 1.3196E-69 |
| ERLEC1 | 7.4389E-74 | 35.1356059 | 0.19 | 0.326 | 1.6923E-69 |
| PEPD | 9.0427E-74 | 2.08170645 | 0.29 | 0.426 | 2.0572E-69 |
| POLR2G | 1.003E-73 | 7.56570635 | 0.227 | 0.363 | 2.2818E-69 |
| HNRNPAB | 1.0182E-73 | 40.7570589 | 0.26 | 0.412 | 2.3164E-69 |
| NIPBL | 1.5716E-73 | 1.83797499 | 0.273 | 0.41 | 3.5754E-69 |
| TMEM208 | 2.1129E-73 | 30.0640751 | 0.245 | 0.393 | 4.8069E-69 |
| SPAG7 | 2.3723E-73 | 5.28547742 | 0.216 | 0.352 | 5.3971E-69 |
| ENSA | 2.9524E-73 | 0.62236113 | 0.24 | 0.374 | 6.7168E-69 |
| SELENOK | 3.406E-73 | 40.8589963 | 0.213 | 0.35 | 7.7486E-69 |
| RRP7A | 5.8149E-73 | 7.63089751 | 0.27 | 0.416 | 1.3229E-68 |
| CMC1 | 6.6452E-73 | 1.55723123 | 0.149 | 0.266 | 1.5118E-68 |
| MBD5 | 8.7013E-73 | 0.31263425 | 0.197 | 0.316 | 1.9795E-68 |
| SUGT1 | 9.2476E-73 | 5.72079562 | 0.241 | 0.383 | 2.1038E-68 |
| PSMG2 | 9.3564E-73 | 5.40758979 | 0.221 | 0.362 | 2.1286E-68 |
| VDAC3 | 1.073E-72 | 2.17056078 | 0.222 | 0.358 | 2.4411E-68 |
| DDX17 | 1.1254E-72 | 23.7528275 | 0.356 | 0.502 | 2.5604E-68 |
| SREK1IP1 | 1.212E-72 | 1.20102471 | 0.143 | 0.258 | 2.7572E-68 |
| EIF3M | 1.2692E-72 | 8.30238931 | 0.3 | 0.461 | 2.8875E-68 |
| TCF12 | 1.3741E-72 | 13.3904029 | 0.273 | 0.408 | 3.1262E-68 |
| HNRNPK | 2.1348E-72 | 32.6942747 | 0.467 | 0.628 | 4.8566E-68 |
| APPL1 | 2.529E-72 | 2.9873625 | 0.176 | 0.3 | 5.7534E-68 |
| BUD31 | 2.8722E-72 | 8.21901643 | 0.23 | 0.371 | 6.5341E-68 |
| EIF2AK2 | 3.2051E-72 | 6.7000988 | 0.304 | 0.433 | 7.2915E-68 |
| CNBP | 3.2785E-72 | 58.00432 | 0.483 | 0.64 | 7.4585E-68 |
| HSPA1A | 3.9746E-72 | 338.339465 | 0.194 | 0.312 | 9.0422E-68 |
| STARD3NL | 4.4757E-72 | 0.75303011 | 0.171 | 0.293 | 1.0182E-67 |
| PSMB4 | 4.6175E-72 | 10.4165502 | 0.229 | 0.368 | 1.0505E-67 |
| PJA2 | 5.1342E-72 | 9.13558024 | 0.272 | 0.403 | 1.168E-67 |
| SNRPE | 5.1877E-72 | 37.0623853 | 0.25 | 0.404 | 1.1802E-67 |

| | | | | | |
|---|---|---|---|---|---|
| PPP2R1A | 7.2605E-72 | 25.2753725 | 0.299 | 0.462 | 1.6518E-67 |
| EIF4E2 | 7.9989E-72 | 6.40199578 | 0.275 | 0.419 | 1.8197E-67 |
| NRBF2 | 9.6506E-72 | 11.8153847 | 0.192 | 0.318 | 2.1955E-67 |
| MDM2 | 1.7432E-71 | 28.0608914 | 0.245 | 0.373 | 3.9657E-67 |
| UBE2J1 | 2.7026E-71 | 11.3547298 | 0.251 | 0.39 | 6.1483E-67 |
| HMGA1 | 3.2666E-71 | 13.6939875 | 0.372 | 0.532 | 7.4316E-67 |
| RSF1 | 8.3914E-71 | 3.82408042 | 0.247 | 0.386 | 1.909E-66 |
| ERCC1 | 9.114E-71 | 9.75090389 | 0.215 | 0.354 | 2.0734E-66 |
| PDIA3 | 1.0133E-70 | 72.8890413 | 0.488 | 0.653 | 2.3052E-66 |
| DYNC1H1 | 1.053E-70 | 14.2558165 | 0.335 | 0.485 | 2.3956E-66 |
| PIN1 | 1.6715E-70 | 1.25647326 | 0.183 | 0.305 | 3.8027E-66 |
| GGNBP2 | 1.8419E-70 | 1.05624928 | 0.232 | 0.361 | 4.1904E-66 |
| NANS | 2.7428E-70 | 11.4662048 | 0.221 | 0.364 | 6.24E-66 |
| NUP62 | 2.8845E-70 | 0.85099623 | 0.154 | 0.269 | 6.5623E-66 |
| XAF1 | 2.9237E-70 | 3.24282075 | 0.147 | 0.255 | 6.6514E-66 |
| PDIA6 | 3.0926E-70 | 105.030144 | 0.394 | 0.56 | 7.0357E-66 |
| LY6E | 3.1955E-70 | 45.4724382 | 0.506 | 0.607 | 7.2698E-66 |
| POLR2E | 3.5421E-70 | 11.7520364 | 0.249 | 0.404 | 8.0582E-66 |
| EIF5B | 3.755E-70 | 14.2880423 | 0.288 | 0.444 | 8.5427E-66 |
| SNTB1 | 5.3175E-70 | 0.56437224 | 0.177 | 0.289 | 1.2097E-65 |
| TRAPPC3 | 6.277E-70 | 6.29289979 | 0.246 | 0.387 | 1.428E-65 |
| PPP2R3C | 1.164E-69 | 1.83622248 | 0.151 | 0.263 | 2.648E-65 |
| TPD52L2 | 1.7201E-69 | 15.0952135 | 0.26 | 0.391 | 3.9132E-65 |
| NDUFS4 | 2.7816E-69 | 4.53364538 | 0.209 | 0.339 | 6.3282E-65 |
| HSD17B12 | 3.1736E-69 | 0.44753147 | 0.179 | 0.3 | 7.2199E-65 |
| GDI2 | 4.9104E-69 | 37.0004492 | 0.419 | 0.569 | 1.1171E-64 |
| RELB | 6.6035E-69 | 5.18486248 | 0.199 | 0.325 | 1.5023E-64 |
| ASH1L | 1.0262E-68 | 2.08584367 | 0.264 | 0.391 | 2.3345E-64 |
| TPR | 1.321E-68 | 15.3152936 | 0.314 | 0.468 | 3.0053E-64 |
| SZRD1 | 1.4891E-68 | 3.38434822 | 0.229 | 0.361 | 3.3876E-64 |
| DDX24 | 1.5702E-68 | 53.8112393 | 0.318 | 0.471 | 3.5722E-64 |
| MRPS25 | 1.8697E-68 | 4.22471046 | 0.198 | 0.325 | 4.2536E-64 |
| KMT2E | 1.9466E-68 | 12.7287916 | 0.27 | 0.402 | 4.4285E-64 |
| SPCS2 | 2.6178E-68 | 3.90126741 | 0.336 | 0.491 | 5.9554E-64 |
| CCT8 | 3.2955E-68 | 12.5133867 | 0.314 | 0.473 | 7.4972E-64 |
| UBE2E1 | 4.742E-68 | 1.51916419 | 0.313 | 0.457 | 1.0788E-63 |
| FAM129B | 5.5733E-68 | 8.30218428 | 0.19 | 0.306 | 1.2679E-63 |
| LAMTOR3 | 7.6305E-68 | 3.57602804 | 0.163 | 0.279 | 1.7359E-63 |
| PARP4 | 8.6403E-68 | 0.58703841 | 0.194 | 0.307 | 1.9657E-63 |
| CWC15 | 1.5872E-67 | 7.72832246 | 0.216 | 0.345 | 3.6108E-63 |
| TSPO | 1.7192E-67 | 10.95406 | 0.213 | 0.339 | 3.9111E-63 |
| RNASEH2C | 1.8669E-67 | 2.77779541 | 0.182 | 0.304 | 4.2471E-63 |
| ARL2BP | 1.8825E-67 | 2.69528027 | 0.179 | 0.301 | 4.2826E-63 |

| | | | | | |
|---|---|---|---|---|---|
| ILK | 1.953E-67 | 3.72649156 | 0.233 | 0.365 | 4.4431E-63 |
| OGFR | 2.2707E-67 | 10.8187719 | 0.183 | 0.306 | 5.1658E-63 |
| CTNNA1 | 3.1855E-67 | 7.3432086 | 0.319 | 0.466 | 7.2471E-63 |
| C14orf119 | 3.9293E-67 | 2.50346987 | 0.224 | 0.358 | 8.9392E-63 |
| PPP4R2 | 4.4591E-67 | 3.80360076 | 0.235 | 0.363 | 1.0144E-62 |
| ADK | 4.6278E-67 | 0.47874525 | 0.237 | 0.36 | 1.0528E-62 |
| SH3GLB1 | 5.6935E-67 | 25.0198408 | 0.313 | 0.459 | 1.2953E-62 |
| PNISR | 6.1765E-67 | 5.61636251 | 0.286 | 0.428 | 1.4052E-62 |
| ANKRD17 | 7.6962E-67 | 2.6494978 | 0.295 | 0.43 | 1.7509E-62 |
| IRF1 | 1.3742E-66 | 10.8477572 | 0.201 | 0.32 | 3.1264E-62 |
| IRF7 | 1.403E-66 | 5.19428645 | 0.187 | 0.301 | 3.1919E-62 |
| PHF11 | 1.9888E-66 | 1.19726836 | 0.147 | 0.258 | 4.5244E-62 |
| ARID4B | 3.9449E-66 | 5.37406169 | 0.24 | 0.369 | 8.9746E-62 |
| PHIP | 4.4163E-66 | 0.46555276 | 0.241 | 0.362 | 1.0047E-61 |
| TUSC2 | 5.0985E-66 | 2.03804636 | 0.145 | 0.258 | 1.1599E-61 |
| SNX5 | 5.1127E-66 | 2.0685407 | 0.229 | 0.351 | 1.1631E-61 |
| SEPT11 | 6.9906E-66 | 3.51189139 | 0.24 | 0.369 | 1.5904E-61 |
| GTF2F1 | 2.0665E-65 | 11.0262784 | 0.241 | 0.372 | 4.7013E-61 |
| DARS | 2.0699E-65 | 5.16766114 | 0.258 | 0.395 | 4.7091E-61 |
| VMP1 | 2.1039E-65 | 6.30030689 | 0.265 | 0.405 | 4.7864E-61 |
| GCC2 | 2.8971E-65 | 0.82314434 | 0.172 | 0.288 | 6.5909E-61 |
| PPIF | 2.9936E-65 | 4.87646646 | 0.276 | 0.403 | 6.8104E-61 |
| ZNF207 | 3.1139E-65 | 7.28318197 | 0.293 | 0.431 | 7.084E-61 |
| AFG3L2 | 5.8882E-65 | 3.0004673 | 0.194 | 0.316 | 1.3396E-60 |
| UBE2V1 | 7.3055E-65 | 11.8268315 | 0.282 | 0.424 | 1.662E-60 |
| PSMD11 | 8.7806E-65 | 24.3620997 | 0.274 | 0.418 | 1.9976E-60 |
| EMC7 | 1.1448E-64 | 17.8478334 | 0.249 | 0.393 | 2.6045E-60 |
| LSM7 | 1.3432E-64 | 29.0604044 | 0.26 | 0.41 | 3.0557E-60 |
| AKR1B1 | 1.5782E-64 | 22.389317 | 0.237 | 0.369 | 3.5904E-60 |
| PHF21A | 2.0282E-64 | 0.94312614 | 0.145 | 0.25 | 4.6141E-60 |
| SF1 | 2.142E-64 | 8.44446536 | 0.317 | 0.45 | 4.873E-60 |
| POLR2F | 2.3786E-64 | 34.257984 | 0.231 | 0.376 | 5.4113E-60 |
| PLEC | 2.9535E-64 | 20.9466158 | 0.172 | 0.283 | 6.7191E-60 |
| CALU | 4.2627E-64 | 45.173366 | 0.33 | 0.492 | 9.6977E-60 |
| MX1 | 4.9003E-64 | 12.2905084 | 0.475 | 0.555 | 1.1148E-59 |
| PSMD2 | 6.0989E-64 | 18.4766646 | 0.274 | 0.422 | 1.3875E-59 |
| VPS13B | 8.1297E-64 | 0.28161114 | 0.249 | 0.361 | 1.8495E-59 |
| EIF3F | 1.0521E-63 | 18.5261579 | 0.324 | 0.477 | 2.3935E-59 |
| ANXA7 | 1.4006E-63 | 10.1087742 | 0.263 | 0.394 | 3.1863E-59 |
| ATG3 | 1.9138E-63 | 3.87135719 | 0.218 | 0.345 | 4.3539E-59 |
| IGBP1 | 2.5758E-63 | 1.51678997 | 0.165 | 0.278 | 5.86E-59 |
| CNIH1 | 3.0846E-63 | 46.58397 | 0.245 | 0.394 | 7.0174E-59 |
| FDFT1 | 3.4888E-63 | 18.4939189 | 0.208 | 0.328 | 7.9371E-59 |

| | | | | | |
|---|---|---|---|---|---|
| MAP2K1 | 4.8515E-63 | 0.63414855 | 0.173 | 0.285 | 1.1037E-58 |
| EMC3 | 5.761E-63 | 11.5754217 | 0.209 | 0.337 | 1.3106E-58 |
| KDM2A | 1.081E-62 | 1.78922887 | 0.217 | 0.333 | 2.4594E-58 |
| HSPA9 | 1.5288E-62 | 42.7592035 | 0.359 | 0.516 | 3.4781E-58 |
| CSNK2A1 | 2.062E-62 | 4.81170053 | 0.246 | 0.378 | 4.691E-58 |
| MRPL33 | 2.318E-62 | 0.99381165 | 0.15 | 0.258 | 5.2733E-58 |
| UPF3A | 2.872E-62 | 2.57826435 | 0.181 | 0.3 | 6.5339E-58 |
| DAP3 | 4.7316E-62 | 10.0777368 | 0.244 | 0.379 | 1.0764E-57 |
| IDH3G | 5.3247E-62 | 0.33831246 | 0.18 | 0.292 | 1.2114E-57 |
| DNM1L | 6.6682E-62 | 0.73503184 | 0.171 | 0.279 | 1.517E-57 |
| NAXE | 6.8587E-62 | 15.0388628 | 0.173 | 0.285 | 1.5604E-57 |
| YY1 | 7.0115E-62 | 11.1897523 | 0.213 | 0.338 | 1.5951E-57 |
| MRPL21 | 9.3302E-62 | 19.4644713 | 0.216 | 0.349 | 2.1226E-57 |
| BPTF | 1.1846E-61 | 3.83363633 | 0.226 | 0.346 | 2.6949E-57 |
| EIF1AX | 2.1804E-61 | 20.2184879 | 0.239 | 0.379 | 4.9604E-57 |
| COX16 | 2.3524E-61 | 7.40421892 | 0.154 | 0.267 | 5.3517E-57 |
| CCND1 | 2.3893E-61 | 56.3503797 | 0.249 | 0.385 | 5.4357E-57 |
| UBE3A | 2.5417E-61 | 9.96852525 | 0.252 | 0.385 | 5.7825E-57 |
| BCAP29 | 2.5466E-61 | 4.8893711 | 0.185 | 0.3 | 5.7934E-57 |
| CCT5 | 3.6501E-61 | 35.3395597 | 0.37 | 0.52 | 8.304E-57 |
| SUB1 | 5.1281E-61 | 70.175267 | 0.4 | 0.556 | 1.1666E-56 |
| ZC3H13 | 5.4851E-61 | 3.87899016 | 0.237 | 0.361 | 1.2479E-56 |
| FAM50A | 6.3295E-61 | 1.92800269 | 0.269 | 0.395 | 1.44E-56 |
| RSL1D1 | 6.3433E-61 | 31.9816763 | 0.318 | 0.47 | 1.4431E-56 |
| UBR4 | 1.0477E-60 | 0.45102487 | 0.241 | 0.356 | 2.3836E-56 |
| PCNX1 | 1.5357E-60 | 5.04274183 | 0.181 | 0.286 | 3.4937E-56 |
| SSH1 | 1.606E-60 | 3.26799255 | 0.172 | 0.278 | 3.6538E-56 |
| TAGLN2 | 1.6429E-60 | 116.16449 | 0.438 | 0.581 | 3.7375E-56 |
| TMEM18 | 1.7685E-60 | 1.77037766 | 0.147 | 0.257 | 4.0234E-56 |
| OSBPL9 | 2.8636E-60 | 0.35573335 | 0.176 | 0.283 | 6.5148E-56 |
| CLIP1 | 3.3659E-60 | 1.00248185 | 0.186 | 0.297 | 7.6574E-56 |
| SRSF10 | 6.2817E-60 | 7.26052946 | 0.286 | 0.423 | 1.4291E-55 |
| DDX18 | 1.0055E-59 | 16.2531641 | 0.241 | 0.375 | 2.2876E-55 |
| CLPP | 1.0576E-59 | 2.90851712 | 0.196 | 0.317 | 2.406E-55 |
| CSNK1D | 1.705E-59 | 2.29963983 | 0.187 | 0.304 | 3.879E-55 |
| EZR | 1.8184E-59 | 19.9563158 | 0.171 | 0.273 | 4.137E-55 |
| TXNL4A | 1.9962E-59 | 7.27051106 | 0.175 | 0.293 | 4.5414E-55 |
| MRPL4 | 3.0004E-59 | 5.93188213 | 0.219 | 0.348 | 6.8259E-55 |
| SRP9 | 3.2139E-59 | 6.58842612 | 0.263 | 0.401 | 7.3116E-55 |
| MAPKAP1 | 3.4636E-59 | 0.63644349 | 0.194 | 0.306 | 7.8796E-55 |
| NHP2 | 4.3763E-59 | 23.2897534 | 0.26 | 0.405 | 9.956E-55 |
| XIAP | 4.8192E-59 | 1.11936603 | 0.178 | 0.285 | 1.0964E-54 |
| HNRNPF | 4.9928E-59 | 17.2070491 | 0.399 | 0.541 | 1.1359E-54 |

| | | | | | |
|---|---|---|---|---|---|
| MRPL43 | 9.9587E-59 | 7.80538833 | 0.181 | 0.301 | 2.2656E-54 |
| WAC | 1.6064E-58 | 5.37885542 | 0.201 | 0.31 | 3.6547E-54 |
| GLB1 | 1.6276E-58 | 1.6837634 | 0.19 | 0.304 | 3.7029E-54 |
| FGFR1OP2 | 2.2822E-58 | 0.33407545 | 0.165 | 0.272 | 5.192E-54 |
| PHB2 | 2.3034E-58 | 19.271821 | 0.286 | 0.434 | 5.2401E-54 |
| TRAPPC4 | 2.349E-58 | 2.03036797 | 0.183 | 0.297 | 5.344E-54 |
| SMAP1 | 3.3656E-58 | 1.68505342 | 0.207 | 0.323 | 7.6567E-54 |
| CDC73 | 6.1304E-58 | 0.264056 | 0.208 | 0.325 | 1.3947E-53 |
| FAAP20 | 7.5774E-58 | 2.89117393 | 0.154 | 0.26 | 1.7239E-53 |
| GYG1 | 7.6465E-58 | 5.22572402 | 0.18 | 0.288 | 1.7396E-53 |
| PAPOLA | 1.5552E-57 | 9.11045232 | 0.276 | 0.406 | 3.5381E-53 |
| TXNL1 | 2.2718E-57 | 8.04334869 | 0.247 | 0.385 | 5.1684E-53 |
| DNAJC3 | 3.1021E-57 | 100.894082 | 0.281 | 0.425 | 7.0572E-53 |
| MT1X | 3.1154E-57 | 100.291209 | 0.169 | 0.273 | 7.0875E-53 |
| MRPS18B | 6.6385E-57 | 4.16179913 | 0.182 | 0.294 | 1.5103E-52 |
| BNIP2 | 8.7999E-57 | 1.73183749 | 0.206 | 0.316 | 2.002E-52 |
| SEPT2 | 1.4297E-56 | 23.6528762 | 0.313 | 0.453 | 3.2527E-52 |
| TXNDC12 | 2.1052E-56 | 5.67958878 | 0.214 | 0.333 | 4.7893E-52 |
| AP3S1 | 2.181E-56 | 6.92898962 | 0.228 | 0.354 | 4.9618E-52 |
| RAD23B | 2.2561E-56 | 25.8727072 | 0.222 | 0.341 | 5.1326E-52 |
| TAF15 | 3.3233E-56 | 25.0890667 | 0.316 | 0.454 | 7.5605E-52 |
| ECHS1 | 3.7017E-56 | 5.36451752 | 0.202 | 0.325 | 8.4215E-52 |
| LUC7L3 | 3.957E-56 | 6.220169 | 0.255 | 0.381 | 9.0023E-52 |
| ARGLU1 | 8.4176E-56 | 10.8908941 | 0.284 | 0.415 | 1.915E-51 |
| SMIM12 | 8.5623E-56 | 0.2534964 | 0.169 | 0.273 | 1.9479E-51 |
| TM9SF4 | 9.1467E-56 | 0.30053565 | 0.154 | 0.254 | 2.0809E-51 |
| DCTN2 | 1.2972E-55 | 5.32555663 | 0.235 | 0.357 | 2.951E-51 |
| DGUOK | 1.4573E-55 | 5.10007089 | 0.23 | 0.351 | 3.3154E-51 |
| SMIM15 | 1.6311E-55 | 0.47487529 | 0.17 | 0.276 | 3.7107E-51 |
| HIPK3 | 1.6362E-55 | 0.62687123 | 0.17 | 0.271 | 3.7224E-51 |
| ISG15 | 1.7534E-55 | 90.3875205 | 0.508 | 0.563 | 3.9889E-51 |
| C1D | 2.1351E-55 | 1.58469184 | 0.188 | 0.297 | 4.8573E-51 |
| ADH5 | 2.9806E-55 | 20.939588 | 0.265 | 0.396 | 6.7808E-51 |
| PHLDA1 | 3.3871E-55 | 75.3144808 | 0.395 | 0.49 | 7.7057E-51 |
| THRAP3 | 3.9657E-55 | 18.7002668 | 0.341 | 0.483 | 9.0219E-51 |
| PSMC1 | 5.3125E-55 | 1.78594241 | 0.234 | 0.356 | 1.2086E-50 |
| ERGIC3 | 6.1929E-55 | 17.3396858 | 0.257 | 0.39 | 1.4089E-50 |
| PDLIM7 | 6.3613E-55 | 14.8818158 | 0.19 | 0.304 | 1.4472E-50 |
| RNF115 | 6.9727E-55 | 1.79575246 | 0.178 | 0.283 | 1.5863E-50 |
| HSPA4 | 8.4915E-55 | 19.6570399 | 0.306 | 0.44 | 1.9318E-50 |
| MAPRE1 | 1.1835E-54 | 4.10777233 | 0.234 | 0.349 | 2.6925E-50 |
| NPEPPS | 1.298E-54 | 1.49593679 | 0.22 | 0.332 | 2.953E-50 |
| HNRNPD | 1.4325E-54 | 12.0703928 | 0.245 | 0.373 | 3.2589E-50 |

| | | | | | |
|---|---|---|---|---|---|
| UFM1 | 1.8339E-54 | 22.5421497 | 0.258 | 0.384 | 4.1722E-50 |
| TRAPPC5 | 1.9457E-54 | 5.21231921 | 0.169 | 0.277 | 4.4264E-50 |
| GLG1 | 3.4421E-54 | 8.86243376 | 0.234 | 0.356 | 7.8308E-50 |
| SF3B1 | 3.8843E-54 | 16.3369566 | 0.333 | 0.47 | 8.8368E-50 |
| TUBB6 | 4.0608E-54 | 16.5210301 | 0.248 | 0.376 | 9.2384E-50 |
| MIER1 | 4.4155E-54 | 1.81587392 | 0.214 | 0.324 | 1.0045E-49 |
| NDUFV1 | 4.6754E-54 | 2.53374157 | 0.228 | 0.345 | 1.0637E-49 |
| TIMMDC1 | 1.1376E-53 | 7.13465676 | 0.172 | 0.278 | 2.5881E-49 |
| YPEL5 | 1.2381E-53 | 0.88652317 | 0.154 | 0.253 | 2.8167E-49 |
| SETX | 1.347E-53 | 7.69621336 | 0.227 | 0.338 | 3.0644E-49 |
| LSM3 | 1.4945E-53 | 13.388169 | 0.208 | 0.322 | 3.4E-49 |
| WIPI2 | 1.6533E-53 | 2.40548651 | 0.165 | 0.272 | 3.7612E-49 |
| PSMD12 | 1.7764E-53 | 11.2914846 | 0.26 | 0.387 | 4.0414E-49 |
| PSMC3 | 1.9376E-53 | 6.56673103 | 0.248 | 0.376 | 4.4081E-49 |
| SRSF7 | 2.1225E-53 | 51.9181784 | 0.305 | 0.448 | 4.8286E-49 |
| MTX1 | 2.5565E-53 | 2.53716758 | 0.154 | 0.259 | 5.816E-49 |
| COA4 | 3.3119E-53 | 23.2326577 | 0.233 | 0.363 | 7.5346E-49 |
| ILF2 | 4.5473E-53 | 38.2570537 | 0.279 | 0.421 | 1.0345E-48 |
| EWSR1 | 5.4453E-53 | 44.0590185 | 0.359 | 0.499 | 1.2388E-48 |
| GOLGA7 | 7.7372E-53 | 4.12567706 | 0.201 | 0.312 | 1.7602E-48 |
| MLEC | 7.998E-53 | 31.4743071 | 0.247 | 0.377 | 1.8195E-48 |
| PRKDC | 8.609E-53 | 9.87063102 | 0.29 | 0.426 | 1.9585E-48 |
| RABAC1 | 9.3224E-53 | 57.0115532 | 0.273 | 0.407 | 2.1209E-48 |
| ZFR | 1.0315E-52 | 9.11213736 | 0.27 | 0.397 | 2.3466E-48 |
| NSRP1 | 1.6035E-52 | 4.81817312 | 0.211 | 0.318 | 3.6478E-48 |
| SAT2 | 1.7902E-52 | 1.26837541 | 0.175 | 0.28 | 4.0726E-48 |
| PTGR1 | 1.9515E-52 | 104.622318 | 0.178 | 0.283 | 4.4398E-48 |
| CCPG1 | 2.5388E-52 | 6.52627342 | 0.255 | 0.371 | 5.7757E-48 |
| GSK3B | 2.8426E-52 | 0.48046654 | 0.264 | 0.375 | 6.467E-48 |
| SNX17 | 3.0934E-52 | 3.21302459 | 0.213 | 0.324 | 7.0376E-48 |
| ARID1B | 3.1173E-52 | 1.09238717 | 0.198 | 0.299 | 7.0918E-48 |
| UBE2N | 3.8233E-52 | 4.90384413 | 0.249 | 0.37 | 8.698E-48 |
| MAGOH | 8.6647E-52 | 5.80643268 | 0.229 | 0.348 | 1.9712E-47 |
| CREM | 8.6649E-52 | 11.8836876 | 0.161 | 0.26 | 1.9713E-47 |
| SNRNP70 | 8.7123E-52 | 13.1971366 | 0.255 | 0.384 | 1.982E-47 |
| TGIF1 | 1.0521E-51 | 16.3382969 | 0.192 | 0.301 | 2.3936E-47 |
| C11orf98 | 1.2414E-51 | 9.59983604 | 0.246 | 0.367 | 2.8242E-47 |
| HNRNPA0 | 1.5361E-51 | 18.4740025 | 0.278 | 0.41 | 3.4947E-47 |
| ELOVL1 | 1.5686E-51 | 6.81169504 | 0.194 | 0.308 | 3.5687E-47 |
| TERF2IP | 1.6557E-51 | 13.5365249 | 0.23 | 0.35 | 3.7667E-47 |
| OASL | 1.6664E-51 | 2.19282091 | 0.168 | 0.254 | 3.791E-47 |
| FLII | 1.7035E-51 | 4.45449601 | 0.229 | 0.336 | 3.8754E-47 |
| SNRPF | 1.8245E-51 | 12.9732009 | 0.222 | 0.35 | 4.1507E-47 |

| | | | | | |
|---|---|---|---|---|---|
| C8orf59 | 1.8551E-51 | 9.07529604 | 0.199 | 0.311 | 4.2203E-47 |
| SEC22B | 2.3577E-51 | 6.80231545 | 0.221 | 0.329 | 5.3637E-47 |
| SERINC1 | 2.4377E-51 | 7.30561424 | 0.253 | 0.37 | 5.5458E-47 |
| SFPQ | 2.7831E-51 | 34.5488068 | 0.266 | 0.389 | 6.3316E-47 |
| SRSF4 | 2.9062E-51 | 6.62085387 | 0.3 | 0.425 | 6.6115E-47 |
| PRPF6 | 4.9598E-51 | 3.14412584 | 0.232 | 0.344 | 1.1283E-46 |
| PABPN1 | 5.5037E-51 | 3.67532215 | 0.183 | 0.29 | 1.2521E-46 |
| RANBP1 | 6.5409E-51 | 17.7583624 | 0.243 | 0.37 | 1.4881E-46 |
| LRP10 | 6.9267E-51 | 0.99960671 | 0.201 | 0.309 | 1.5758E-46 |
| CCDC90B | 1.1979E-50 | 1.43715023 | 0.168 | 0.27 | 2.7251E-46 |
| SEPT9 | 1.2068E-50 | 7.44188072 | 0.271 | 0.38 | 2.7455E-46 |
| RNMT | 1.5231E-50 | 1.78972451 | 0.184 | 0.29 | 3.4651E-46 |
| KRAS | 1.5493E-50 | 5.90747705 | 0.17 | 0.272 | 3.5246E-46 |
| DSTN | 1.5804E-50 | 28.737315 | 0.247 | 0.378 | 3.5955E-46 |
| NSA2 | 2.217E-50 | 6.59172075 | 0.24 | 0.361 | 5.0438E-46 |
| NDUFA10 | 2.4048E-50 | 4.51709214 | 0.215 | 0.327 | 5.4708E-46 |
| SKIL | 2.6505E-50 | 5.8622768 | 0.166 | 0.262 | 6.0298E-46 |
| MRPL36 | 2.652E-50 | 9.14905478 | 0.185 | 0.296 | 6.0334E-46 |
| GNAI3 | 3.3386E-50 | 3.71940168 | 0.25 | 0.364 | 7.5954E-46 |
| PUM1 | 3.5031E-50 | 4.62062345 | 0.241 | 0.356 | 7.9696E-46 |
| RBM6 | 5.3104E-50 | 0.58179584 | 0.253 | 0.36 | 1.2081E-45 |
| SEC13 | 1.0793E-49 | 41.448426 | 0.286 | 0.419 | 2.4555E-45 |
| PSMC2 | 1.1067E-49 | 26.1792596 | 0.235 | 0.352 | 2.5177E-45 |
| TSPAN4 | 1.1159E-49 | 11.7412618 | 0.168 | 0.27 | 2.5387E-45 |
| CCT7 | 1.1342E-49 | 34.0848629 | 0.315 | 0.456 | 2.5804E-45 |
| IBTK | 1.2978E-49 | 3.34136068 | 0.189 | 0.293 | 2.9525E-45 |
| PPP1R11 | 1.3815E-49 | 1.77671503 | 0.197 | 0.3 | 3.1428E-45 |
| UBAP1 | 3.868E-49 | 1.41327853 | 0.187 | 0.284 | 8.7998E-45 |
| IMP3 | 3.9227E-49 | 10.9365765 | 0.197 | 0.31 | 8.924E-45 |
| NAA10 | 4.3432E-49 | 12.2200006 | 0.21 | 0.329 | 9.8808E-45 |
| GTF3A | 4.3952E-49 | 4.90535304 | 0.267 | 0.386 | 9.9991E-45 |
| SYNCRIP | 4.6104E-49 | 25.3673904 | 0.312 | 0.451 | 1.0489E-44 |
| POLR1D | 6.0989E-49 | 11.0039812 | 0.225 | 0.338 | 1.3875E-44 |
| EIF4G1 | 6.5069E-49 | 35.1923339 | 0.388 | 0.523 | 1.4803E-44 |
| STUB1 | 6.6991E-49 | 4.65792531 | 0.183 | 0.293 | 1.524E-44 |
| GLRX3 | 6.9243E-49 | 5.36881824 | 0.245 | 0.368 | 1.5753E-44 |
| ZMAT2 | 8.3667E-49 | 4.15670823 | 0.23 | 0.339 | 1.9034E-44 |
| MARK3 | 8.4454E-49 | 4.15439454 | 0.178 | 0.273 | 1.9213E-44 |
| AC138123.1 | 1.2798E-48 | 6.26479685 | 0.172 | 0.273 | 2.9115E-44 |
| STAT3 | 2.1733E-48 | 38.8631972 | 0.34 | 0.471 | 4.9442E-44 |
| RPL7L1 | 2.4303E-48 | 20.1965403 | 0.243 | 0.366 | 5.529E-44 |
| NONO | 2.8879E-48 | 87.2485003 | 0.342 | 0.475 | 6.57E-44 |
| MVP | 4.1049E-48 | 13.8433159 | 0.196 | 0.303 | 9.3387E-44 |

| | | | | | |
|---|---|---|---|---|---|
| SCAMP3 | 5.3932E-48 | 3.74680441 | 0.221 | 0.335 | 1.2269E-43 |
| OAS3 | 7.5675E-48 | 9.42727927 | 0.239 | 0.332 | 1.7216E-43 |
| TRPS1 | 8.2968E-48 | 9.39563509 | 0.261 | 0.373 | 1.8875E-43 |
| ETF1 | 1.1016E-47 | 6.63629559 | 0.247 | 0.356 | 2.5062E-43 |
| HDAC8 | 1.1459E-47 | 0.95467261 | 0.192 | 0.288 | 2.6069E-43 |
| KDM5A | 1.1974E-47 | 0.99911956 | 0.217 | 0.32 | 2.7241E-43 |
| CTNNB1 | 1.2288E-47 | 19.7778972 | 0.322 | 0.44 | 2.7955E-43 |
| WDFY1 | 1.6769E-47 | 1.51312664 | 0.16 | 0.252 | 3.8149E-43 |
| MAPK1IP1L | 1.9971E-47 | 6.25278535 | 0.25 | 0.36 | 4.5433E-43 |
| UGCG | 2.1096E-47 | 8.89196872 | 0.161 | 0.254 | 4.7992E-43 |
| RNF5 | 2.2281E-47 | 3.22047966 | 0.172 | 0.27 | 5.069E-43 |
| POP4 | 3.6566E-47 | 3.22455686 | 0.157 | 0.252 | 8.3188E-43 |
| BRD2 | 3.7516E-47 | 6.07879534 | 0.278 | 0.399 | 8.5348E-43 |
| EIF4A1 | 3.9212E-47 | 6.33502185 | 0.218 | 0.328 | 8.9208E-43 |
| FNBP4 | 3.9595E-47 | 6.66790932 | 0.226 | 0.338 | 9.0078E-43 |
| CSRP1 | 4.1922E-47 | 31.6638574 | 0.158 | 0.251 | 9.5373E-43 |
| TDG | 4.198E-47 | 3.28618903 | 0.169 | 0.266 | 9.5504E-43 |
| EIF4E | 4.8923E-47 | 10.4940267 | 0.244 | 0.358 | 1.113E-42 |
| PSMF1 | 5.9947E-47 | 6.00674472 | 0.248 | 0.362 | 1.3638E-42 |
| STAU1 | 8.4451E-47 | 10.1590272 | 0.244 | 0.357 | 1.9213E-42 |
| COPB2 | 8.9648E-47 | 17.3891627 | 0.28 | 0.406 | 2.0395E-42 |
| TAF7 | 9.3196E-47 | 9.2356693 | 0.219 | 0.331 | 2.1202E-42 |
| ITCH | 1.045E-46 | 1.32545905 | 0.21 | 0.309 | 2.3775E-42 |
| EEA1 | 1.1862E-46 | 0.55762032 | 0.165 | 0.258 | 2.6987E-42 |
| SNF8 | 1.2673E-46 | 2.67431556 | 0.159 | 0.255 | 2.8831E-42 |
| ATRX | 1.2808E-46 | 5.56018416 | 0.25 | 0.358 | 2.9139E-42 |
| MRPS23 | 1.6714E-46 | 3.74163231 | 0.171 | 0.271 | 3.8024E-42 |
| MRPS12 | 1.6993E-46 | 15.8535206 | 0.216 | 0.33 | 3.8658E-42 |
| SNRPB2 | 2.1587E-46 | 19.3588848 | 0.228 | 0.343 | 4.911E-42 |
| LSM5 | 4.2924E-46 | 10.7205122 | 0.183 | 0.285 | 9.7652E-42 |
| REST | 4.7535E-46 | 10.6692231 | 0.16 | 0.254 | 1.0814E-41 |
| TUBGCP2 | 5.1322E-46 | 2.09002984 | 0.188 | 0.286 | 1.1676E-41 |
| YIF1A | 5.8381E-46 | 16.1475861 | 0.197 | 0.31 | 1.3282E-41 |
| SRPRA | 7.7191E-46 | 18.5118738 | 0.269 | 0.386 | 1.7561E-41 |
| ARID1A | 8.1551E-46 | 0.30643887 | 0.169 | 0.26 | 1.8553E-41 |
| HIKESHI | 9.0091E-46 | 6.43304063 | 0.157 | 0.252 | 2.0496E-41 |
| PDAP1 | 1.0057E-45 | 2.44357167 | 0.243 | 0.363 | 2.2879E-41 |
| SLC39A1 | 1.5374E-45 | 3.5263591 | 0.18 | 0.282 | 3.4977E-41 |
| STK17A | 1.5876E-45 | 8.54385672 | 0.172 | 0.268 | 3.6119E-41 |
| GRSF1 | 1.7643E-45 | 9.49824001 | 0.21 | 0.318 | 4.0137E-41 |
| LSM2 | 2.7445E-45 | 20.1006827 | 0.221 | 0.333 | 6.2438E-41 |
| METAP2 | 4.6418E-45 | 1.71914853 | 0.229 | 0.339 | 1.056E-40 |
| GSS | 4.7115E-45 | 0.60339529 | 0.164 | 0.256 | 1.0719E-40 |

| | | | | | |
|---|---|---|---|---|---|
| NCBP2 | 6.4948E-45 | 8.95113048 | 0.164 | 0.262 | 1.4776E-40 |
| IST1 | 6.639E-45 | 0.32037295 | 0.182 | 0.277 | 1.5104E-40 |
| HECTD1 | 7.0563E-45 | 2.96657262 | 0.259 | 0.371 | 1.6053E-40 |
| RSRC2 | 7.3877E-45 | 1.05730827 | 0.224 | 0.332 | 1.6807E-40 |
| GPATCH8 | 8.0647E-45 | 1.60931014 | 0.196 | 0.295 | 1.8347E-40 |
| TMED2 | 9.3144E-45 | 27.476813 | 0.274 | 0.4 | 2.119E-40 |
| SPIDR | 1.0571E-44 | 13.0262704 | 0.235 | 0.332 | 2.405E-40 |
| PSMD13 | 1.2175E-44 | 6.11622019 | 0.227 | 0.339 | 2.7697E-40 |
| NDUFC2 | 1.5903E-44 | 9.01199341 | 0.167 | 0.261 | 3.618E-40 |
| BRD4 | 1.5975E-44 | 1.34053508 | 0.22 | 0.323 | 3.6344E-40 |
| GOLGB1 | 2.0389E-44 | 9.97816695 | 0.244 | 0.351 | 4.6385E-40 |
| UBA1 | 2.0648E-44 | 2.13602302 | 0.166 | 0.258 | 4.6975E-40 |
| VPS37A | 2.2098E-44 | 3.29519637 | 0.165 | 0.254 | 5.0272E-40 |
| ANKRD11 | 2.398E-44 | 8.89263618 | 0.283 | 0.398 | 5.4554E-40 |
| TTC1 | 2.4063E-44 | 0.44946378 | 0.196 | 0.293 | 5.4743E-40 |
| HERC2 | 2.4473E-44 | 3.11099848 | 0.219 | 0.319 | 5.5677E-40 |
| FAM136A | 2.8652E-44 | 4.71808253 | 0.174 | 0.271 | 6.5184E-40 |
| COPB1 | 2.8742E-44 | 21.4007933 | 0.251 | 0.362 | 6.5388E-40 |
| S100A6 | 3.1049E-44 | 173.872297 | 0.506 | 0.588 | 7.0636E-40 |
| ZMYM2 | 3.9776E-44 | 2.28505628 | 0.172 | 0.263 | 9.049E-40 |
| EMP3 | 4.5083E-44 | 111.83641 | 0.579 | 0.655 | 1.0256E-39 |
| ABL2 | 4.7851E-44 | 29.4479936 | 0.195 | 0.288 | 1.0886E-39 |
| CTNND1 | 4.7954E-44 | 3.12066792 | 0.163 | 0.254 | 1.0909E-39 |
| MRPL13 | 5.2835E-44 | 6.96979138 | 0.193 | 0.297 | 1.202E-39 |
| PSMC6 | 5.7204E-44 | 10.6235843 | 0.216 | 0.326 | 1.3014E-39 |
| GTF2B | 7.2128E-44 | 7.72981082 | 0.214 | 0.315 | 1.6409E-39 |
| NDUFA9 | 8.7514E-44 | 3.72226857 | 0.17 | 0.268 | 1.9909E-39 |
| RAB1B | 9.8557E-44 | 2.5956124 | 0.223 | 0.326 | 2.2422E-39 |
| CCDC47 | 1.0633E-43 | 8.08248496 | 0.248 | 0.361 | 2.4189E-39 |
| DENR | 1.1999E-43 | 9.23800645 | 0.249 | 0.36 | 2.7299E-39 |
| HK1 | 1.5635E-43 | 10.4210776 | 0.214 | 0.313 | 3.557E-39 |
| WARS | 1.8504E-43 | 188.295012 | 0.268 | 0.381 | 4.2097E-39 |
| COMMD4 | 2.2742E-43 | 2.28802255 | 0.167 | 0.259 | 5.1737E-39 |
| SMIM7 | 2.4522E-43 | 8.80532252 | 0.209 | 0.316 | 5.5788E-39 |
| AP3D1 | 2.5876E-43 | 5.55022376 | 0.22 | 0.322 | 5.8869E-39 |
| UBE2Z | 2.6046E-43 | 0.69438143 | 0.164 | 0.253 | 5.9255E-39 |
| COPA | 2.6424E-43 | 6.23863365 | 0.262 | 0.37 | 6.0115E-39 |
| RPL26L1 | 3.5916E-43 | 12.9271768 | 0.176 | 0.279 | 8.171E-39 |
| PDLIM5 | 3.7599E-43 | 34.2665588 | 0.244 | 0.347 | 8.5538E-39 |
| STIP1 | 6.6634E-43 | 19.8738878 | 0.338 | 0.463 | 1.5159E-38 |
| FAM204A | 7.5809E-43 | 1.69030777 | 0.187 | 0.285 | 1.7247E-38 |
| TMCO1 | 9.0372E-43 | 15.2489456 | 0.201 | 0.307 | 2.056E-38 |
| GBE1 | 1.0565E-42 | 0.68128174 | 0.206 | 0.3 | 2.4035E-38 |

| | | | | | |
|---|---|---|---|---|---|
| LSM10 | 1.25E-42 | 4.36245194 | 0.212 | 0.316 | 2.8437E-38 |
| LMAN1 | 1.2858E-42 | 40.6090423 | 0.252 | 0.379 | 2.9252E-38 |
| EXOSC4 | 1.5711E-42 | 12.4741111 | 0.186 | 0.288 | 3.5743E-38 |
| CIAO1 | 2.0724E-42 | 3.53338437 | 0.186 | 0.284 | 4.7147E-38 |
| HUWE1 | 2.1429E-42 | 2.80004027 | 0.251 | 0.352 | 4.8751E-38 |
| CD276 | 3.6916E-42 | 2.02157686 | 0.176 | 0.271 | 8.3983E-38 |
| DTX3L | 4.0097E-42 | 7.72953351 | 0.176 | 0.26 | 9.122E-38 |
| GTF2E2 | 4.0872E-42 | 1.6252895 | 0.162 | 0.252 | 9.2983E-38 |
| CCDC12 | 4.1206E-42 | 2.28007196 | 0.173 | 0.267 | 9.3744E-38 |
| DDX6 | 4.2517E-42 | 2.28288793 | 0.211 | 0.306 | 9.6726E-38 |
| RPAIN | 4.5251E-42 | 2.80917012 | 0.172 | 0.265 | 1.0295E-37 |
| PYURF | 4.7944E-42 | 19.018193 | 0.209 | 0.315 | 1.0907E-37 |
| METTL26 | 6.0817E-42 | 6.3262942 | 0.175 | 0.273 | 1.3836E-37 |
| ASNA1 | 8.0958E-42 | 4.25115923 | 0.202 | 0.306 | 1.8418E-37 |
| MEAF6 | 1.0786E-41 | 1.85995286 | 0.188 | 0.282 | 2.4538E-37 |
| GOLGA4 | 1.1715E-41 | 15.61759 | 0.306 | 0.426 | 2.6651E-37 |
| MARCH7 | 1.6685E-41 | 8.19392023 | 0.204 | 0.296 | 3.7958E-37 |
| MRPL28 | 1.71E-41 | 6.95382992 | 0.186 | 0.285 | 3.8902E-37 |
| LSM14A | 1.8154E-41 | 8.89198489 | 0.256 | 0.365 | 4.13E-37 |
| EIF4B | 2.1682E-41 | 17.7425899 | 0.335 | 0.453 | 4.9326E-37 |
| SNW1 | 2.6909E-41 | 1.44903072 | 0.193 | 0.284 | 6.1218E-37 |
| PSMD14 | 3.0717E-41 | 3.60604347 | 0.247 | 0.354 | 6.9881E-37 |
| CBX3 | 3.9375E-41 | 7.12275699 | 0.224 | 0.33 | 8.9578E-37 |
| YME1L1 | 3.9952E-41 | 10.659093 | 0.249 | 0.355 | 9.089E-37 |
| MRPL16 | 6.3785E-41 | 2.40494963 | 0.188 | 0.282 | 1.4511E-36 |
| PPP1CC | 6.6123E-41 | 0.44356934 | 0.234 | 0.342 | 1.5043E-36 |
| RAB5A | 7.1798E-41 | 1.00858349 | 0.2 | 0.292 | 1.6334E-36 |
| SF3B4 | 7.4508E-41 | 6.49993548 | 0.233 | 0.342 | 1.6951E-36 |
| DNAJB14 | 9.1564E-41 | 0.82453261 | 0.199 | 0.29 | 2.0831E-36 |
| MCFD2 | 9.5455E-41 | 27.8995347 | 0.234 | 0.348 | 2.1716E-36 |
| MRPS15 | 1.011E-40 | 11.4479246 | 0.207 | 0.312 | 2.3001E-36 |
| USP9X | 1.8637E-40 | 2.22308806 | 0.176 | 0.264 | 4.2399E-36 |
| ATG5 | 1.9558E-40 | 1.70410781 | 0.189 | 0.281 | 4.4494E-36 |
| PPP1R15A | 2.5541E-40 | 92.0807168 | 0.355 | 0.436 | 5.8105E-36 |
| SAP30BP | 2.7358E-40 | 11.4499502 | 0.188 | 0.282 | 6.224E-36 |
| ITGAV | 2.9413E-40 | 4.93370858 | 0.213 | 0.311 | 6.6915E-36 |
| NDUFA11 | 3.1094E-40 | 9.28147875 | 0.202 | 0.304 | 7.0739E-36 |
| ASPH | 3.1361E-40 | 0.75684468 | 0.428 | 0.528 | 7.1346E-36 |
| MZT2B | 3.327E-40 | 11.8209283 | 0.177 | 0.273 | 7.5689E-36 |
| RAB11B | 3.5708E-40 | 7.2968873 | 0.194 | 0.292 | 8.1235E-36 |
| APEX1 | 4.415E-40 | 21.2327705 | 0.266 | 0.385 | 1.0044E-35 |
| YIPF3 | 6.0752E-40 | 6.77316497 | 0.189 | 0.289 | 1.3821E-35 |
| SRSF2 | 7.3695E-40 | 20.4040447 | 0.254 | 0.369 | 1.6766E-35 |

| | | | | | |
|---|---|---|---|---|---|
| CUEDC2 | 8.3643E-40 | 4.32170434 | 0.16 | 0.251 | 1.9029E-35 |
| RAD21 | 1.0809E-39 | 10.1553154 | 0.248 | 0.35 | 2.4592E-35 |
| HDAC2 | 1.4457E-39 | 10.3470882 | 0.249 | 0.358 | 3.2889E-35 |
| IFIT3 | 1.8223E-39 | 29.3965848 | 0.355 | 0.415 | 4.1458E-35 |
| PDXDC1 | 1.8736E-39 | 1.15963856 | 0.201 | 0.291 | 4.2623E-35 |
| LARP7 | 1.9222E-39 | 0.56088457 | 0.167 | 0.252 | 4.3729E-35 |
| U2SURP | 2.5638E-39 | 3.42350103 | 0.255 | 0.367 | 5.8327E-35 |
| RPN1 | 2.599E-39 | 44.9789793 | 0.315 | 0.44 | 5.9128E-35 |
| LRRFIP2 | 2.863E-39 | 1.39119521 | 0.187 | 0.276 | 6.5133E-35 |
| EPN1 | 3.7222E-39 | 2.20732095 | 0.164 | 0.253 | 8.4679E-35 |
| FAF2 | 3.8797E-39 | 7.04701787 | 0.208 | 0.31 | 8.8263E-35 |
| PSMD9 | 4.3863E-39 | 8.23672721 | 0.174 | 0.267 | 9.9789E-35 |
| SCFD1 | 4.8627E-39 | 0.51351291 | 0.178 | 0.264 | 1.1063E-34 |
| OLA1 | 5.3734E-39 | 3.86337005 | 0.278 | 0.392 | 1.2225E-34 |
| KANSL1 | 5.7036E-39 | 8.18012312 | 0.184 | 0.267 | 1.2976E-34 |
| CCAR1 | 6.6539E-39 | 6.85576452 | 0.225 | 0.328 | 1.5138E-34 |
| UFD1 | 8.9869E-39 | 4.21364408 | 0.218 | 0.319 | 2.0445E-34 |
| NARS | 1.258E-38 | 10.983077 | 0.216 | 0.315 | 2.862E-34 |
| EIF4EBP1 | 1.4034E-38 | 26.2851597 | 0.197 | 0.297 | 3.1928E-34 |
| SDF2L1 | 1.8279E-38 | 64.199984 | 0.217 | 0.316 | 4.1585E-34 |
| DERL1 | 1.9355E-38 | 4.97901094 | 0.175 | 0.264 | 4.4032E-34 |
| MAP7D1 | 2.1531E-38 | 3.21387571 | 0.179 | 0.268 | 4.8984E-34 |
| MRPL47 | 2.3525E-38 | 5.49822267 | 0.177 | 0.272 | 5.3519E-34 |
| AP1S1 | 2.8519E-38 | 13.5329634 | 0.188 | 0.285 | 6.4881E-34 |
| SYF2 | 3.9371E-38 | 1.23472048 | 0.177 | 0.264 | 8.9568E-34 |
| AFF4 | 5.9827E-38 | 3.73232869 | 0.268 | 0.368 | 1.3611E-33 |
| SMARCA5 | 6.1893E-38 | 7.75284929 | 0.254 | 0.357 | 1.4081E-33 |
| EPRS | 8.2854E-38 | 1.4024751 | 0.226 | 0.331 | 1.8849E-33 |
| SUCLG1 | 8.3827E-38 | 4.64330888 | 0.173 | 0.262 | 1.9071E-33 |
| BDP1 | 1.0181E-37 | 1.77706371 | 0.243 | 0.334 | 2.3161E-33 |
| NDUFS1 | 1.0938E-37 | 3.05597987 | 0.185 | 0.275 | 2.4883E-33 |
| TIMP3 | 1.1307E-37 | 5.07696812 | 0.273 | 0.341 | 2.5723E-33 |
| NME1 | 1.2607E-37 | 25.5220133 | 0.226 | 0.339 | 2.8681E-33 |
| DYNC1I2 | 1.4E-37 | 1.61087045 | 0.199 | 0.288 | 3.1851E-33 |
| ANO6 | 1.7937E-37 | 8.11386935 | 0.223 | 0.317 | 4.0807E-33 |
| MAF1 | 4.3277E-37 | 2.83554479 | 0.183 | 0.27 | 9.8455E-33 |
| LARP4 | 6.4928E-37 | 0.29504582 | 0.199 | 0.283 | 1.4771E-32 |
| AIMP1 | 7.6659E-37 | 5.06632742 | 0.196 | 0.29 | 1.744E-32 |
| ZC3HAV1 | 1.01E-36 | 27.6922273 | 0.199 | 0.282 | 2.2978E-32 |
| RPL36A | 1.0427E-36 | 16.1529413 | 0.275 | 0.392 | 2.3721E-32 |
| KHDRBS1 | 1.1728E-36 | 18.0550084 | 0.28 | 0.39 | 2.6681E-32 |
| BTG3 | 1.1755E-36 | 32.3674887 | 0.191 | 0.281 | 2.6742E-32 |
| CCDC124 | 1.4245E-36 | 11.5688384 | 0.217 | 0.321 | 3.2408E-32 |

| | | | | | |
|---|---|---|---|---|---|
| SRP19 | 1.4742E-36 | 43.3416604 | 0.222 | 0.322 | 3.3538E-32 |
| NUFIP2 | 1.8245E-36 | 2.18836935 | 0.212 | 0.299 | 4.1508E-32 |
| GPBP1 | 2.1849E-36 | 1.12260878 | 0.208 | 0.294 | 4.9706E-32 |
| IFI44L | 3.6596E-36 | 3.63840638 | 0.239 | 0.315 | 8.3257E-32 |
| PRPF4B | 3.797E-36 | 3.90761558 | 0.23 | 0.32 | 8.6382E-32 |
| ACIN1 | 4.2719E-36 | 6.93236454 | 0.305 | 0.412 | 9.7186E-32 |
| RTF1 | 4.2821E-36 | 2.17245928 | 0.185 | 0.273 | 9.7419E-32 |
| RBCK1 | 4.3877E-36 | 5.86484682 | 0.185 | 0.273 | 9.9821E-32 |
| PRPF8 | 4.5432E-36 | 3.64050733 | 0.271 | 0.373 | 1.0336E-31 |
| CGGBP1 | 5.1738E-36 | 1.47309736 | 0.18 | 0.262 | 1.177E-31 |
| KPNA4 | 6.9328E-36 | 3.54554491 | 0.197 | 0.281 | 1.5772E-31 |
| NUP98 | 7.3645E-36 | 5.27452295 | 0.205 | 0.291 | 1.6754E-31 |
| MRPL15 | 9.3456E-36 | 9.92614903 | 0.192 | 0.287 | 2.1261E-31 |
| DEF8 | 1.0582E-35 | 5.50048491 | 0.17 | 0.255 | 2.4074E-31 |
| INSIG1 | 1.225E-35 | 16.5484646 | 0.221 | 0.32 | 2.7868E-31 |
| GRPEL1 | 1.7348E-35 | 14.7935512 | 0.185 | 0.274 | 3.9467E-31 |
| UROD | 2.2082E-35 | 2.94672928 | 0.194 | 0.287 | 5.0236E-31 |
| ECHDC1 | 3.499E-35 | 14.4861179 | 0.215 | 0.31 | 7.9603E-31 |
| MED8 | 3.7633E-35 | 2.45743629 | 0.181 | 0.268 | 8.5614E-31 |
| SH3PXD2B | 3.9282E-35 | 21.803032 | 0.261 | 0.355 | 8.9367E-31 |
| DCTN3 | 5.3903E-35 | 3.72715472 | 0.193 | 0.282 | 1.2263E-30 |
| RBM42 | 8.9206E-35 | 11.9086532 | 0.222 | 0.319 | 2.0294E-30 |
| ADAM17 | 1.0212E-34 | 4.31983695 | 0.22 | 0.302 | 2.3233E-30 |
| NOSIP | 1.2011E-34 | 10.4247265 | 0.226 | 0.326 | 2.7326E-30 |
| FAM129A | 1.3944E-34 | 1.80379273 | 0.224 | 0.306 | 3.1722E-30 |
| ITGA5 | 1.4836E-34 | 4.61789062 | 0.204 | 0.293 | 3.3752E-30 |
| EIF2S3 | 1.8133E-34 | 11.0383111 | 0.265 | 0.366 | 4.1254E-30 |
| FARSA | 2.0221E-34 | 23.3346359 | 0.245 | 0.349 | 4.6004E-30 |
| GAPVD1 | 2.1642E-34 | 1.13670584 | 0.17 | 0.25 | 4.9235E-30 |
| THOC7 | 2.4031E-34 | 3.44918348 | 0.17 | 0.254 | 5.467E-30 |
| SNHG8 | 2.9365E-34 | 17.9359786 | 0.168 | 0.257 | 6.6806E-30 |
| SLC4A7 | 3.0599E-34 | 2.275745 | 0.202 | 0.291 | 6.9614E-30 |
| IWS1 | 3.2792E-34 | 3.85876162 | 0.191 | 0.278 | 7.4603E-30 |
| PPP6R3 | 4.9556E-34 | 2.04687847 | 0.197 | 0.277 | 1.1274E-29 |
| HNRNPR | 5.2079E-34 | 16.5408836 | 0.266 | 0.375 | 1.1848E-29 |
| MRPL18 | 5.2335E-34 | 42.5858607 | 0.199 | 0.296 | 1.1906E-29 |
| IMMP2L | 5.4388E-34 | 15.1533671 | 0.188 | 0.267 | 1.2373E-29 |
| PCNP | 5.6721E-34 | 3.23237999 | 0.21 | 0.304 | 1.2904E-29 |
| MPHOSPH8 | 6.2378E-34 | 5.92506987 | 0.215 | 0.303 | 1.4191E-29 |
| TRA2B | 7.6955E-34 | 9.02645924 | 0.262 | 0.364 | 1.7507E-29 |
| BAZ1B | 8.4406E-34 | 10.0507172 | 0.225 | 0.319 | 1.9202E-29 |
| CD151 | 1.2863E-33 | 28.3647741 | 0.224 | 0.329 | 2.9263E-29 |
| DHRS7 | 1.4473E-33 | 6.67227013 | 0.205 | 0.289 | 3.2926E-29 |

| | | | | | |
|---|---|---|---|---|---|
| LARS | 1.5269E-33 | 9.56652673 | 0.246 | 0.347 | 3.4736E-29 |
| ATP2A2 | 1.7573E-33 | 9.89809223 | 0.197 | 0.281 | 3.9978E-29 |
| GON4L | 3.0603E-33 | 0.56500908 | 0.172 | 0.251 | 6.9621E-29 |
| NDUFAF8 | 3.1959E-33 | 14.3541103 | 0.174 | 0.26 | 7.2706E-29 |
| MED10 | 3.4015E-33 | 12.9152696 | 0.183 | 0.269 | 7.7385E-29 |
| DDOST | 3.4824E-33 | 27.116486 | 0.276 | 0.388 | 7.9224E-29 |
| POLDIP2 | 4.1811E-33 | 3.18492601 | 0.174 | 0.26 | 9.5121E-29 |
| ZFC3H1 | 4.3875E-33 | 4.38603295 | 0.184 | 0.262 | 9.9816E-29 |
| USP34 | 8.6685E-33 | 2.05411063 | 0.221 | 0.304 | 1.9721E-28 |
| BZW1 | 1.1129E-32 | 23.4867184 | 0.358 | 0.47 | 2.5318E-28 |
| CDKN1A | 1.3664E-32 | 16.6185615 | 0.385 | 0.484 | 3.1086E-28 |
| NFIC | 1.8612E-32 | 6.08678805 | 0.198 | 0.287 | 4.2343E-28 |
| DNAJA2 | 1.8786E-32 | 13.7051862 | 0.239 | 0.339 | 4.2738E-28 |
| DNTTIP2 | 2.3159E-32 | 6.85904756 | 0.247 | 0.34 | 5.2688E-28 |
| FAM32A | 2.7546E-32 | 4.25102418 | 0.192 | 0.277 | 6.2666E-28 |
| AP2B1 | 3.9104E-32 | 0.37592397 | 0.181 | 0.258 | 8.8963E-28 |
| PDHA1 | 4.3076E-32 | 2.64507532 | 0.17 | 0.251 | 9.7999E-28 |
| SUPT6H | 5.7682E-32 | 6.37898316 | 0.263 | 0.356 | 1.3123E-27 |
| USP14 | 6.2633E-32 | 4.88056307 | 0.257 | 0.352 | 1.4249E-27 |
| DERL2 | 6.4954E-32 | 9.95987542 | 0.169 | 0.256 | 1.4777E-27 |
| ESF1 | 7.0648E-32 | 1.42152198 | 0.182 | 0.264 | 1.6072E-27 |
| ZNF593 | 7.3303E-32 | 10.3084897 | 0.194 | 0.284 | 1.6676E-27 |
| TOX4 | 8.2759E-32 | 8.97927248 | 0.212 | 0.299 | 1.8828E-27 |
| RHOBTB3 | 9.3543E-32 | 22.2747714 | 0.224 | 0.314 | 2.1281E-27 |
| STOML2 | 1.185E-31 | 17.750369 | 0.223 | 0.325 | 2.696E-27 |
| NSD1 | 1.5286E-31 | 0.51052478 | 0.2 | 0.28 | 3.4775E-27 |
| LSM1 | 2.0378E-31 | 2.62981168 | 0.17 | 0.251 | 4.636E-27 |
| ANKFY1 | 2.2455E-31 | 1.87888518 | 0.177 | 0.251 | 5.1085E-27 |
| TNPO1 | 2.6054E-31 | 15.5642217 | 0.253 | 0.35 | 5.9273E-27 |
| TNFRSF1A | 3.1997E-31 | 13.7655049 | 0.233 | 0.339 | 7.2792E-27 |
| MAT2A | 3.9281E-31 | 16.9504046 | 0.229 | 0.315 | 8.9365E-27 |
| SND1 | 5.3253E-31 | 41.9391073 | 0.325 | 0.433 | 1.2115E-26 |
| CACUL1 | 5.7369E-31 | 2.05264135 | 0.236 | 0.322 | 1.3051E-26 |
| ACTR1A | 6.7622E-31 | 3.18129573 | 0.228 | 0.315 | 1.5384E-26 |
| CDK12 | 8.5412E-31 | 2.99810717 | 0.217 | 0.301 | 1.9431E-26 |
| CHD8 | 9.0588E-31 | 1.35570727 | 0.193 | 0.268 | 2.0609E-26 |
| PPP2CA | 9.4911E-31 | 6.35859145 | 0.204 | 0.293 | 2.1592E-26 |
| RAB3GAP1 | 9.6328E-31 | 0.7658198 | 0.218 | 0.299 | 2.1915E-26 |
| COPS6 | 2.1624E-30 | 5.68258478 | 0.212 | 0.307 | 4.9194E-26 |
| SBDS | 2.297E-30 | 23.8319139 | 0.215 | 0.313 | 5.2257E-26 |
| HNRNPM | 2.7981E-30 | 13.7573247 | 0.36 | 0.47 | 6.3656E-26 |
| ATG12 | 3.2089E-30 | 7.03778994 | 0.197 | 0.278 | 7.3002E-26 |
| ECPAS | 3.6845E-30 | 1.64642061 | 0.205 | 0.285 | 8.3822E-26 |

| | | | | | |
|---|---|---|---|---|---|
| SRP54 | 3.8389E-30 | 0.58627163 | 0.212 | 0.294 | 8.7334E-26 |
| ARID5B | 4.1092E-30 | 13.4967086 | 0.221 | 0.31 | 9.3485E-26 |
| ABCF1 | 5.4223E-30 | 4.43320408 | 0.24 | 0.333 | 1.2336E-25 |
| MFF | 5.8692E-30 | 6.62899132 | 0.201 | 0.287 | 1.3352E-25 |
| PNN | 6.137E-30 | 21.4224514 | 0.271 | 0.367 | 1.3962E-25 |
| SENP6 | 6.3103E-30 | 3.17984578 | 0.236 | 0.323 | 1.4356E-25 |
| ZNF644 | 1.1368E-29 | 2.59693759 | 0.224 | 0.308 | 2.5862E-25 |
| IGF2BP1 | 1.4776E-29 | 5.25756579 | 0.175 | 0.252 | 3.3615E-25 |
| UBE2V2 | 1.5068E-29 | 4.55351278 | 0.179 | 0.261 | 3.428E-25 |
| IFIT1 | 1.57E-29 | 19.5039623 | 0.286 | 0.34 | 3.5718E-25 |
| USP16 | 2.0745E-29 | 4.04616748 | 0.183 | 0.263 | 4.7195E-25 |
| SLC25A39 | 2.4772E-29 | 11.9483449 | 0.199 | 0.291 | 5.6357E-25 |
| EMD | 4.4242E-29 | 25.6195638 | 0.199 | 0.29 | 1.0065E-24 |
| NUDC | 6.1458E-29 | 22.0925061 | 0.307 | 0.418 | 1.3982E-24 |
| KIF2A | 6.2137E-29 | 7.94706958 | 0.223 | 0.309 | 1.4136E-24 |
| ZNF428 | 6.5621E-29 | 12.0547208 | 0.182 | 0.262 | 1.4929E-24 |
| CDK4 | 7.48E-29 | 24.1499685 | 0.262 | 0.367 | 1.7017E-24 |
| SURF4 | 7.5459E-29 | 35.3765217 | 0.241 | 0.338 | 1.7167E-24 |
| SMARCA4 | 8.2321E-29 | 7.58255935 | 0.198 | 0.282 | 1.8728E-24 |
| LRRC59 | 1.2541E-28 | 77.060842 | 0.283 | 0.386 | 2.8531E-24 |
| UBA2 | 1.3394E-28 | 13.4849072 | 0.243 | 0.336 | 3.0471E-24 |
| TFAM | 1.4858E-28 | 10.2869474 | 0.183 | 0.268 | 3.3801E-24 |
| TIPRL | 1.763E-28 | 5.83421914 | 0.208 | 0.291 | 4.0108E-24 |
| RAB18 | 1.8665E-28 | 5.43209482 | 0.219 | 0.304 | 4.2464E-24 |
| TRIP11 | 1.9468E-28 | 1.21231742 | 0.177 | 0.252 | 4.429E-24 |
| CLIC4 | 2.5355E-28 | 36.2451489 | 0.305 | 0.396 | 5.7683E-24 |
| ZNF24 | 2.8339E-28 | 1.46433851 | 0.178 | 0.252 | 6.4472E-24 |
| PRMT1 | 3.4213E-28 | 28.0298184 | 0.247 | 0.35 | 7.7835E-24 |
| PPHLN1 | 3.8806E-28 | 1.23505304 | 0.181 | 0.255 | 8.8283E-24 |
| YTHDF2 | 4.2168E-28 | 7.68066462 | 0.227 | 0.316 | 9.5933E-24 |
| PHF3 | 4.8669E-28 | 4.79045129 | 0.206 | 0.283 | 1.1072E-23 |
| PQBP1 | 5.2992E-28 | 4.25477154 | 0.213 | 0.299 | 1.2056E-23 |
| EI24 | 5.8044E-28 | 5.65671483 | 0.182 | 0.263 | 1.3205E-23 |
| RRAGA | 8.7389E-28 | 5.724899 | 0.174 | 0.252 | 1.9881E-23 |
| DCUN1D5 | 9.8893E-28 | 9.86281517 | 0.201 | 0.289 | 2.2498E-23 |
| HNRNPH1 | 1.1195E-27 | 19.6945016 | 0.255 | 0.345 | 2.5469E-23 |
| TMX1 | 1.3569E-27 | 1.91350649 | 0.178 | 0.255 | 3.087E-23 |
| VAT1 | 1.6802E-27 | 2.14201296 | 0.292 | 0.374 | 3.8224E-23 |
| TOMM40 | 1.9293E-27 | 13.8530167 | 0.208 | 0.295 | 4.3891E-23 |
| RSL24D1 | 2.3359E-27 | 6.96640393 | 0.212 | 0.302 | 5.3142E-23 |
| EIF3J | 3.4903E-27 | 4.0143901 | 0.192 | 0.274 | 7.9405E-23 |
| EIF4H | 3.9245E-27 | 23.443097 | 0.323 | 0.426 | 8.9282E-23 |
| PSMD3 | 4.5022E-27 | 10.7109625 | 0.207 | 0.295 | 1.0243E-22 |

| | | | | | |
|---|---|---|---|---|---|
| THOC2 | 6.113E-27 | 3.10864286 | 0.214 | 0.296 | 1.3907E-22 |
| TRA2A | 6.4942E-27 | 1.89459662 | 0.2 | 0.279 | 1.4774E-22 |
| SMC1A | 6.7317E-27 | 32.9875785 | 0.215 | 0.295 | 1.5315E-22 |
| YWHAQ | 7.3647E-27 | 18.7941517 | 0.322 | 0.431 | 1.6755E-22 |
| OGA | 1.1142E-26 | 4.61345817 | 0.182 | 0.254 | 2.5347E-22 |
| KPNA2 | 1.2447E-26 | 81.8958048 | 0.196 | 0.274 | 2.8318E-22 |
| IK | 1.2864E-26 | 2.89143445 | 0.242 | 0.323 | 2.9266E-22 |
| BCLAF1 | 1.4417E-26 | 16.6292609 | 0.338 | 0.441 | 3.2799E-22 |
| POLR2A | 1.8546E-26 | 22.2126347 | 0.22 | 0.297 | 4.2191E-22 |
| COPS5 | 2.1498E-26 | 4.35430719 | 0.187 | 0.267 | 4.8908E-22 |
| COPG1 | 2.1608E-26 | 8.98468253 | 0.21 | 0.289 | 4.9159E-22 |
| NUCB1 | 2.2053E-26 | 16.5801347 | 0.298 | 0.386 | 5.0171E-22 |
| ZNF148 | 2.8358E-26 | 1.52298796 | 0.2 | 0.272 | 6.4513E-22 |
| DDX3X | 2.9414E-26 | 28.0562166 | 0.277 | 0.359 | 6.6916E-22 |
| CXCL1 | 2.9806E-26 | 18.0612323 | 0.392 | 0.452 | 6.781E-22 |
| SERINC3 | 3.5414E-26 | 2.89696781 | 0.191 | 0.267 | 8.0567E-22 |
| ALG3 | 3.8497E-26 | 6.88411249 | 0.184 | 0.267 | 8.7581E-22 |
| STRAP | 7.4473E-26 | 18.7009451 | 0.244 | 0.343 | 1.6943E-21 |
| RERE | 8.3485E-26 | 1.52573712 | 0.184 | 0.255 | 1.8993E-21 |
| JAGN1 | 8.7665E-26 | 6.14930764 | 0.18 | 0.258 | 1.9944E-21 |
| ACBD6 | 9.0072E-26 | 2.19858052 | 0.227 | 0.306 | 2.0491E-21 |
| CDC27 | 1.0786E-25 | 6.7775867 | 0.192 | 0.266 | 2.4539E-21 |
| PSMD6 | 1.8988E-25 | 8.35794281 | 0.223 | 0.307 | 4.3198E-21 |
| TMEM87A | 2.0582E-25 | 7.97196542 | 0.197 | 0.273 | 4.6824E-21 |
| GDI1 | 2.8243E-25 | 2.91126722 | 0.214 | 0.286 | 6.4253E-21 |
| ZRANB2 | 2.8265E-25 | 16.2150181 | 0.259 | 0.349 | 6.4303E-21 |
| UBR5 | 3.3452E-25 | 1.69044898 | 0.199 | 0.27 | 7.6103E-21 |
| PYGL | 3.7609E-25 | 4.62598347 | 0.22 | 0.288 | 8.556E-21 |
| SIVA1 | 4.1449E-25 | 14.4991396 | 0.204 | 0.288 | 9.4297E-21 |
| MORF4L2 | 5.8941E-25 | 30.6815532 | 0.287 | 0.395 | 1.3409E-20 |
| PTP4A1 | 6.3302E-25 | 5.11189778 | 0.226 | 0.301 | 1.4401E-20 |
| PWP1 | 6.3986E-25 | 7.95654173 | 0.195 | 0.271 | 1.4557E-20 |
| GIGYF2 | 6.5795E-25 | 2.18864236 | 0.217 | 0.292 | 1.4968E-20 |
| EFTUD2 | 7.0481E-25 | 3.94237498 | 0.232 | 0.312 | 1.6034E-20 |
| PHAX | 1.0786E-24 | 3.99019173 | 0.183 | 0.257 | 2.4539E-20 |
| NOTCH2 | 1.2814E-24 | 9.54100724 | 0.185 | 0.255 | 2.9151E-20 |
| SF3A1 | 1.3578E-24 | 19.3185568 | 0.221 | 0.298 | 3.0891E-20 |
| STAT2 | 1.7036E-24 | 2.49967337 | 0.204 | 0.274 | 3.8756E-20 |
| DCTN1 | 1.8127E-24 | 8.10845367 | 0.199 | 0.271 | 4.124E-20 |
| TTC17 | 1.8781E-24 | 0.77629458 | 0.239 | 0.312 | 4.2727E-20 |
| DHX36 | 1.931E-24 | 11.7541052 | 0.205 | 0.286 | 4.3929E-20 |
| NSD3 | 2.0905E-24 | 2.1934515 | 0.213 | 0.288 | 4.7558E-20 |
| DIAPH1 | 2.2598E-24 | 0.75966776 | 0.191 | 0.255 | 5.141E-20 |

| | | | | | |
|---|---|---|---|---|---|
| PRKAR2A | 2.334E-24 | 5.15951876 | 0.189 | 0.26 | 5.3099E-20 |
| SETD5 | 3.0163E-24 | 3.94231037 | 0.222 | 0.296 | 6.8622E-20 |
| SRPK2 | 3.064E-24 | 2.0753249 | 0.229 | 0.306 | 6.9706E-20 |
| HNRNPH3 | 3.249E-24 | 27.5173133 | 0.287 | 0.378 | 7.3915E-20 |
| SPART | 3.6162E-24 | 3.20556047 | 0.187 | 0.254 | 8.2269E-20 |
| YBX3 | 3.8657E-24 | 30.291642 | 0.183 | 0.266 | 8.7946E-20 |
| DDB1 | 4.3496E-24 | 15.1114736 | 0.237 | 0.318 | 9.8953E-20 |
| MAX | 5.4855E-24 | 0.28451554 | 0.271 | 0.347 | 1.248E-19 |
| WDR33 | 9.9424E-24 | 1.98853379 | 0.184 | 0.253 | 2.2619E-19 |
| SNRPA | 1.8015E-23 | 6.54559936 | 0.19 | 0.265 | 4.0984E-19 |
| HSD17B4 | 2.3097E-23 | 21.1132259 | 0.319 | 0.399 | 5.2545E-19 |
| ZCCHC7 | 2.3548E-23 | 0.8057187 | 0.209 | 0.278 | 5.3572E-19 |
| SMARCB1 | 3.1698E-23 | 9.16974781 | 0.199 | 0.28 | 7.2114E-19 |
| DHX9 | 3.2151E-23 | 11.0903584 | 0.259 | 0.344 | 7.3145E-19 |
| PPP2R2A | 3.885E-23 | 4.10001668 | 0.215 | 0.288 | 8.8384E-19 |
| RAB32 | 5.3593E-23 | 3.61486334 | 0.203 | 0.272 | 1.2192E-18 |
| LONP2 | 5.3702E-23 | 1.64637423 | 0.183 | 0.25 | 1.2217E-18 |
| COPS2 | 7.4069E-23 | 25.8851463 | 0.198 | 0.276 | 1.6851E-18 |
| CNOT2 | 8.9001E-23 | 0.8844067 | 0.198 | 0.27 | 2.0248E-18 |
| SUPT16H | 9.9482E-23 | 31.4187044 | 0.261 | 0.344 | 2.2632E-18 |
| AATF | 1.0698E-22 | 11.3179912 | 0.255 | 0.332 | 2.4339E-18 |
| DNAJC1 | 1.3385E-22 | 17.823341 | 0.179 | 0.252 | 3.045E-18 |
| DNAJC2 | 1.3507E-22 | 18.2754599 | 0.21 | 0.287 | 3.0727E-18 |
| SETD2 | 1.4625E-22 | 0.92248457 | 0.195 | 0.261 | 3.3271E-18 |
| HNRNPUL1 | 4.282E-22 | 4.34717003 | 0.189 | 0.258 | 9.7415E-18 |
| TRIO | 6.2822E-22 | 26.723786 | 0.306 | 0.389 | 1.4292E-17 |
| TCERG1 | 7.6809E-22 | 7.76806529 | 0.225 | 0.303 | 1.7474E-17 |
| ARIH2 | 8.4653E-22 | 1.98911939 | 0.194 | 0.264 | 1.9259E-17 |
| BUB3 | 1.0597E-21 | 6.07064084 | 0.207 | 0.28 | 2.4109E-17 |
| NAA50 | 1.0616E-21 | 3.67661056 | 0.219 | 0.294 | 2.4152E-17 |
| LARP1 | 1.1948E-21 | 2.99634212 | 0.189 | 0.255 | 2.7181E-17 |
| PABPC4 | 1.2289E-21 | 10.0758054 | 0.191 | 0.264 | 2.7958E-17 |
| SNRNP200 | 2.1582E-21 | 39.5017603 | 0.23 | 0.306 | 4.9098E-17 |
| SF3A3 | 2.2386E-21 | 12.2287576 | 0.223 | 0.299 | 5.0928E-17 |
| SYPL1 | 2.7535E-21 | 13.9288563 | 0.207 | 0.282 | 6.2642E-17 |
| NSFL1C | 3.5884E-21 | 7.24658228 | 0.221 | 0.298 | 8.1636E-17 |
| PSMA2 | 5.8195E-21 | 6.14018031 | 0.182 | 0.253 | 1.3239E-16 |
| ELAVL1 | 7.8183E-21 | 5.82367562 | 0.184 | 0.255 | 1.7787E-16 |
| CHD1 | 1.0031E-20 | 4.30681082 | 0.199 | 0.269 | 2.2822E-16 |
| CAPRIN1 | 1.876E-20 | 49.2198293 | 0.309 | 0.39 | 4.2678E-16 |
| SEC63 | 2.6225E-20 | 25.334587 | 0.254 | 0.341 | 5.9661E-16 |
| PUF60 | 3.0491E-20 | 14.359318 | 0.227 | 0.31 | 6.9366E-16 |
| DNMT1 | 3.5135E-20 | 5.51319951 | 0.231 | 0.304 | 7.9933E-16 |

| | | | | | |
|---|---|---|---|---|---|
| CLTB | 4.0626E-20 | 7.96237269 | 0.199 | 0.272 | 9.2425E-16 |
| EHMT1 | 5.1966E-20 | 2.50589043 | 0.224 | 0.292 | 1.1822E-15 |
| SAR1A | 5.6804E-20 | 12.5749204 | 0.189 | 0.262 | 1.2923E-15 |
| USP10 | 8.8694E-20 | 1.94724795 | 0.198 | 0.261 | 2.0178E-15 |
| SEL1L | 1.1444E-19 | 21.2876534 | 0.197 | 0.261 | 2.6036E-15 |
| NIFK | 1.3646E-19 | 17.5553855 | 0.226 | 0.303 | 3.1044E-15 |
| EIF4A3 | 1.485E-19 | 8.58366311 | 0.233 | 0.308 | 3.3783E-15 |
| PHF5A | 3.2614E-19 | 6.45241325 | 0.188 | 0.255 | 7.4196E-15 |
| TSG101 | 3.323E-19 | 3.69015589 | 0.219 | 0.286 | 7.5598E-15 |
| IFITM3 | 4.1532E-19 | 381.620384 | 0.577 | 0.591 | 9.4486E-15 |
| MANF | 4.8809E-19 | 88.7510336 | 0.254 | 0.34 | 1.1104E-14 |
| TM9SF2 | 5.4303E-19 | 1.73638154 | 0.303 | 0.377 | 1.2354E-14 |
| SHISA5 | 6.5612E-19 | 16.6238518 | 0.25 | 0.333 | 1.4927E-14 |
| DDX21 | 8.0023E-19 | 23.379101 | 0.346 | 0.434 | 1.8205E-14 |
| RNF216 | 8.1594E-19 | 1.61007416 | 0.204 | 0.266 | 1.8563E-14 |
| PTPN11 | 8.6187E-19 | 4.68190238 | 0.226 | 0.292 | 1.9607E-14 |
| HNRNPL | 1.3411E-18 | 3.77218831 | 0.195 | 0.26 | 3.051E-14 |
| MPG | 7.5553E-18 | 8.40023826 | 0.211 | 0.282 | 1.7188E-13 |
| SEC31A | 8.1714E-18 | 8.8974006 | 0.256 | 0.333 | 1.859E-13 |
| RNPS1 | 1.0505E-17 | 12.8917259 | 0.263 | 0.347 | 2.3899E-13 |
| NORAD | 1.6226E-17 | 11.6396494 | 0.229 | 0.296 | 3.6914E-13 |
| SNRPA1 | 2.1571E-17 | 7.37889247 | 0.188 | 0.253 | 4.9074E-13 |
| YTHDC1 | 2.5014E-17 | 7.25864656 | 0.202 | 0.267 | 5.6908E-13 |
| NOP53 | 3.0271E-17 | 2.01028302 | 0.219 | 0.284 | 6.8867E-13 |
| TGFBR2 | 3.2054E-17 | 19.394991 | 0.234 | 0.3 | 7.2922E-13 |
| PPP1CB | 3.4502E-17 | 14.4829654 | 0.201 | 0.262 | 7.8492E-13 |
| SRP72 | 3.6722E-17 | 5.68966022 | 0.27 | 0.347 | 8.3543E-13 |
| EIF2S1 | 3.8391E-17 | 5.47521043 | 0.271 | 0.344 | 8.7339E-13 |
| TMEM30A | 3.9499E-17 | 18.0033545 | 0.206 | 0.263 | 8.986E-13 |
| UBQLN1 | 1.0987E-16 | 13.5781148 | 0.233 | 0.294 | 2.4996E-12 |
| PPP1R10 | 4.0314E-16 | 44.3919419 | 0.197 | 0.255 | 9.1715E-12 |
| MAP4 | 4.642E-16 | 12.9659968 | 0.322 | 0.396 | 1.0561E-11 |
| IFRD1 | 4.9179E-16 | 6.68171643 | 0.202 | 0.259 | 1.1188E-11 |
| SAFB | 5.7864E-16 | 5.78895371 | 0.234 | 0.298 | 1.3164E-11 |
| DUT | 1.1296E-15 | 21.9822503 | 0.192 | 0.256 | 2.5699E-11 |
| TSC22D1 | 1.2065E-15 | 56.820486 | 0.224 | 0.287 | 2.7448E-11 |
| CLNS1A | 3.9139E-15 | 6.84419635 | 0.21 | 0.275 | 8.9041E-11 |
| NUTM2A-AS1 | 4.298E-15 | 2.15280847 | 0.209 | 0.263 | 9.7779E-11 |
| ANAPC5 | 6.1249E-15 | 4.15200776 | 0.198 | 0.258 | 1.3934E-10 |
| NASP | 6.3622E-15 | 16.7095366 | 0.208 | 0.269 | 1.4474E-10 |
| TFG | 6.9536E-15 | 16.7082624 | 0.217 | 0.289 | 1.5819E-10 |
| SUPT5H | 7.5312E-15 | 4.99123468 | 0.199 | 0.253 | 1.7134E-10 |
| RBBP4 | 1.1579E-14 | 15.6563852 | 0.276 | 0.35 | 2.6343E-10 |

| | | | | | |
|---|---|---|---|---|---|
| PCMT1 | 1.4542E-14 | 2.53758679 | 0.245 | 0.306 | 3.3084E-10 |
| TMED3 | 1.6299E-14 | 6.6507782 | 0.19 | 0.251 | 3.7081E-10 |
| USP22 | 1.6868E-14 | 13.9140199 | 0.221 | 0.28 | 3.8376E-10 |
| SSR1 | 1.8099E-14 | 17.5399913 | 0.234 | 0.303 | 4.1174E-10 |
| GTPBP4 | 2.8381E-14 | 30.9203074 | 0.24 | 0.311 | 6.4566E-10 |
| TMF1 | 2.9873E-14 | 18.5535224 | 0.198 | 0.255 | 6.7961E-10 |
| TIAL1 | 3.3397E-14 | 4.27434078 | 0.194 | 0.251 | 7.5978E-10 |
| SMARCC1 | 3.3491E-14 | 8.10834547 | 0.227 | 0.292 | 7.6192E-10 |
| AK2 | 3.3586E-14 | 10.052663 | 0.25 | 0.318 | 7.6408E-10 |
| SLC2A3 | 4.1288E-14 | 6.38505384 | 0.238 | 0.29 | 9.3929E-10 |
| BBX | 4.7009E-14 | 7.12229251 | 0.196 | 0.253 | 1.0695E-09 |
| ARCN1 | 5.3385E-14 | 17.9131564 | 0.251 | 0.31 | 1.2145E-09 |
| PHF14 | 5.7393E-14 | 3.2104958 | 0.2 | 0.256 | 1.3057E-09 |
| UBAP2L | 8.7329E-14 | 5.58567907 | 0.201 | 0.254 | 1.9867E-09 |
| MRTO4 | 2.5268E-13 | 19.8094861 | 0.193 | 0.254 | 5.7484E-09 |
| EIF4A2 | 2.7769E-13 | 34.4614771 | 0.277 | 0.352 | 6.3175E-09 |
| PTBP1 | 3.499E-13 | 6.04581489 | 0.234 | 0.294 | 7.9602E-09 |
| USP47 | 3.6334E-13 | 1.85743817 | 0.205 | 0.258 | 8.2661E-09 |
| CHORDC1 | 3.6358E-13 | 11.0741871 | 0.202 | 0.256 | 8.2714E-09 |
| CFDP1 | 4.4183E-13 | 4.05982736 | 0.214 | 0.272 | 1.0052E-08 |
| SEC61A1 | 7.722E-13 | 23.7782967 | 0.239 | 0.307 | 1.7568E-08 |
| ANXA1 | 1.3976E-12 | 260.433999 | 0.552 | 0.616 | 3.1795E-08 |
| SRSF1 | 1.4501E-12 | 9.97514258 | 0.204 | 0.262 | 3.2991E-08 |
| GSPT1 | 1.5635E-12 | 13.7922634 | 0.228 | 0.287 | 3.5569E-08 |
| TRIM44 | 4.8269E-12 | 13.0827549 | 0.231 | 0.286 | 1.0981E-07 |
| IFITM1 | 5.5872E-12 | 247.379585 | 0.453 | 0.372 | 1.2711E-07 |
| U2AF2 | 6.7635E-12 | 11.8388449 | 0.215 | 0.269 | 1.5387E-07 |
| FKBP4 | 1.0773E-11 | 50.998383 | 0.28 | 0.343 | 2.4509E-07 |
| IMPDH2 | 1.9691E-11 | 15.6454263 | 0.222 | 0.287 | 4.4796E-07 |
| ILF3 | 2.0421E-11 | 21.6939339 | 0.29 | 0.367 | 4.6458E-07 |
| SMU1 | 2.0984E-11 | 3.93898106 | 0.205 | 0.256 | 4.7739E-07 |
| BCCIP | 3.4531E-11 | 6.89580532 | 0.219 | 0.277 | 7.8558E-07 |
| TCP1 | 4.4985E-11 | 39.2407284 | 0.269 | 0.337 | 1.0234E-06 |
| IFITM2 | 4.9415E-11 | 71.4409351 | 0.51 | 0.505 | 1.1242E-06 |
| SPARC | 4.9799E-11 | 484.903829 | 0.303 | 0.291 | 1.1329E-06 |
| METTL9 | 6.8804E-11 | 9.69402247 | 0.258 | 0.316 | 1.5653E-06 |
| HIST1H4C | 8.2532E-11 | 2.20184361 | 0.209 | 0.261 | 1.8776E-06 |
| AHSA1 | 1.3333E-10 | 9.83373443 | 0.208 | 0.264 | 3.0333E-06 |
| NAP1L4 | 1.5617E-10 | 5.56500964 | 0.207 | 0.255 | 3.5528E-06 |
| IGF2BP2 | 2.5666E-10 | 5.24455637 | 0.252 | 0.301 | 5.8389E-06 |
| PDIA4 | 2.5775E-10 | 83.750489 | 0.271 | 0.337 | 5.8638E-06 |
| PITPNB | 3.1657E-10 | 13.4813504 | 0.228 | 0.278 | 7.2019E-06 |
| ASCC3 | 7.2724E-10 | 14.4822146 | 0.213 | 0.264 | 1.6545E-05 |

| | | | | | |
|---|---|---|---|---|---|
| PRKCSH | 9.5353E-10 | 11.0042248 | 0.221 | 0.278 | 2.1693E-05 |
| LMNA | 9.5912E-10 | 10.5344425 | 0.331 | 0.393 | 2.182E-05 |
| ABCE1 | 9.7338E-10 | 10.6052395 | 0.218 | 0.275 | 2.2144E-05 |
| SRRT | 2.4421E-09 | 26.1927789 | 0.218 | 0.264 | 5.5558E-05 |
| PRDX2 | 5.3447E-09 | 28.1562735 | 0.201 | 0.256 | 0.00012159 |
| ATXN10 | 5.5905E-09 | 22.3808689 | 0.27 | 0.337 | 0.00012718 |
| SMIM3 | 6.3441E-09 | 27.9350084 | 0.234 | 0.287 | 0.00014433 |
| LUC7L2 | 9.6169E-09 | 5.20721705 | 0.22 | 0.268 | 0.00021879 |
| CLINT1 | 3.2257E-07 | 9.3591814 | 0.272 | 0.255 | 0.00733855 |
| NOP58 | 3.7152E-07 | 30.8841396 | 0.232 | 0.282 | 0.00845216 |
| PTPRG | 5.5731E-07 | 42.3718477 | 0.232 | 0.277 | 0.01267877 |
| C9orf78 | 5.8515E-07 | 2.35699203 | 0.228 | 0.269 | 0.01331205 |
| AHNAK | 6.1476E-07 | 156.212857 | 0.34 | 0.314 | 0.01398571 |
| NOP56 | 1.2011E-06 | 37.229662 | 0.212 | 0.256 | 0.02732615 |
| TNFRSF12A | 2.1487E-06 | 107.507034 | 0.269 | 0.264 | 0.04888385 |
