## Supplemental Table 2 for "Myeloid PINK1 represses mtDNA release and immune signaling that impacts neuronal pathology in patient-derived idiopathic PD models"

**Supplementary Table 2**

| Gene |
|---|
| CXCL6 |
| CXCL8 |
| CXCL1 |
| HAPLN1 |
| RGS18 |
| CXCL3 |
| RSAD2 |
| IFI44L |
| CXCL5 |
| CMPK2 |
| IL6 |
| AB13BP |
| CXCL10 |
| HBG1 |
| TNFSF18 |
| VNN2 |
| BST2 |
| HBG2 |
| GPR1 |
| CT88 |
| MX1 |
| ANO3 |
| NR4A3 |
| WISP1 |
| PDE4B |
| OAS1 |
| OLR1 |
| TNFSF15 |
| ANKRD34B |
| CXCL2 |
| OAS2 |
| SLITRK2 |
| TNFAIP3 |
| IL1B |
| ADAMT59 |
| C6ORF4 |
| ITIH5 |
| VNN3 |
| CCL13 |
| PDPN |

| |
|---|
| MX2 |
| IL20RB |
| MUSK |
| HERC6 |
| TNFSF13B |
| VEPH1 |
| C3 |
| ANGPTL4 |
| TNXB |
| C11ORF57 |
| CCL7 |
| ADAMT58 |
| SPTLC3 |
| ACKR4 |
| CHI3L1 |
| STAT1 |
| SP100 |
| IFI204 |
| ZBP1 |
| IFI44 |
| ISG15 |
| OASL2 |
| OASL1 |
| USP18 |
| IRF7 |
| NLRC5 |
| XAF1 |
| RNF213 |
| IFI27 |
| IFI2712A |
| RTP4 |
| IFIT2 |
| IFIT3 |
| IFIT3B |
| IFIT1 |
| B2M |
| TAP1 |
| H2-K1 |
| H2-D1 |
| H2-Q7 |
| SLAMF9 |
| CD84 |
| LY9 |

| |
|---|
| CTSZ |
| CTS7 |
| CYBB |
| CD52 |
| CD72 |
| GPNMB |
| CLEC7A |
| CD9 |
| ITGAM |
| ITGAX |
| LYZ2 |
| LILRB4A |
| LGALS3 |
| PLAU |
| ST14 |
| MMP12 |
| CCL6 |
| CD68 |
| CXCL16 |
| LGALS3BP |
| GPR84 |
| TSPO |
| APOD |
| CD86 |
| TREM2 |
